# Mammal placental phenotypes are predictable from microRNA repertoires

**DOI:** 10.1101/2025.06.27.660101

**Authors:** Jonathan Fenn, Jessica C. Edge, Vladimir Ovchinnikov, Olga Amelkina, Meléa Sinclair, Tiago H. C. De Bem, Alessandra Bridi, Irene Malo-Estepa, Angela Gonella-Diaza, Felipe Perecin, Juliano C. da Silveira, Guilherme Pugliesi, Flávio Vieira Meirelles, Zenab Butt, Haidee Tinning, Giulio Formenti, Erich Jarvis, Bastian Fromm, James O. McInerney, Niamh Forde, Mary J. O’Connell

## Abstract

Similar placental morphologies evolved multiple times independently in the history of mammal evolution^1,2^. Yet the genetic architecture that repeatedly guides distinct mammal lineages towards similar complex placental phenotypes has remained elusive. MicroRNAs (miRNAs), despite their diversity in mammals^3–7^ and known roles as developmental regulators^8–10^, remain under-examined as drivers of morphological innovation. We identified presence-absence patterns for 429 miRNA gene families across 398 mammalian genomes and discovered that placental phenotype is highly predictable from genomic miRNA repertoires (classification accuracy 74.5-95.8%). We identified 42 miRNA gene families significantly associated with placentation type, whose gene targets are enriched for developmental processes. Notably, convergent placental morphologies consistently involve identical miRNA families, revealing that evolution of this trait is constrained to predictable genetic pathways. We demonstrate that MIR-11986, uniquely associated with cotyledonary placentation, has tissue-specific expression in key reproductive tissues. MiRNA-mediated regulation therefore constrains placental morphological diversification into reproducible programs, offering insights into how genetic architecture shapes the predictability of convergent evolution. This striking pattern reveals a fundamental principle of evolution: that the miRNA regulatory networks available to control and guide complex placental morphological innovation are constrained and predictable.

## Introduction

Extant mammals share key defining traits, including implantation of the embryo and development of the placenta throughout pregnancy. Amongst eutherian mammals these traits manifest high levels of inter-species diversity and convergence, with well documented differences in key developmental checkpoints, degree of implantation into the maternal endometrium, development of the placental shape and venous system, and the degree of invasiveness of the maternal blood supply at the maternal-fetal interface^2,11^. These innovations, in particular in placental morphology, are cited as a potential mechanism for increasing reproductive isolation and fuelling speciation^2^, making them a vital consideration when investigating diversification, adaptation and speciation within Eutheria. Here we address two fundamental questions in evolutionary biology: what molecular mechanisms drive the evolution of eutherian mammal placental diversity? And, when similar complex placental morphologies emerge independently, have they evolved by shared or distinct solutions?

Placental divergence has been studied intensely in terms of protein coding contributors, for example: gene families disproportionately expressed in placenta^12^; lineage-specific variation in gene expansions in growth hormone and prolactin gene family and pregnancy specific glycoprotein gene family size^13^; lineage-specific gene families such as IFNT in ruminants (which also displays variation across ruminants)^14,15^; and the independent integration of endogenous retro-viral elements such as synctins^16,17^ in various lineages and their subsequent co-option to similar functions. MicroRNAs (miRNAs) are underexplored in the evolution of phenotypic diversity, yet these regulatory molecules have the capacity to affect thousands of genes in dynamic, flexible and tissue-specific ways^18–20^, coordinating paths to placental phenotypic variation over short evolutionary timescales^4,21–25^.

The largest known miRNA gene family expansion in animals is in the eutherian clade^26^. This expansion in miRNA repertoire has been linked to the evolution of diversity in mammal placental and implantation phenotypes, embryonic diversity, and other unique aspects of mammal reproduction and development^6,27,28^. Appropriate expression of many of these eutherian-unique miRNAs is required for successful embryonic implantation^29–31^ and subsequent placental development^32,33^. There is a potential distinction between miRNAs that originated at or within the eutherian clade that promote placentation and embryonic invasion, and miRNAs that pre-date the emergence of mammals that play a more inhibitory role^4,34–37^. The essential role of eutherian-unique and highly conserved miRNAs in the origin of the placenta has previously been shown^4^. For example, MIR-378^29^ facilitates trophoblast invasion, the physiological process during early pregnancy where embryonic cells attach to and penetrate the maternal uterine lining (the endometrium) to establish the placenta, through targeting of the Nodal pathway. Conversely, miRNA families that pre-date the origin of mammals such as MIR-155^38^ (conserved across vertebrates^39^), and MIR-210^40^ and MIR-29^41^ (conserved across Bilateria^42^), have been shown to inhibit trophoblast invasion, and can be pathogenic if overexpressed^40,41^. These associations suggest that characteristic elements of eutherian reproduction may be connected to the rapid expansion and diversification of miRNA repertoires in eutherian genomes since their divergence from marsupial (metatherian) mammals. Today, the rapid growth of sequenced high-quality genomes and transcriptomes for mammals, and particularly those part of the Vertebrate Genomes Project Phase 1^43^, can be used to identify the miRNA gene families that are indispensable for eutherian reproductive diversity, and those that underpin phenotypic diversity more broadly^44–46^ (Supplementary Fig S1).

Whilst broad associations between mammal placental phenotypes and life history have been drawn, *e.g.* larger longer-lived species with smaller litters are associated with diffuse placentas and epitheliochorial membranes (superficial, non-invasive placentas), while smaller shorter-lived species with larger litters are associated with discoid shaped placentas (like in mouse), labyrinthine venous systems and hemochorial membranes^47^, these patterns do not universally hold true across Eutheria. Convergent placental phenotypes are observed in species inhabiting quite different ecological niches - for example villous placental venous systems have evolved in Xenarthra (sloths and armadillos), Perissodactyla (horses, rhinos and tapirs), Artiodactyla (even-toed ungulates), and primates. There is, however, generally strong phylogenetic structure to their distribution, with some traits being highly clade-specific (*e.g.* the cotyledonary placental phenotype in Pecoran ruminants).

We hypothesise that despite the remarkable diversity of placental phenotypes across Eutheria, convergent evolution of specific placental traits is underpinned by consistent molecular mechanisms, specifically involvement of the same miRNA gene families (*e.g.* independent loss of the same miRNA family in different lineages). If true, it would also imply that placental morphology may be predictable by taking a census of the miRNA repertoire.

Using complete-genome sequence data from 398 mammal species, of which 124 also have placental phenotype data, we investigated whether the miRNA profiles specifically associated with convergent placental phenotypes evolved independently, or if similar regulatory mechanisms have repeatedly emerged to control similar morphological outcomes. We find that miRNA repertoire is highly predictive of placental phenotype, and we uncover two broad patterns of miRNA-linked phenotypic innovation. Convergent traits are commonly associated with loss of specific miRNA families, whilst non-convergent, single-origin traits are associated with the acquisition of completely new miRNA genes, as is most apparent in species with cotyledonary placentas. This demonstrates that lineage-specific phenotypes evolve via novel gene acquisition, mimicking natural gene insertion experiments. Furthermore, *in vitro* and *in vivo* analysis of the function of miRNA gene family MIR-11968, identified in our study as having very strong and clade-specific associations with the cotyledonary placental phenotype, revealed tissue- and species-specific expression patterns in bovine endometrial and placental tissue. This highlights the impact expression of a single miRNA can have on crucial stages of mammal development. This work provides insight into the molecular underpinnings of mammal pregnancy success, identifies regulatory drivers of placental diversification, and highlights miRNA families as candidates for investigating reproductive pathologies in clinical and agricultural settings.

## RESULTS

Across 398 mammal genomes, we used MirMachine^48^ and BLAST^49^ to identify 429 miRNA gene families present only in mammals (as denoted by MirGeneDB)^26^: 301 of which were exclusive to Eutheria (*i.e*. not in monotremes or marsupials), whereas 107 were unique to marsupials and monotremes. After quality filtering for genome completeness, 57 miRNA gene families were found in >90% of eutherian genomes (Supplementary Table S5). Hierarchical clustering on the miRNA presence-absence matrix, based on Jaccard dissimilarity, both at the species and miRNA gene family level, showed that Eutherian species fell into 7 well-supported clusters, defining Muroid Rodents, Simian Primates, all other Euarchontiglires, ruminant Artiodactyla, all other Artiodactyla, Carnivora and Chiroptera (notably this group is mainly characterised by a relatively depleted miRNA repertoire, and as such incorporates some other species). miRNA families also formed 6 stable clusters (Fig 1). Concordance between the dendrogram derived from hierarchical clustering and the consensus species phylogeny was very low (normalised Jaccard Robinson-Foulds distance = 0.782) indicating that across Eutheria, miRNA repertoires generally have low agreement with phylogenies derived from standard molecular phylogenetics. The clade-specific clusters in rodents and simian primates represent distinct exceptions to this pattern, as each clade contains well-defined clusters of miRNA gene families uniquely associated with their respective taxonomic groups. Dollo parsimony-based ancestral reconstruction placed 114 miRNA gene families in the eutherian last common ancestor (LCA), with only 8 inherited from the LCA of all mammals (including monotremes) and 13 from the therian LCA, indicating a burst of eutherian-specific miRNA diversification. One miRNA family present in the mammalian LCA, MIR-1327, was lost in the Eutherian LCA. The miRNA full repertoire of these ancestral nodes is provided in Supplementary Table S6.

**Figure 1.**
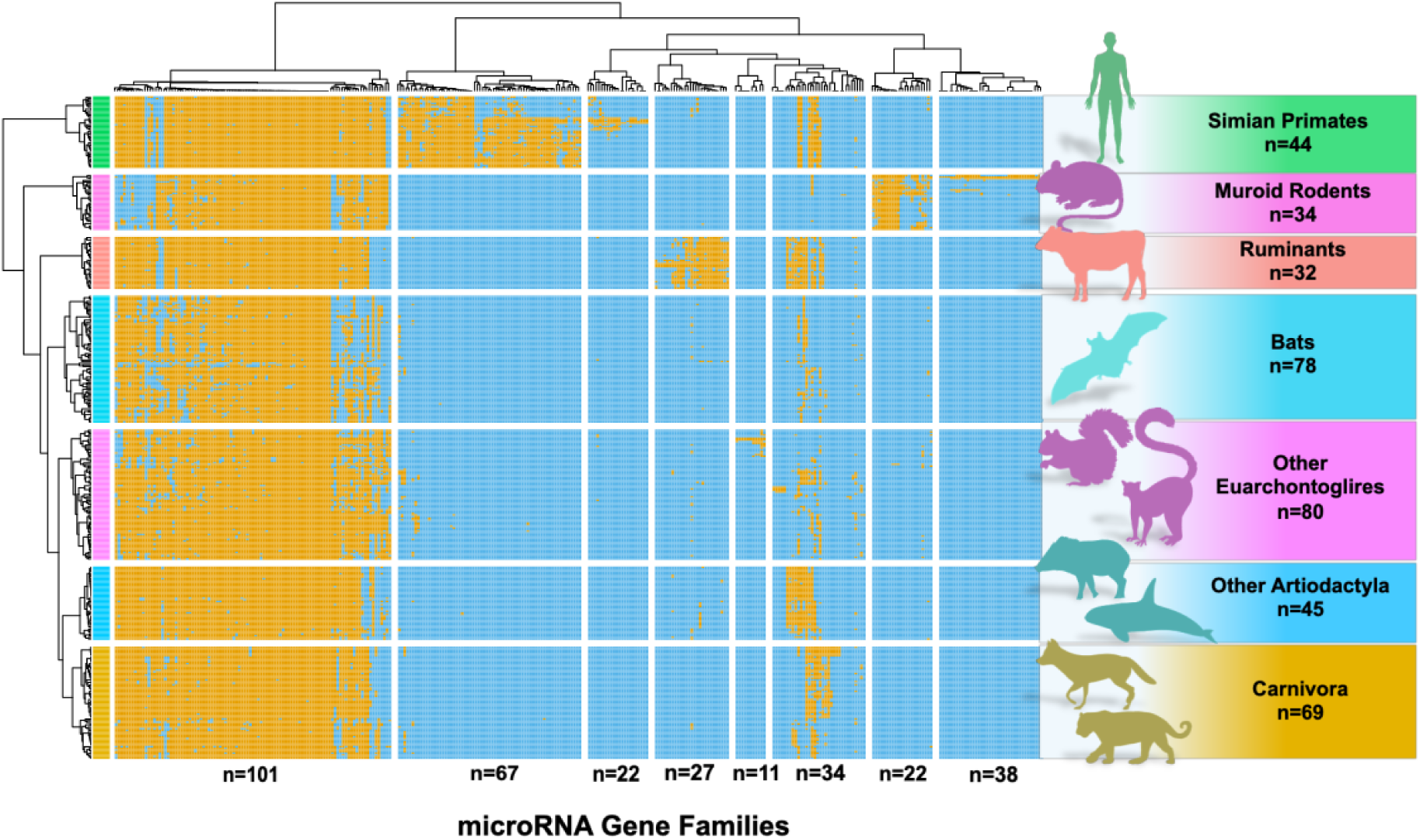
Clustered miRNA Gene Family presence/absence matrix. Matrix showing presence (*orange*) vs absence (*blue*) of 529 mammal-specific miRNA gene families across 398 mammal species, as detected by BLAST and MirMachine. Species are clustered by miRNA repertoire (*n=7*, horizontal breaks), while miRNA families are clustered by co-occurrence patterns in species (*n=8* clusters, vertical breaks), with associated clustering dendrograms. The taxonomic groupings of the species-level clusters are listed on the right-hand side of the plot.

We identify 10 miRNA gene families to be synapomorphic across Eutheria, allowing for loss in one species: Mir28, Mir324, Mir331, Mir378, Mir423 (as previously reported in^4^), along with Mir136, Mir154, Mir 320, Mir 362 and Mir376. The remaining 8 miRNA gene families previously reported to be synapomorphic^4^, while widely distributed, show discrete losses across Eutheria when examined across this comprehensive sampling (Fig. 1), *e.g.* MIR-505 is convergently lost from 6 bat species, 5 rodent species and 2 members of Eulipotyphla. Despite these sporadic losses, prevalence of these 10 eutherian unique miRNA families was still very high, ranging from 96-99.5%, indicating significant roles in eutherian evolution.

### Uniquely Derived phenotypes are associated with miRNA gain

Most associations identified between miRNAs and placental phenotype, and that are supported by both random forest and Evolink analyses, appear in miRNAs which have been widely lost, but preferentially retained in specific phenotype groups. In some instances, these loss associations can be linked to phenotype convergence. For example, MIR-872, which emerged in the therian ancestor, is widely lost across eutheria, with two such losses occurring at ancestral nodes to both zonary placenta shape clades, *i.e.* the *Carnivora* and *Afrotheria*. MIR-2483 arose in the eutherian ancestor and has been widely lost, but at much higher rates in taxa retaining the ancestral discoid/bidiscoid placental shape.

A group of 15 miRNA families arose in the ancestor of ruminants, and are thus strongly associated with the cotyledonary placenta shape specific to the pecoran ruminants. Two of these families, MIR-11968 and MIR-2331 have complete 1-to-1 association with the cotyledonary phenotype, being present in every species in which this phenotype is recorded, and not present alongside any other recorded phenotype, following a loss leading to the Javan lesser chevrotain *(Tragulus javanicus)*, which does not have this phenotype. MIR-2331 was recorded absent from the Dama gazelle, *Nanger dama*, for which phenotype data is not available, but which is a member of Pecora, and thus may be assumed to have a cotyledonary placenta. Two other families, MIR-3431 and MIR-10182 were not lost in *T. javanicus*, and so do not have this complete mutual association. The other 11 families in this group show some level of loss across pecoran ruminants - in some cases just a single loss. MIR-3956 and MIR-3432 arose earlier in the Artiodactyla LCA and thus do not have 1-to-1 associations with the cotyledonary phenotype (Fig. 2).

**Figure 2.**
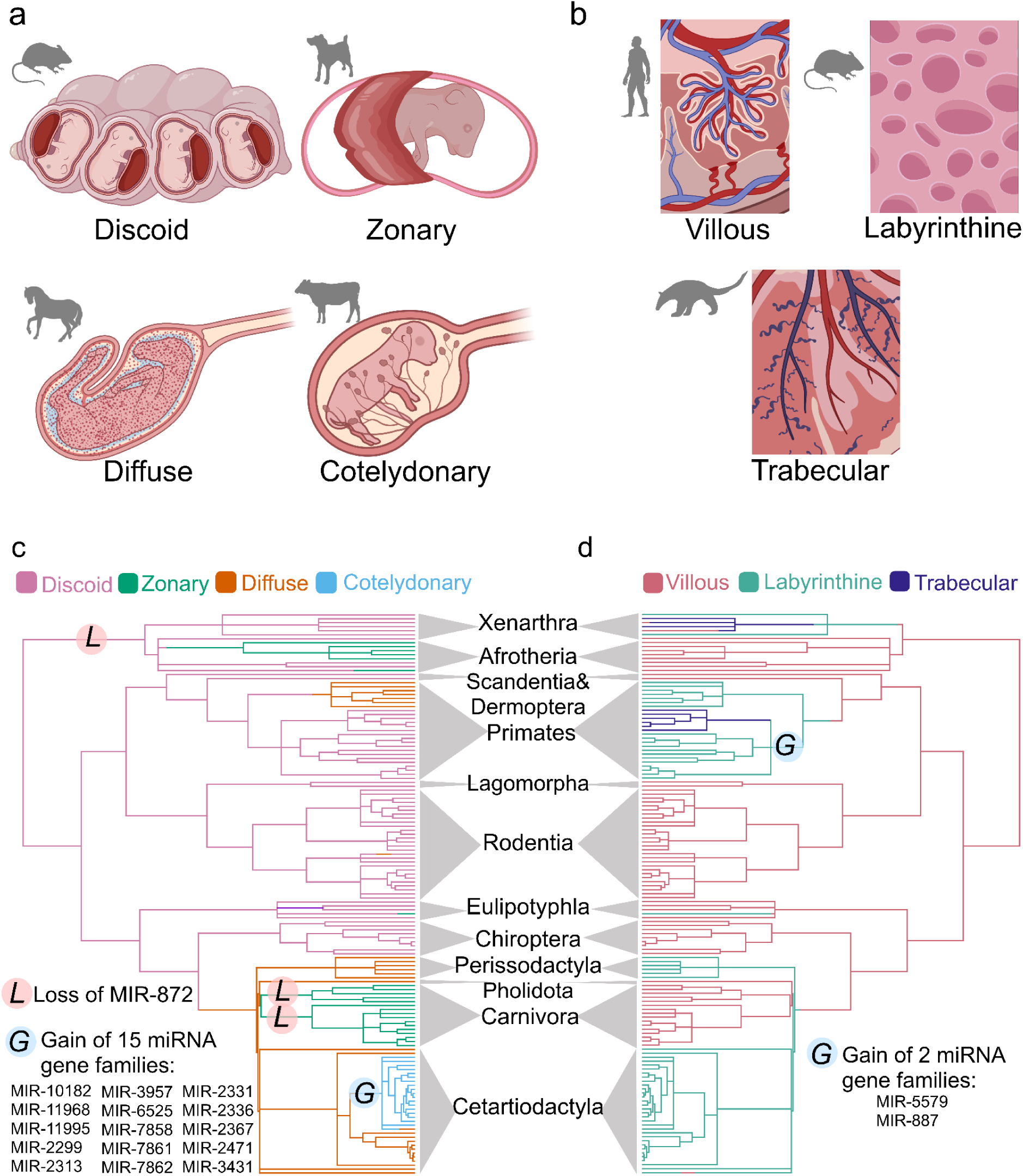
Diversity of placental shapes and venous patterns across Eutheria and phenotype associated miRNA families. a) Variation in placental shape across eutheria, *i.e.* discoid, zonary, diffuse and cotelydonary, alongside silhouettes from phylopic (https://www.phylopic.org/) depicting examples of species with those placental shapes, mouse, dog, horse and bovine respectively. b) Variation in placental venous patterns across eutheria, *i.e.* villous, labyrinthine and trabecular, alongside silhouettes from phylopic (https://www.phylopic.org/) depicting examples of species with those venous patterns, *i.e.* human, mouse, and anteater respectively. Both c) and d) display an identical eutherian species topology in mirror image with orders shown for orientation purposes. c) Shows the reconstruction of placental shape, and d) shows the reconstruction of venous pattern. Positions on the trees where there is strong evidence for miRNA-phenotype associations are depicted as “L” for losses of miRNAs gene families, and “G” for gains of miRNA gene families. The miRNA gene families implicated are listed in each case by their MirGeneDB identifiers. Images were created in https://BioRender.com.

Two phenotype-associated miRNA families (MIR-5579, MIR-887) show no secondary losses and are exclusively linked to trabecular venous patterns, both being ancestral to the primary trabecular group, the New World monkeys *(Platyrrhini),* having emerged in the ancestor to Simian primates. A summary of the strongest associations of miRNA presence/absence are shown in Fig. 2, and the full record of miRNA-phenotype associations is provided in Supplementary Table S7.

In contrast to the unique and unreversed property of MIR-11968, the majority of non-ancestral phenotypes have some evidence of convergent evolution. The miRNA-phenotype associations across these convergent and non-convergent phenotypes reveal two quite different evolutionary patterns. Convergent traits show repeated miRNA loss events, suggesting that certain developmental constraints permit specific regulatory losses to facilitate the evolution of particular phenotypes. Non-convergent traits, in contrast, are associated with miRNA gains, as seen with cotyledonary placenta and the simultaneous acquisition of multiple associated miRNA families in ruminants, including MIR-11968 and MIR-2331. This suggests that novel regulatory gains can enable unique morphological innovations but depend on rare, lineage-specific evolutionary opportunities. The balance between convergence driven by loss, and innovation driven by gain, are likely to reflect the developmental complexity of each trait and the genomic flexibility available for its evolution.

### MIR-11968 Expression Analysis

To investigate the functional significance of the association between MIR-11968 and the cotyledonary phenotype, we examined MIR-11968 expression across bovine endometrial tissues and cell types, providing the first insights into how clade-specific miRNAs may drive specialised placental phenotypes. miRBase describes three paralogs of this gene family (miRBase family ID = MIR-11986) whereas only two are described in MirGeneDB. All three paralogs were tested, and as such, the paralog not present in MirGeneDB will be referred to by its miRBase identifier - bta-miR-11986^26,42,50^. As expected (given the clade specific nature of MIR-11968), we found no detectable expression of btamiR-11986, Mir-11968-P1 or Mir-11968-P2 in human Ishikawa (endometrial epithelial cell line) cells. *In vitro* analysis of expression levels in bovine endometrial luminal epithelial, stromal and organoid cells showed bta-miR-11986 and Mir-11968-P2 are not detected. However, Mir-11968-P1 was expressed in bovine luminal epithelial cells and organoids, but not in stromal cells. Bta-miR-11986 was significantly (p<0.0001, one-way ANOVA with Šidák’s test for multiple comparisons) more highly expressed in bovine endometrial organoids than luminal epithelial cells. Treatment with the pregnancy recognition signal roIFNT did not significantly alter expression of bta-Mir-11968-P1 in organoids or luminal epithelial cells compared to their respective controls. Bta-Mir-11968-P1 was more highly expressed (p<0.0001, one-way ANOVA with Šidák’s test for multiple comparisons) in organoids treated with roIFNT than luminal epithelial cells treated with roIFNT (Fig. 3a). To determine whether MIR-11968 shows tissue-specific expression patterns relevant to cotyledonary placentation, we next examined its expression in caruncular versus intercaruncular endometrial tissues, which represent the specialized versus non-specialized regions of the bovine endometrium. In bovine species, the caruncular tissue forms the primary site of fetal-maternal interaction, where caruncles interdigitate with cotyledons of the placenta, whereas the surrounding intercaruncular regions contain glandular tissue. Using caruncular and intercaruncular endometrial samples collected *in vivo*, we find expression of bta-miR-11986, bta-Mir-11968-P1 and bta-Mir-11968-P2 in both tissues. The expression of bta-miR-11986 (p=0.0002), bta-mir-11968-P1 (p=0.0004) and bta-Mir-11968-P2 was significantly (p<0.0001) higher in caruncular tissue compared to intercaruncular (Fig. 3c-e). Finally, embryo developmental competency had no significant effect (p>0.05) on bta-miR-11986, bta-Mir-11968-P1 or bta-Mir-11968-P2 expression *in vivo*. Together, these findings illustrate that MIR-11968, a miRNA with strong evolutionary association to a derived placental phenotype, has tissue- and species-specific expression patterns supporting its role in the development and/or maintenance of the highly derived cotyledonary placental structure of pecoran ruminants.

**Figure 3.**
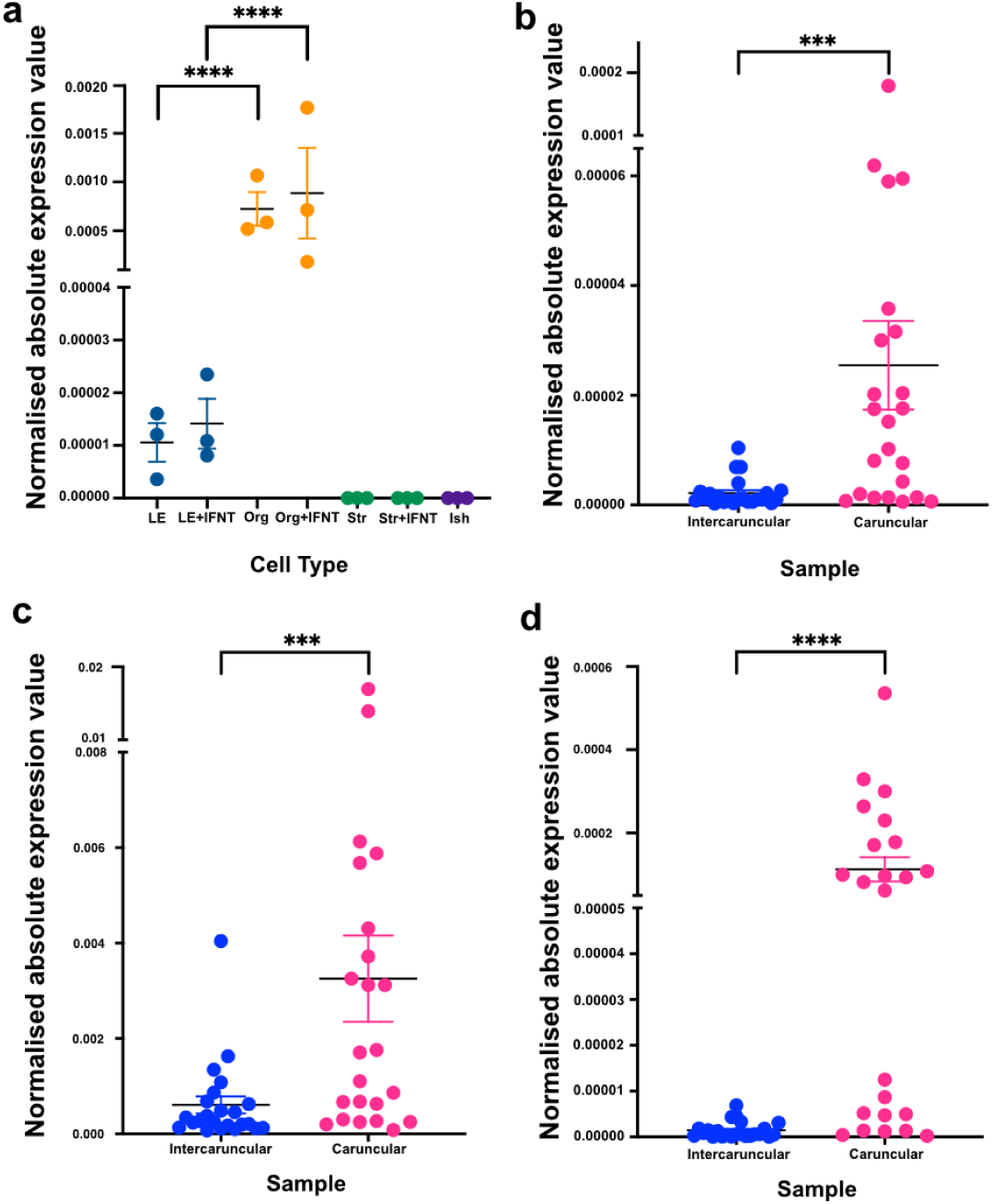
MIR-11968, showing complete evolutionary association with cotyledonary placentas, shows species- and tissue-specific expression in bovine endometrium. Function of the cotyledonary placenta-associated MIR-11968 paralogues were assessed through *in vitro* and *in vivo* analysis. a) Mir-11968-P1 expression in different *in vitro* models of the endometrium. Expression of Mir-11968-P1 was measured by qRT-PCR in primary bovine luminal epithelial cells (LE=Luminal Epithelium, Org=Endometrial Organoids, Str=Stromal Cells, Ish=Ishikawa Cells, IFNT=Interferon Tau), (*n=3*), bovine glandular epithelial organoids (*n=3*) and bovine stromal cells (*n=3*) and in each cell type treated with 1μg/μl recombinant ovine IFNT for 24 hours (*n=3*). Mir-11968-P1 expression was also measured in human endometrial epithelial (Ishikawa) cells (*n=3*) as a negative control. Significant differences where p<0.0001 are denoted by **** (b-d) bta-miR-11986, Mir-11968-P1 and Mir-11968-P2 expression in bovine caruncular or intercaruncular endometrial tissue. Expression of b) miR-11986, c) Mir-11968-P1 and d) Mir-11968-P2, were measured by qRT-PCR in bovine caruncular (*n=25*) and intercaruncular (*n=25*) endometrial samples on Day 16 of pregnancy. Summary lines in all boxplots represent mean values, error bars represent standard error of the mean. Significant differences in expression where p<0.001 are denoted by ***, and where p<0.0001 are denoted by ****.

### MiRNA gene family repertoire accurately predicts phenotype

Using a continuous-time reversible Markov model of placental phenotype evolution, we reconstructed the phenotype of the eutherian LCA inferring it to consist of a hemochorial membrane, labyrinthine venous system and discoid shape (Supplementary Figs S10-S12) and in concordance with previous studies^2^. We also identified several phenotypes that have arisen independently in Eutheria: Epitheliochorial membranes evolved independently in the Lemuroidea, and in the common ancestor of Artiodactyla and Perissodactyla, while endotheliochorial membranes later emerged as a secondary innovation in the Carnivora. Diffuse membranes arose separately in Lemuroidea and in the LCA of Perissodactyla and Artiodactyla. Zonary placental structure evolved independently in three lineages: Carnivora; Paenungulata (hyraxes & elephants), and aardvarks. Similarly, villous membranes emerged independently in four distinct groups: Perissodactyla; Artiodactyla; Cingulata (armadillos), and in Primates. Together, these patterns highlight extensive convergent evolution of placental traits in mammals, implying that similar selective pressures have repeatedly driven the emergence of similar placental phenotypes across diverse lineages, or at the very least that there are fundamental constraints on the combinations of phenotypes that can evolve.

Random forest analysis (with leave-one-out phylogenetic correction) was used to determine the predictability of placental phenotype from miRNA gene family presence/absence. The placental phenotypes of extant species were well predicted by miRNA repertoire, with out-of-bag error rates ranging from 11.9-22.4% when predicting venous pattern, 18.3-23.7% when predicting membrane type, and 1.7-16% when predicting placental shape. While the reconstructed eutherian LCA miRNA repertoire did in all but one case correctly predict discoid/bidiscoid placental shape, it also incorrectly predicted epitheliochorial membrane, and villous venous pattern, instead of the inferred ancestral hemochorial-discoid-labyrinthine combination. Naturally, more recently-evolved miRNAs are not available to use in the predictions for the eutherian LCA. Despite both random forest tests showing high predictability of placental shape from miRNA repertoire, the prediction failure across membrane types and venous patterns suggests extensive regulatory network rewiring throughout eutherian evolutionary history.

### Targets of “core eutherian” and “phenotype-associated” miRNAs display a range of biological functions including developmental processes

The gene targets of miRNAs from the “core eutherian” and “phenotype-associated” sets were predicted using Targetscan^51^ and TEC-miTarget^52^, and subsequently subject to overrepresentation analysis on a by-miRNA, by-species level. Prominent parent terms conserved among the targets of “core eutherian” miRNAs include development-associated terms including ‘positive regulation of cell differentiation’, as well as terms associated with immune function including ‘thymic T cell selection’. For the groups of phenotype-associated miRNAs, parent terms were commonly associated with developmental processes and morphogenesis, including ‘pattern specification process’ and ‘animal organ morphogenesis’ from the targets of miRNAs associated with chorial membrane phenotype, and ‘developmental maturation’ and ‘mesenchyme development’ from the targets of miRNAs associated with placental venous patterning and placental shape. Treemap plots of reduced-terms summaries are shown in Fig. 4. Full target prediction and enrichment analysis outputs are provided in Supplementary Tables S8 & S9, respectively. Reduced-terms summary tables are found in Supplementary Table S13.

**Figure 4.**
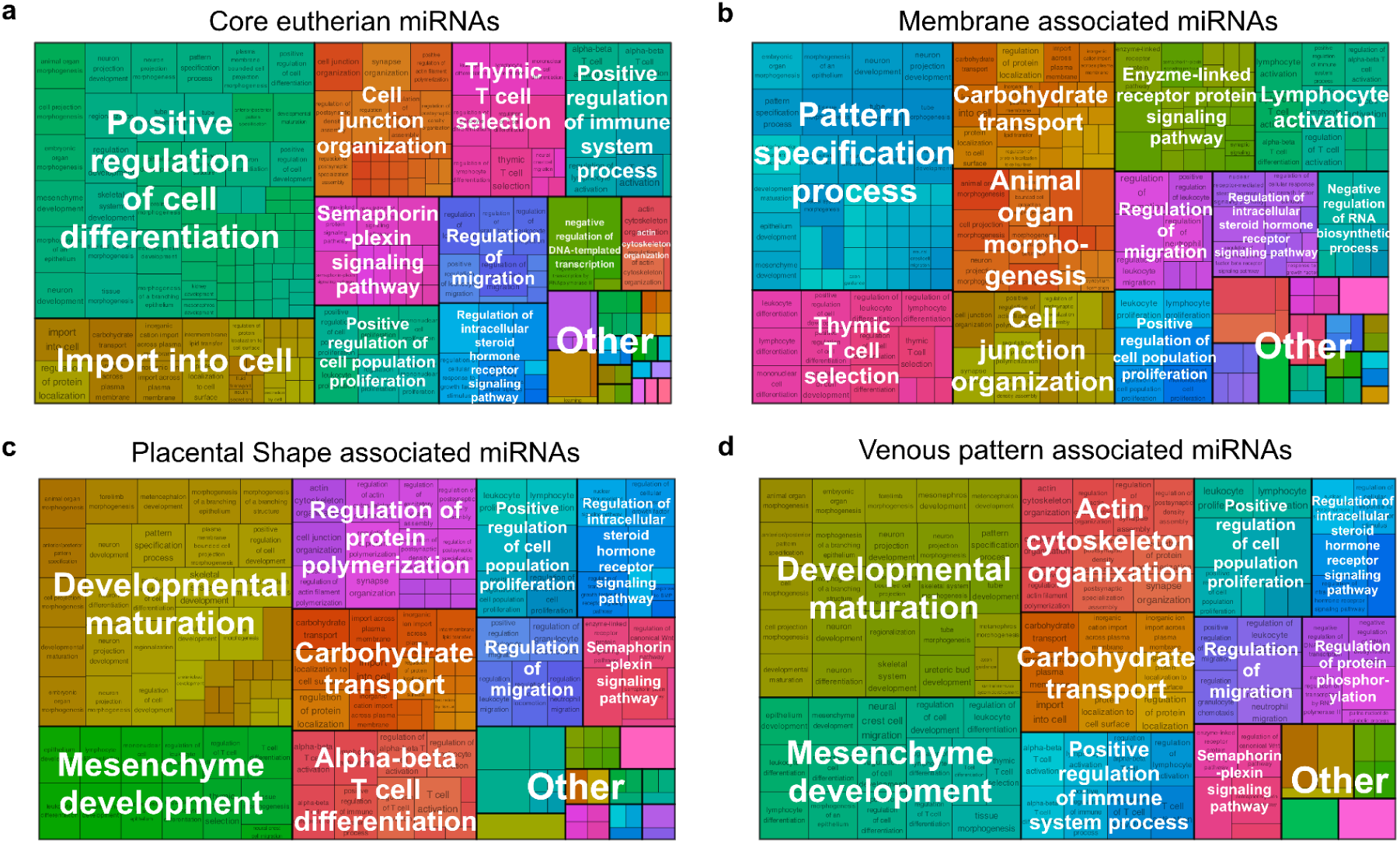
Reduced-Terms Summaries of GO Terms Enriched in Targets of Evolutionarily-Significant miRNA Families. Treemap plots show reduced-terms summaries of gene ontology terms, with each term weighted by the number of occurrences among the top 100 most conserved GO terms enriched among the targets of miRNAs in that group, these groups being: a) “core eutherian” miRNAs, b) chorial membrane-associated miRNAs, c) placenta shape-associated miRNAs, and, d) placental venous patterns-associated miRNAs.

### miRNA Expression Data from Literature

Meta-analysis of publicly-available miRNA expression data for endometrial tissue across seven species datasets taken from (Fig. 5, full species datasets and references in Table S4) revealed expression of all ten “core eutherian” synapomorphic miRNA families in at least one species, with eight (MIR-136, MIR-154, MIR-28, MIR-324, MIR-331, MIR-362,MIR-376 and MIR-423) expressed across all species. Overall, of 136 instances where a phenotype-associated miRNA was present in a species genome across this dataset, there were 81 instances of recorded expression in the endometrium. This pattern varied between species, ranging from 17 of 20 families being expressed in humans, to only 10 of 27 families in goat, though experimental design differences may influence these patterns. Due to the time-sensitivity of gene expression patterns in the endometrium through implantation and pregnancy, a lack of recorded miRNA reads for a given miRNA family should not imply that they are not expressed at any point in the endometrium of that species.

**Figure 5.**
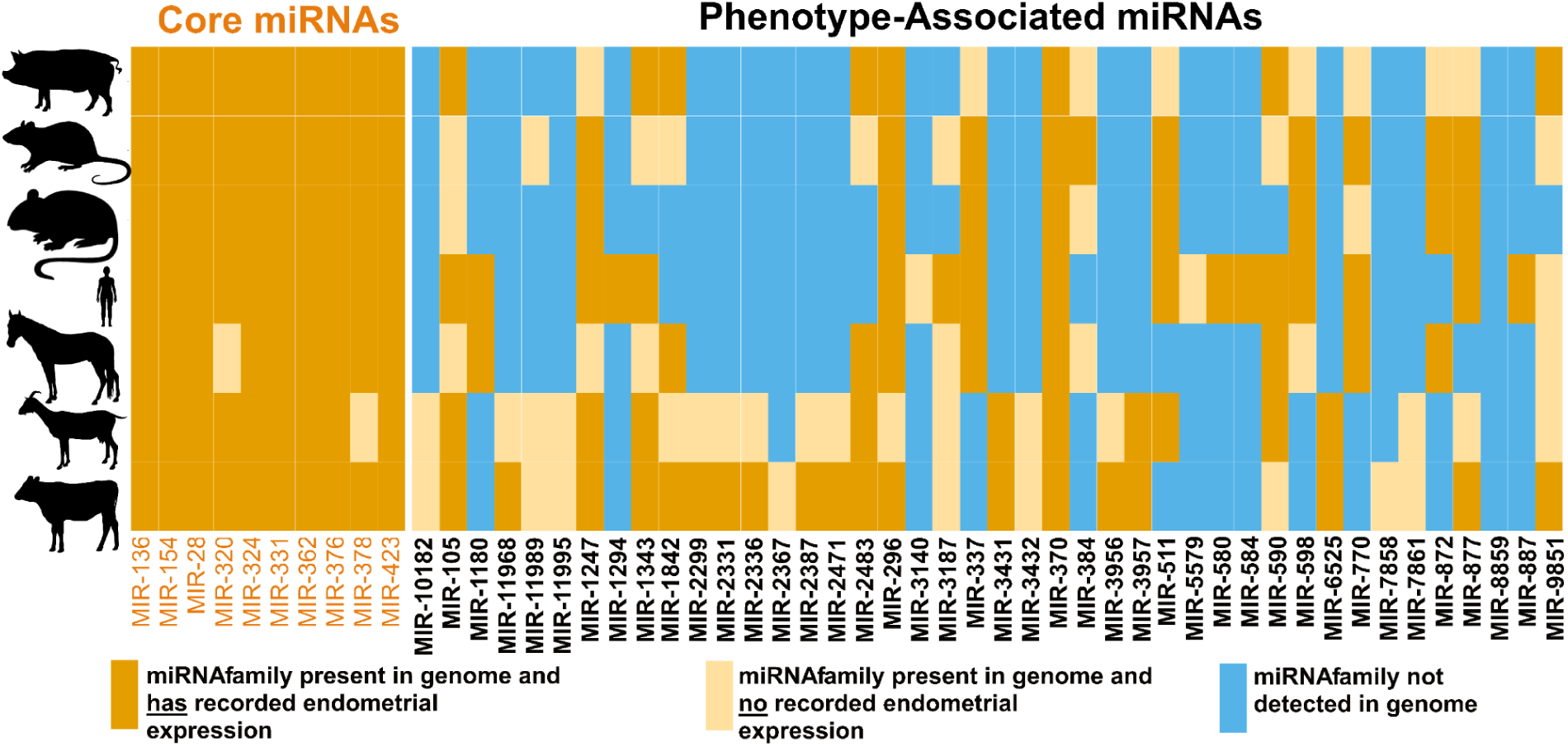
Occurrence of “core eutherian” and “phenotype-associated” miRNA families from endometrial expression datasets across 6 species. The 10 “Core eutherian” miRNA families are represented by the left matrix, and the 42 “phenotype-associated” miRNA families by the right matrix. Species listed on the *y-axis* from top to bottom are Pig, Rat, Mouse, Human, Horse, Goat, and Cow. Orange cells indicate an miRNA which is present in the genome of a species, and has recorded endometrial expression. Pale orange cells indicate an miRNA family which is present in the genome of a species, but has no recorded expression in the endometrium (*i.e.* no RNAseq reads reported from endometrial tissue) in published RNAseq data. Blue cells indicate an miRNA family is not detected in the genome of that species.

## DISCUSSION

Placental evolution shows surprising genomic predictability: miRNA gene family repertoires can reasonably accurately predict placental morphology across mammal species, with convergent phenotypes consistently involving losses of the same miRNA families. The ability to accurately predict placental characteristics from miRNA gene family presence-absence patterns indicates that developmental constraints channel placental innovation through limited genetic pathways rather than allowing unlimited mechanistic diversity, and that there is a tight link between evolutionary patterns for these particular elements and the resulting phenotypes. The independent evolution of these tight genotype-phenotype pairs suggests that there are only a limited number of evolutionary avenues available for placental diversity. The mammal placenta is a complex organ, its evolution having required exaptation of maternal and foetal tissues, as well as co-option of existing networks of gene regulation, in a manner which may be evolutionarily constrained, as suggested by the convergent evolution of viviparity in mammals and squamates^1,53–55^. It stands to reason that similar patterns of evolutionary constraint may influence the diversification of placental form and function, such that shared physiological and genomic characteristics in distantly-related taxa may lead to the evolution of similar placental phenotypes, given a similar background of selective pressures.

Our reconstructions suggest multiple independent losses of MIR-872, which is strongly associated with the absence of a zonary placenta. The level of loss observed seems to conflict with the documented rarity of miRNA gene loss in evolution. Two alternative explanations merit consideration: firstly, rapid sequence divergence may cause some miRNA families to become undetectable rather than truly lost. Alternatively, the commonly held assumption of single-origin for miRNA families may be incorrect, and multiple independent gains are not accounted for by the taxonomy of miRNA gene families used in this study^42^. Future analyses that account for similarity and convergence between mature miRNA sequences from different miRNA families will provide further insight. Irrespective of the underlying evolutionary mechanism, the genomic presence-absence patterns for recognisably homologous miRNA families, remain strongly predictive of placental phenotype. This suggests that whether through loss, gain, or sequence divergence, the detectability of these miRNA families correlates strongly with specific developmental programs underlying placental morphology.

Ancestral reconstruction of miRNA gene families showed one miRNA family which arose at the ancestor of ruminants (MIR-11968), a subclade of which *(Pecora)* has a characteristic cotyledonary placenta. This gene is part of a group of 15 cotyledonary placenta-associated miRNA families which arose at this node, with 2 others arising at the LCA of Artiodactyla. Of these, only MIR-11968 showed presence in every taxon tested within Pecora and no others, and as such it is the strongest candidate within our phenotype-associated miRNA gene families of a novel miRNA gene family directly involved in a marked shift in placental phenotype, and being necessary to maintain that phenotype. The detection of MIR-11968 in bovine glandular endometrial epithelial organoids but not in human Ishikawa cells, as well as its presence in the caruncular tissue *in vivo* provides strong evidence that this miRNA gene family is involved in the peri-implantation period of pregnancy. Our *in vitro* and *in vivo* analyses of MIR-11968 demonstrated persistent expression in relevant reproductive tissues and supported a causal effect on cotyledonary placental shape, showing that this miRNA-phenotype relationship is not simply correlational. The opposing patterns of expression of MIR-11968 in glandular and aglandular cell types in pregnant vs non-pregnant animals, suggests that the presence of the embryo influences upregulation of MIR-11968 in non-glandular cells of the endometrium via conceptus-derived molecules or maternal-conceptus interactions. A similar pattern is observed in MIR-5579 and MIR-887 which arose at a node ancestral to the *Platyrhinni*, which have characteristic trabecular placental venous pattern.

Dense genomic sampling revealed that many previously identified “core eutherian” miRNA families show sporadic losses, contradicting assumptions about their universal requirement for mammal pregnancy. While miRNA loss is typically rare, these specific losses show no significant phenotypic associations and mostly occur in taxa retaining ancestral placental states. The losses may affect developmental processes not examined here, such as implantation strategy, rather than the three placental phenotypes analysed in this study.

While the targets of miRNAs are diverse across Eutheria, the targets of both the “core eutherian” and “phenotype-associated” miRNA families identified in this study have biological processes which show potential association with the development of complex organs like the placenta. While “core eutherian” and “phenotype-associated” miRNA families show developmental pathway enrichment, the low conservation of specific predicted pathways across species suggests that placental evolution operates through distributed regulatory networks, rather than discrete pathway switches. This pattern indicates that miRNAs quite likely influence placental phenotypes through coordinated fine-tuning of multiple regulatory circuits. Evolving such a mechanism may be more robust than relying on single developmental pathways, or the focussed activity of a single protein-coding gene. The evolution of placenta through miRNA regulation stands in contrast to the way that, say, Homeobox genes operate in development. Homeobox genes act as master switches, defining broad body plans with discrete, often dramatic effects^56,57^, like positioning major body segments or specifying cell lineages^58,59^. In contrast, miRNAs simultaneously fine-tune the expression of thousands of genes across complex regulatory networks. Where homeobox gene mutations can directly affect radical developmental disruptions, changes in miRNA repertoire clearly produce subtle modifications that manifest as alternative placental phenotypes. This means that miRNAs can generate nuanced phenotypic diversity through distributed, coordinated genetic adjustments across entire developmental landscapes.

The outcome of miRNA repertoire evolution has been a sophisticated fine-tuning of placental development. The extensive gene target repertoires, typically numbering in the thousands, are characteristic of miRNAs and make them ideally suited for such distributed regulatory control, potentially explaining both their evolutionary plasticity but also the predictable phenotypic outcomes we observe. We show that placental phenotypes are predictable from miRNA gene repertoires across 124 mammal species with available genomic and phenotype data, with convergent morphologies consistently involving the same miRNA families. This genomic predictability reveals developmental constraints that channel placental evolution down a small number of avenues, *via* both miRNA gene gain and loss events. Experimental validation of MIR-11968 in bovine caruncular tissue provides direct evidence linking clade-specific miRNA evolution to morphological innovation.

The current findings establish miRNAs as key molecular architects of subsequent mammal reproductive diversity and provide a framework for understanding how genomic constraints shape complex trait evolution. Importantly, placental phenotypic diversity has not emerged through a single genetic master switch, but through this complex, multilayered regulatory landscape. It has been coordinated by a network of miRNAs, some ancient and persistent fixtures in mammal genomes since the origin of Eutheria, others displaying more variable patterns of presence and absence.

## Methods

Placental phenotype data was obtained from the set originally collated in Wildman et al^2^, with these phenotypes verified by reference to the original source material to check whether original categorisations should be updated. In total, 27 new phenotype states for species with associated genomes were added, and 30 updates to existing phenotype states were input with reference to original descriptions and sources. Amendments and additions to the Wildman paper are given in Supplementary Table S2. A total of 124 species with genomes available also had associated placental phenotype data - specifically, the state of the chorial membrane (hemochorial, endotheliochorial or epitheliochorial) the venous patterning of the placenta (labyrinthine, villous or trabecular) and the shape of the placenta as a whole (discoid/bidiscoid, diffuse, zonary or cotyledonary).

All 231 mammal genomes available as part of the Vertebrate Genomes Project Phase I data freeze had miRNA repertoires annotated using MirMachine 0.3.0.1^48^. Three genomes were omitted as their miRNA detection step failed: GCF_030020395 *(Manis pentadactyla)*, GCA_951394435 *(Stenella coeruleoalba)* and GCA_045843805 *(Urocitellus parryii)*. The remaining mammal genomes (*i.e. n=228*) and their associated miRNA annotations were incorporated for this study. To maximise on species representation from our phenotype dataset, the 228 VGP mammal genomes were supplemented by genomes released as part of the Zoonomia (*n=136*) and DNAzoo (*n=142*) repositories. Ultimately the goal was to ensure we had representation across all species for whom we had placental phenotype data.

MirMachine 0.3.0.1^48^ was used to detect miRNA families across genomes, using the ‘Mammalia’ node and the ‘--add-all-nodes’ flag. The resulting filtered GFF files were converted to a presence/absence matrix in R. In addition, BLAST was used to search for known mammalian pre-miRNA sequences as recorded in MirGeneDB, in order to reduce the prevalence of false negatives and include miRNA families not used in MirMachine. An e-value cutoff of <1e^-5^ was used. BLAST output files were also converted to a presence/absence matrix, and combined with the one produced by MirMachine. Genomes were filtered by removing any which did not have the full set of 53 miRNA families highly conserved across all Vertebrata (allowing for 1 gene absence, in the case of a rare loss event). Following this filtering, genomes of 377 eutherian species, 19 marsupials and 2 monotremes were included for further analysis (full details of genomes in Supplementary Table S1). The “core eutherian” miRNA set were identified as miRNA families completely absent from marsupial and monotreme mammal genomes but present and extremely rarely lost* in Eutherian genomes (*per miRNA family we permitted loss in a single eutherian species only to allow for low-level detection error).

To obtain a phylogeny for downstream analysis, the ROADIES-based phylogeny developed as part of the Vertebrate Genomes Project Phase I^43^ used for downstream analyses. For species not represented in this tree, a consensus tree was formed with additions from TimeTree, using the VGP phylogeny as a backbone. Where there was no placement for a given species in either tree, it was placed randomly among congenerics. The consensus tree was time calibrated using the penalised-likelihood estimation method *chronos* in the *ape* R package^60^, using calibration points from Yu et al^61^, supplemented with dates on additional nodes taken from Alvarez et al^62^ (dating calibration information found in Supplementary Table S3), and parameters (‘correlated’ model, lambda smoothing parameter=0.01) selected according to minimal PHIIC values and maximal penalised log likelihood scores.

To ascertain levels of co-occurrence between miRNA gene families, and the presence of any defined clusters (of miRNA families themselves, and of species according to their miRNA repertoires), the miRNA gene family presence-absence matrix was used to compute Jaccard dissimilarities between miRNA gene families across all genomes, and between genomes. Hierarchical clusters were generated from the resultant dissimilarity matrices, using the *hclust* function of the *stats* package. Clustering methods were compared by bootstrapping for optimal stability using the *clusterboot* function of the *fpc* package^63^, using the Ward D2 clustering method. The *MetaMDS* function of the *vegan* package^64^ was used to visualise these clusters and extract the weightings of the distinguishing miRNA gene families. As a broad measure of phylogenetic signal in species miRNA repertoires, the Jaccard Robinson Foulds distance between the consensus phylogeny and the tree derived from hierarchical clustering of species by miTNA presence was computed.

In order to detect miRNA families potentially involved with the evolution of convergent placental phenotypes, a phylogeny-aware random forest approach was used. A cophenetic distance matrix was derived from the consensus tree using the *cophenetic.phylo* function of the *ape* package^60^, which was then subject to principal coordinate analysis. The number of relevant principal coordinate axes was determined using the broken-stick method, via the *bsDimension* function of the *PCDimension* package^65^. The by-species scores for these axes, inferring relevant phylogenetic information, were included alongside the presence of every mammal-specific miRNA as predictors of placental phenotype across the three categories. In order to further minimise phylogenetic signal, a ‘leave-one-out’ approach was used, in which each taxonomic order with at least 5 species represented was, in-turn, excluded from each random forest (taxonomic orders taken from NCBI). For each forest, a 50:50 training/testing split was used, with each forest using 5000 trees. To infer predictability of placental phenotype from miRNA repertoire alone, predictions were made from each forest, using only this information, and the out-of-bag error rates calculated. As a statistical control, we repeated all random forest analyses, using miRNA presence/absence data that was at the same prevalence as real data, but randomly distributed across the eutherian tree. For each phenotype, the variable importance scores for each excluded-order run were collated, and the minimum importance factor extracted for each. miRNA families were deemed to be associated with a phenotype if their variable importance score was greater than the mean value for that analysis, plus one standard deviation, and was more than twice that of the randomly-distributed control run for that miRNA family.

### Evolink

Due to the leave-one-out approach used in the random forest, phenotypes-genotype associations with very strong phylogenetic signal are at risk of being undetected. For instance, if an miRNA family and a specific phenotype are specific to one order of mammals, the significance of that miRNA would be null when that order was excluded. While associations with such strong phylogenetic signal are problematic, in that it may be impossible to statistically distinguish the relative contribution of gene presence and phylogeny, we still wanted to detect these associations and report them, with this caveat. To that end, Evolink analysis^66^ was also performed to find phylogeny-corrected association statistics between miRNA presence/absence and placental phenotype. For each phenotype state, positive and negative associations between miRNA gene presence and phenotype presence were recorded.

### Phenotype-Associated miRNA Gene Families

A final shortlist of 42 miRNA gene families (hereafter termed ‘phenotype-associated miRNAs’) was taken from the overlap of the random forest and Evolink outputs (*n=20*), as well as any significant results from Evolink which would be excluded due to the clade-specificity of the miRNA gene in question (*n=22*).

### miRNA Ancestral Reconstruction

For each mammalian miRNA, ancestral reconstruction by Dollo parsimony was performed in order to determine patterns of miRNA gain and loss across the mammal tree, using the *map_dollo_changes* function of the *Claddis* package^67–69^. Each reconstruction was performed 1000 times. Where reconstructions disagreed on the placement of gene gain and losses, the set of node(s) suggested by the plurality of runs was accepted. These reconstructions were used to determine the miRNA repertoires of the eutherian, therian and mammalian ancestors. By using the reconstructed miRNAs of the eutherian ancestor, we tested whether the predicted ancestral phenotype could be predicted from miRNA alone, using our previously-created random forest models. The ancestral reconstruction of the phenotype-associated miRNAs was compared with the phenotypic reconstructions to determine whether miRNA gene families were gained or lost prior to-, coincident with-, or after-, the emergence of derived phenotypes.

### Target Prediction

In order to ascertain the functional role of miRNAs from both “core eutherian” and “phenotype-associated” categories, target prediction was performed on 66 eutherian genomes produced as part of the Vertebrate Genomes Project which have accompanying egapx genome annotations. The sequences of 3’ UTR regions mapped as part of these annotations were extracted, as were miRNA seed sequences predicted through MirMachine. These sequences were used to perform target prediction using TargetScan. To refine the target predictions for each miRNA and minimise the prevalence of false-positive target predictions, a machine-learning approach, TECimiTarget^52^, was used to validate seed sequence-UTR matches, with each match being scored for validity according to the provided model (trained on the miRaw dataset from TEC-miTarget). Any targets with prediction scores of 0.5 or higher were brought forward for further analysis.

The resulting lists of predicted gene targets were used to perform, for each miRNA family in each species, overrepresentation analysis, using the full repertoire of predicted targets in each species as the background gene universe. For each miRNA, the top 100 GO terms by species representation (excluding species in which a given miRNA is not present) were extracted. To examine broad groupings of biological pathways across targeted genes, both within the “core eutherian” miRNA set, and within each group of “phenotype-associated” miRNAs (*i.e.* membrane-associated, venous pattern-associated and placenta shape-associated), the number of miRNAs in which each top 100 term was found was used as a scoring metric for reduced terms analysis (clustering threshold=0.8.) For example, if GO term X was enriched among the targets of 8 miRNA families, its scoring metric would be 8.

### miRNA Expression Data from Literature

Endometrial miRNA expression data was compiled from published datasets for 7 eutherian species as of April-May 2024 (Supplementary Table S4), and the presence of phenotype-associated miRNA expression across these species was determined. In cases where the miRNA is present in a species’ genome but there are no RNAseq reads recorded for its expression in endometrial tissue, this was taken as a lack of recorded expression. However, this does not preclude the potential expression under different experiential conditions, *i.e.* under different detection methods or different timepoints of endometrial development during pregnancy. It was not possible to analyse the expression of the miRNA gene families in placental tissue due to reduced availability of cross-species datasets in public repositories.

### MIR-11968 Expression Analysis

Phenotype-associated miRNA gene family MIR-11968, uniquely associated with cotyledonary placenta shape, was taken for further functional analysis. The expression levels of three MIR-11968 paralogs were experimentally assessed in Bos taurus (a model cotyledonary species). We used *in vivo* bovine endometrial samples collected on Day 16 of pregnancy (the day of pregnancy recognition) following embryo transfer of *in vivo* (*n=6*), *in vitro* (*n=8*), or cloned (*n=9*) embryos on day 7 of a synchronised cycle as described in De Bem et al ^70^. Samples were collected from caruncular (aglandular regions and the site of cotyledonary formation) and intercaruncular regions from each animal (*n=23*). Bovine endometrial luminal epithelial (*n=3*) and stromal (*n=3*) cells were isolated from reproductive tracts collected from an abattoir and grown *in vitro* as described in (Tinning *et al.*, 2020)^71^. Stromal and luminal epithelial cells were separately treated with vehicle control (PBS) or 1μg/ml recombinant ovine IFNT (roIFNT) for 24 hours. Bovine endometrial epithelial glandular organoids (*n=3*) were derived from reproductive tracts collected from an abattoir^72^ and treated with vehicle control (PBS) or 1μg/ml recombinant roIFNT for 24 hours on day 8 of growth post isolation. To act as a control human immortalised endometrial epithelial cells (Ishikawa cells; *n=3*) were grown to confluency in DMEM/F12 with 10% FBS (fetal bovine serum) and 1% GSP (glutamine, streptomycin and penicillin). Untreated cells were plated (approximately 200,000 cells per well in a 6 well plate) for 24 hours. All RNA was extracted using the miRNeasy Mini Kit as per manufacturers’ instructions (Qiagen N.V., Hilden, Germany). For all samples, RNA concentration was measured using the DeNovix DS-11 FX+ spectrophotometer (DeNovix Inc., Delaware, USA) and diluted to 5ng/μl using DNase/RNase free water. The miRCURY LNA RT Kit (Qiagen N.V.) was used to reverse transcribe cDNA from RNA. MiRCURY LNA miRNA PCR Assays for bta-miR-11986 (miR-11968 paralog described in miRBase, but not MirGeneDB), bta-Mir-11968-P1 and bta-Mir-11968-P2 were obtained from Qiagen. cDNA was diluted 1:60 for use in the assays and 3μl added to each well along with 5μl SYBR green, 1μl PCR assay and 1μl water. Real time qPCR was performed on the LightCycler 96 (Roche, BAsel, Switzerland) and LightCycler 96 software used to calculate absolute quantification of results. Data was then exported to Microsoft Excel for further analysis and statistical tests and graphing was performed using Prism Graphpad. A one-way ANOVA with multiple comparisons test was performed to compare bta-Mir-11968-P1 expression in *in vitro* cell types with and without roIFNT treatment. A paired t-test was used to compare caruncular with intercaruncular expression of bta-miR-11986, bta-Mir-11968-P1 and bta-Mir-11968-P2. For the two MIR-11968 paralogs described by mirGeneDB (P1 & P2) we wanted to know whether there were specific functional pathways overrepresented within their pools of targets (as predicted from TargetScan and TEC-miTarget analyses). This was done using the cow-specific gene ontology database (Org.Bt.eg.db)^73^.

## Acknowledgements

This work was supported by the Biotechnology and Biological Sciences Research Council (award numbers BB/X007332/1, BB/X007367/1 and BB/R017522/1) and FAPESP (2016/50433-9, 2016/22790-1, 2017/50438-3 and 2018/14137-1). MJO’C would like to thank the Leverhulme Trust for her personal fellowship to complete this work (RF-2024-492). This research received general VGP support from the Howard Hughes Medical Institute and Rockefeller University start-up funds to EDJ. We also thank DNAzoo and Zoonomia for open access to their range of mammal genomes, and the high-performance computing service at the University of Nottingham for access to computing facilities. BF acknowledges funding through the Tromsø forskningsstiftelse grant (TFS) [20_SG_BF ‘MIRevolution’].

## Author Contributions

MJO’C and NF conceived the project. JF designed and carried out all computational and statistical analyses with input from MJO’C and JOMcI. JCE, HT and ZB designed and performed *in vitro* analyses with input from NF. NF, FVM, JdS, FP, GP, and TDB designed the *in vivo* experiment. HT, TDB, AB, IME, AGD, JdS, FP, GP, FVM and NF performed the *in vivo* experiments. NF, HT, IME, and JCE performed the *in vivo* sample analyses. OA collated endometrial miRNA expression datasets, VO prepared placental morphology data, MS contributed to generating the consensus tree for phylogeny-informed analyses, VO and BF provided technical assistance with MirGeneDB. GF and EJ provided access to data and critical methodological insights. All authors contributed to writing the manuscript.

