## Supplementary figures and images for "Mammal placental phenotypes are predictable from microRNA repertoires"

### Figure S10

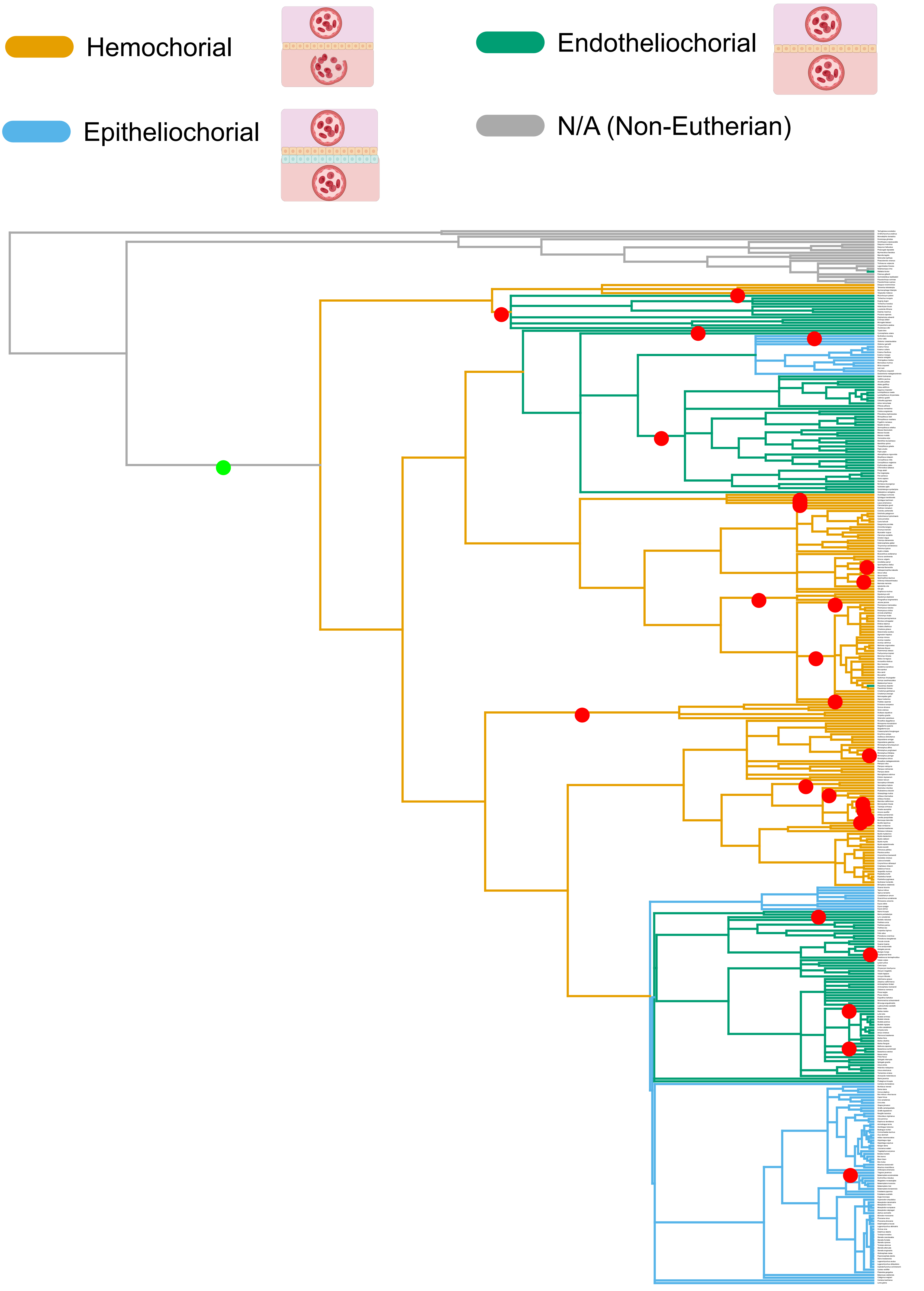

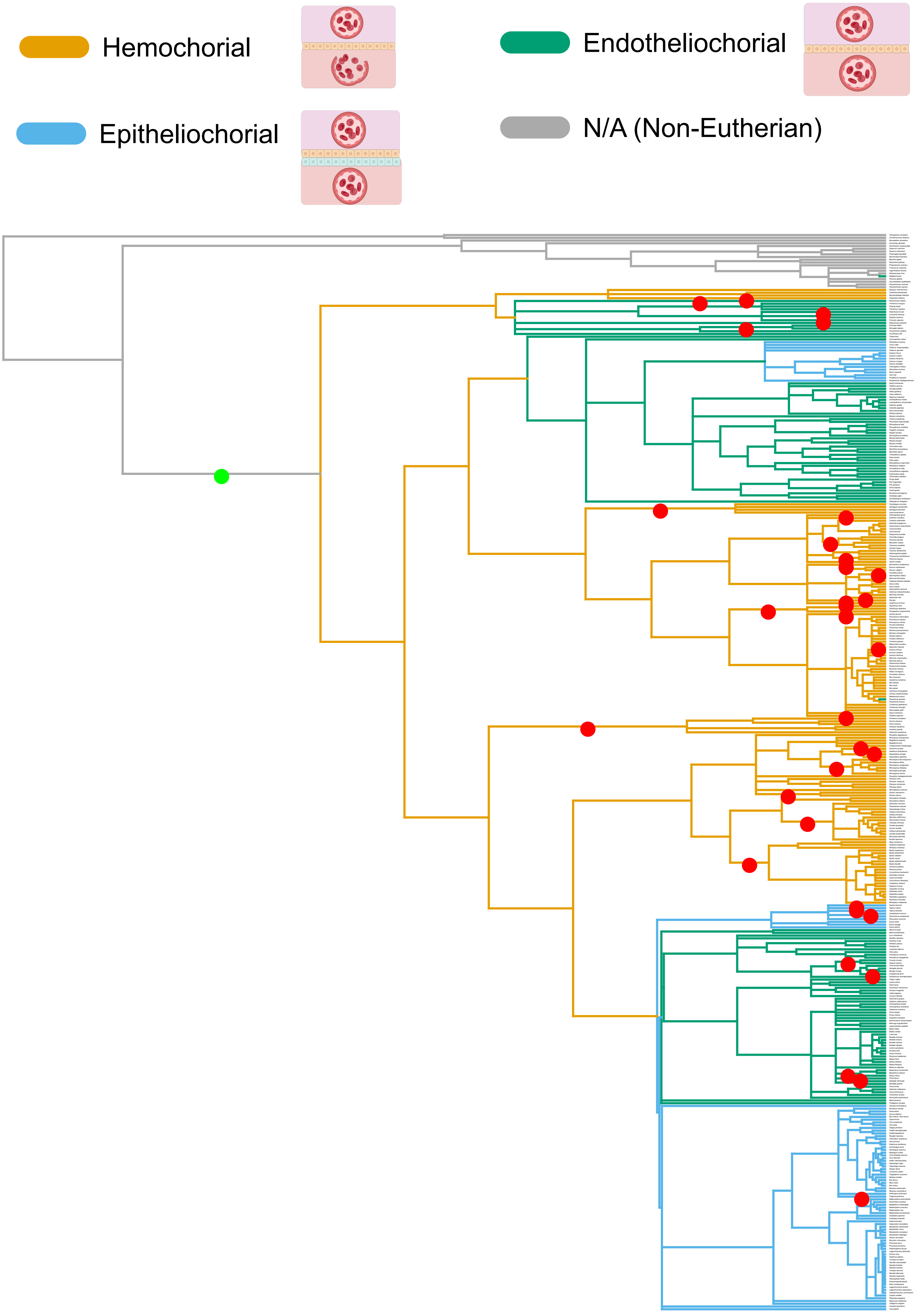

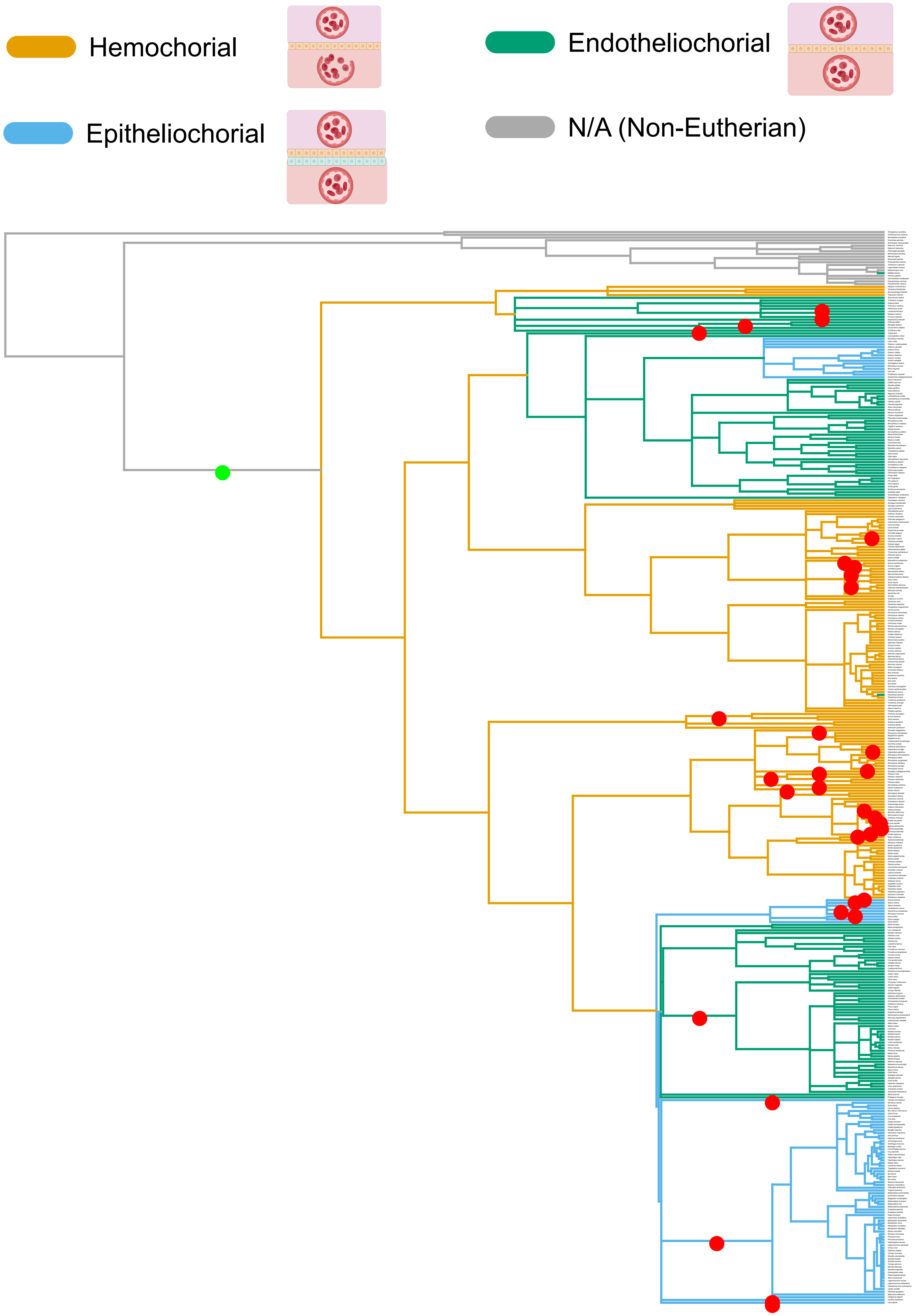

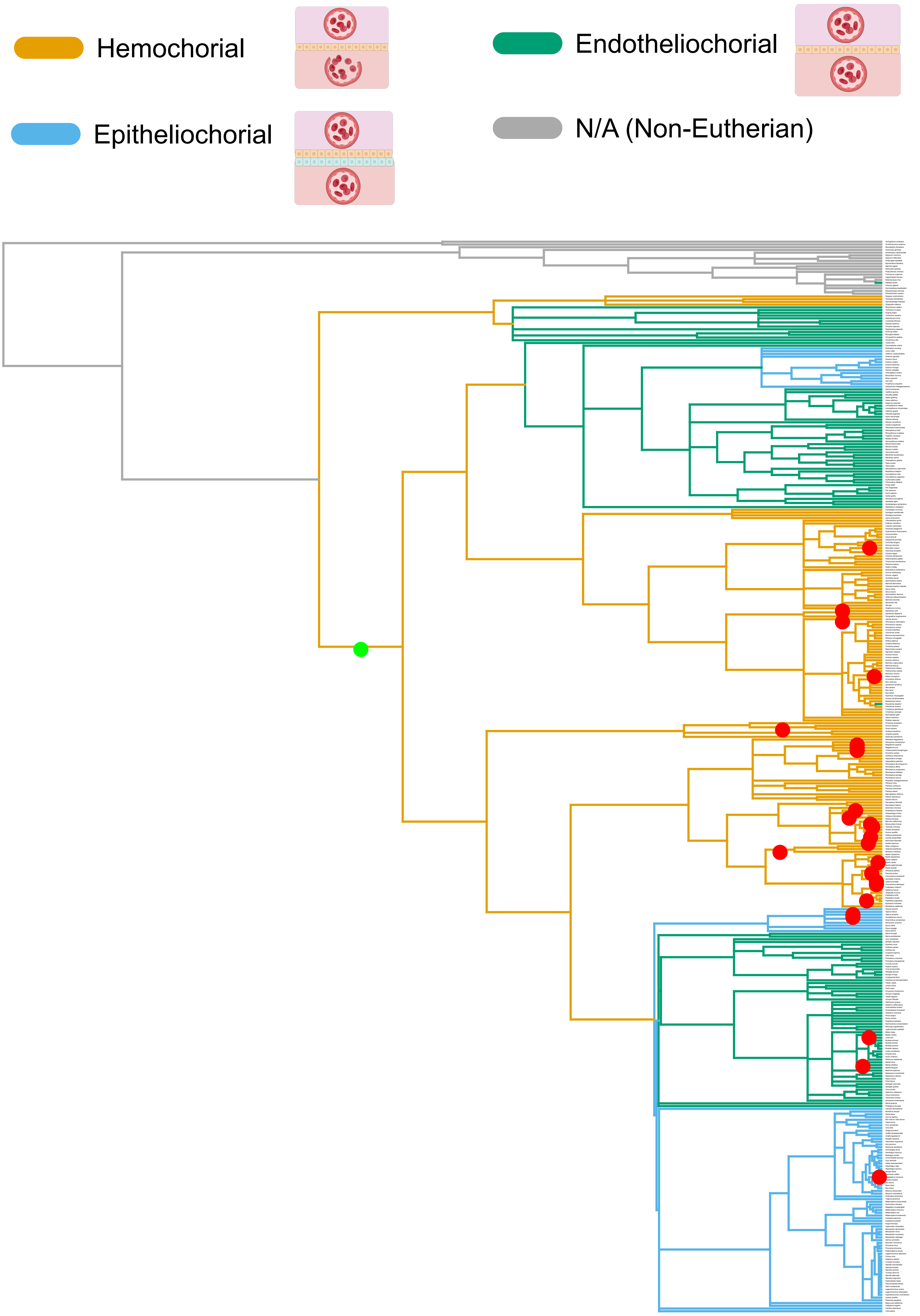

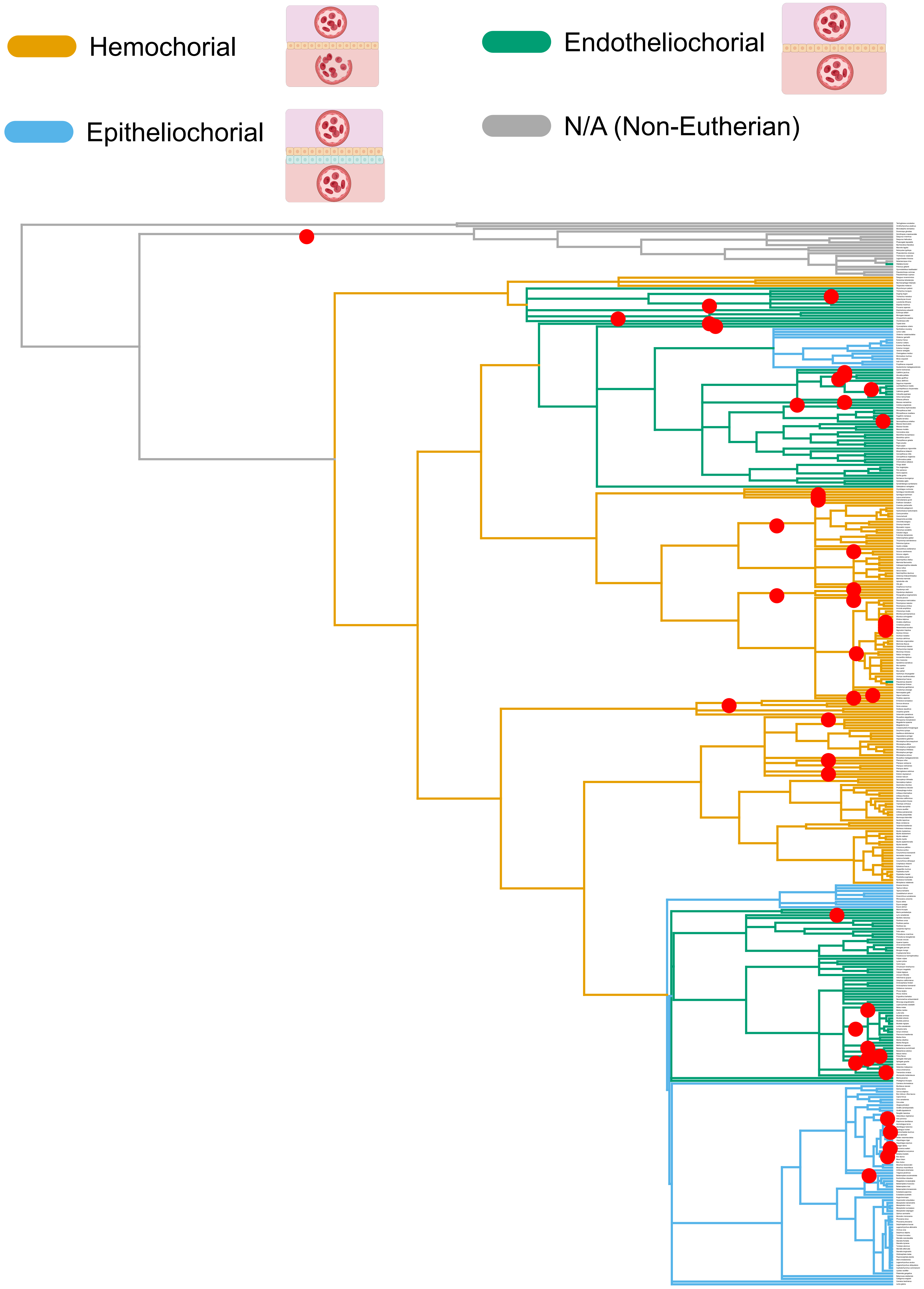

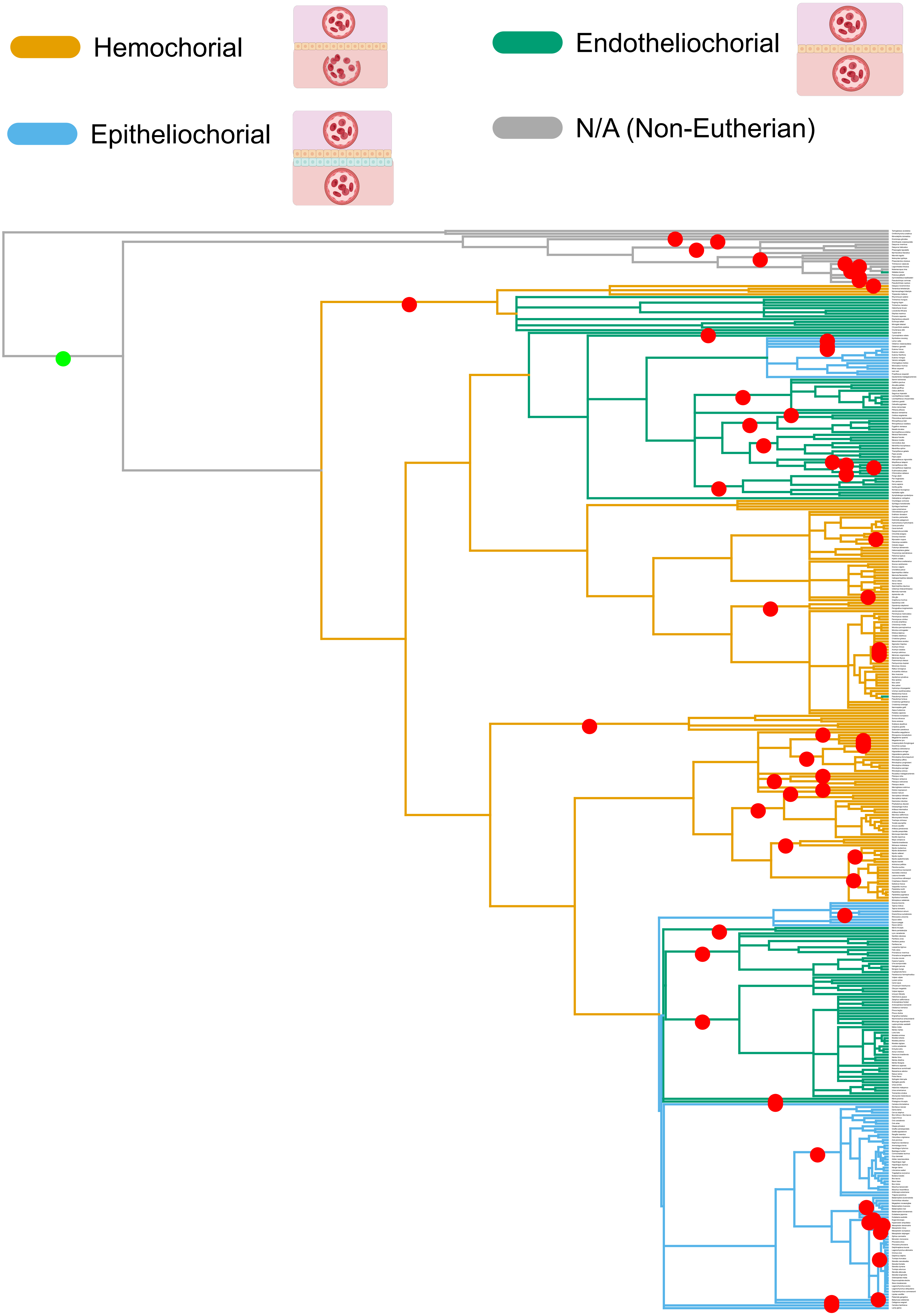

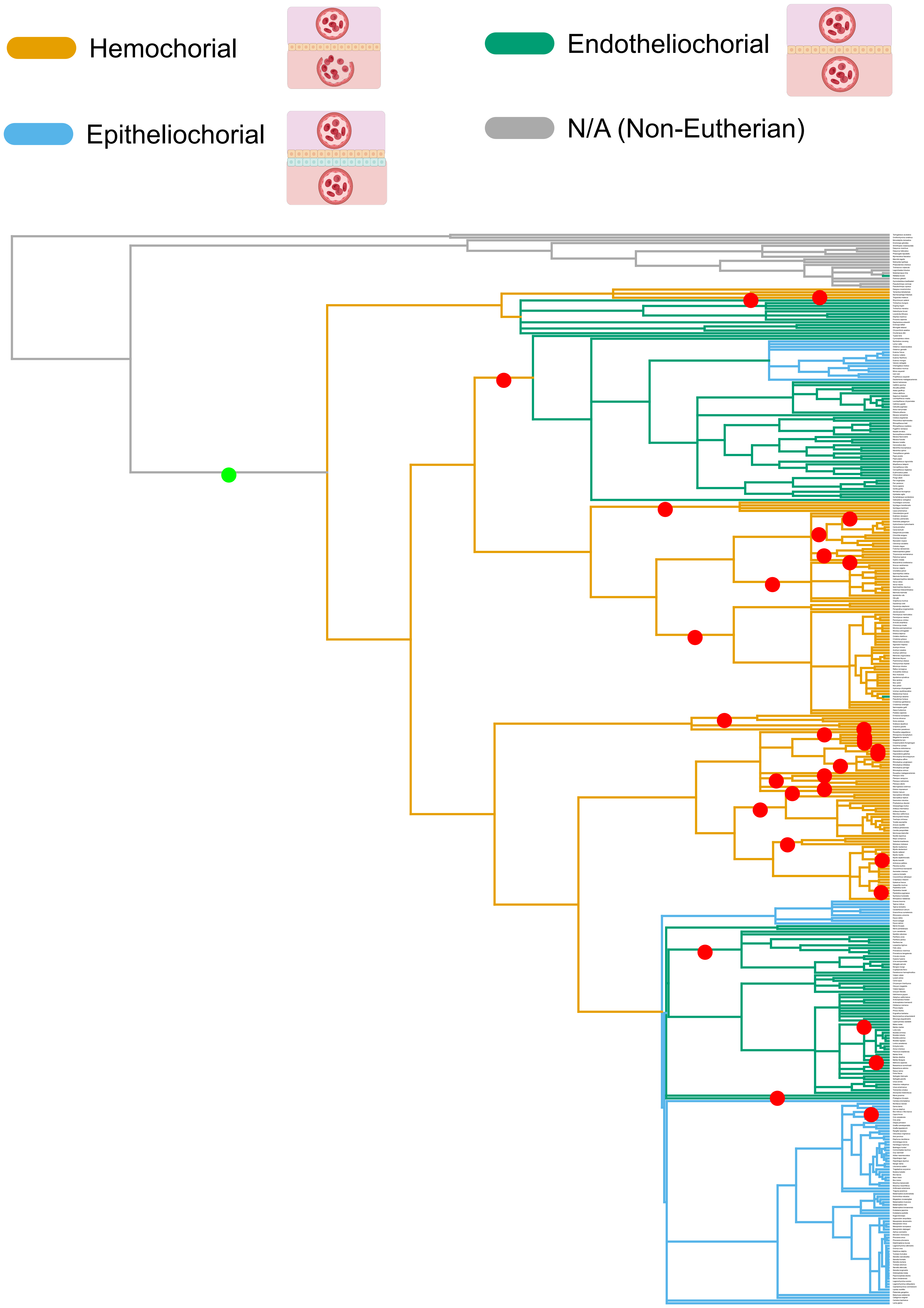

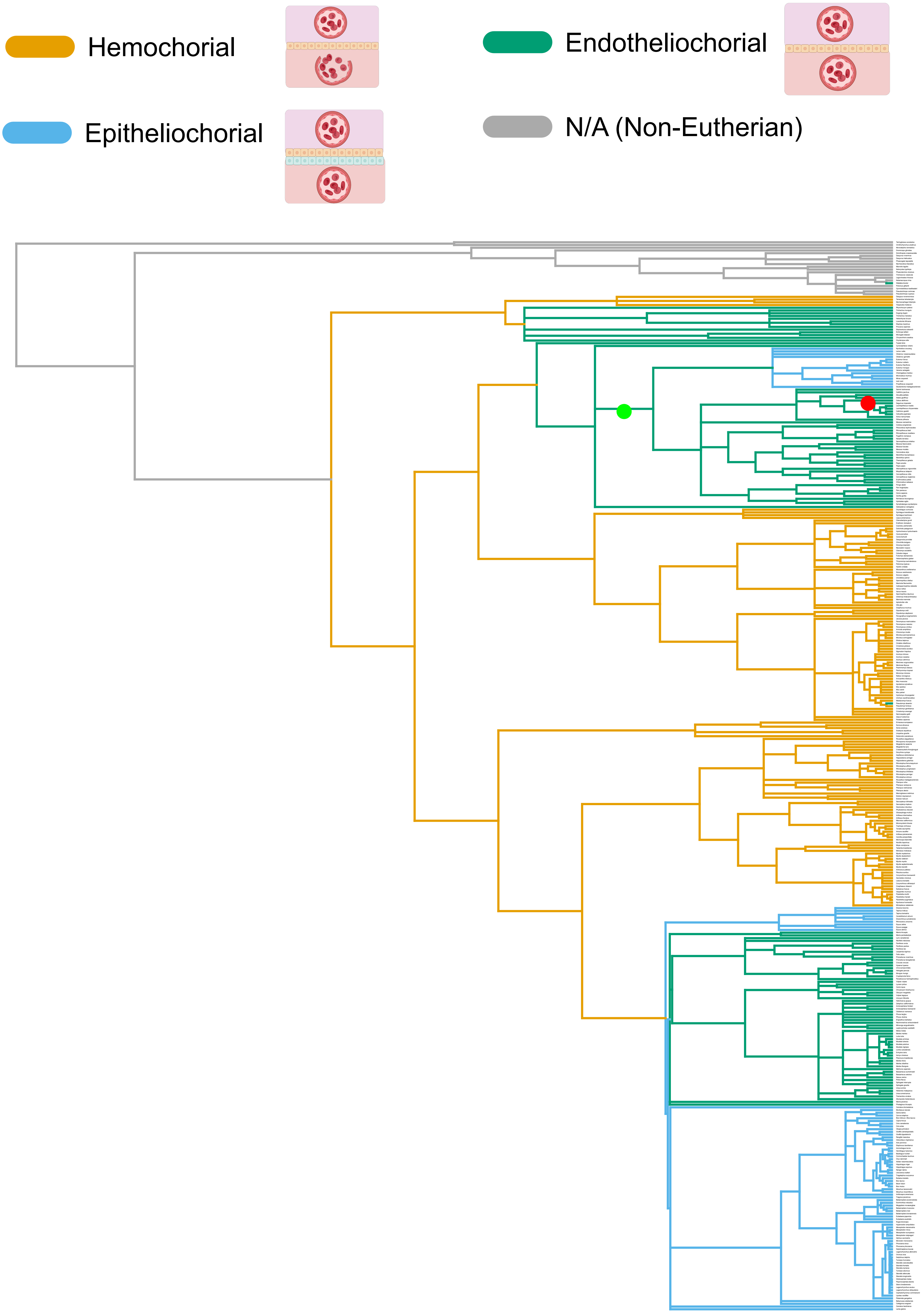

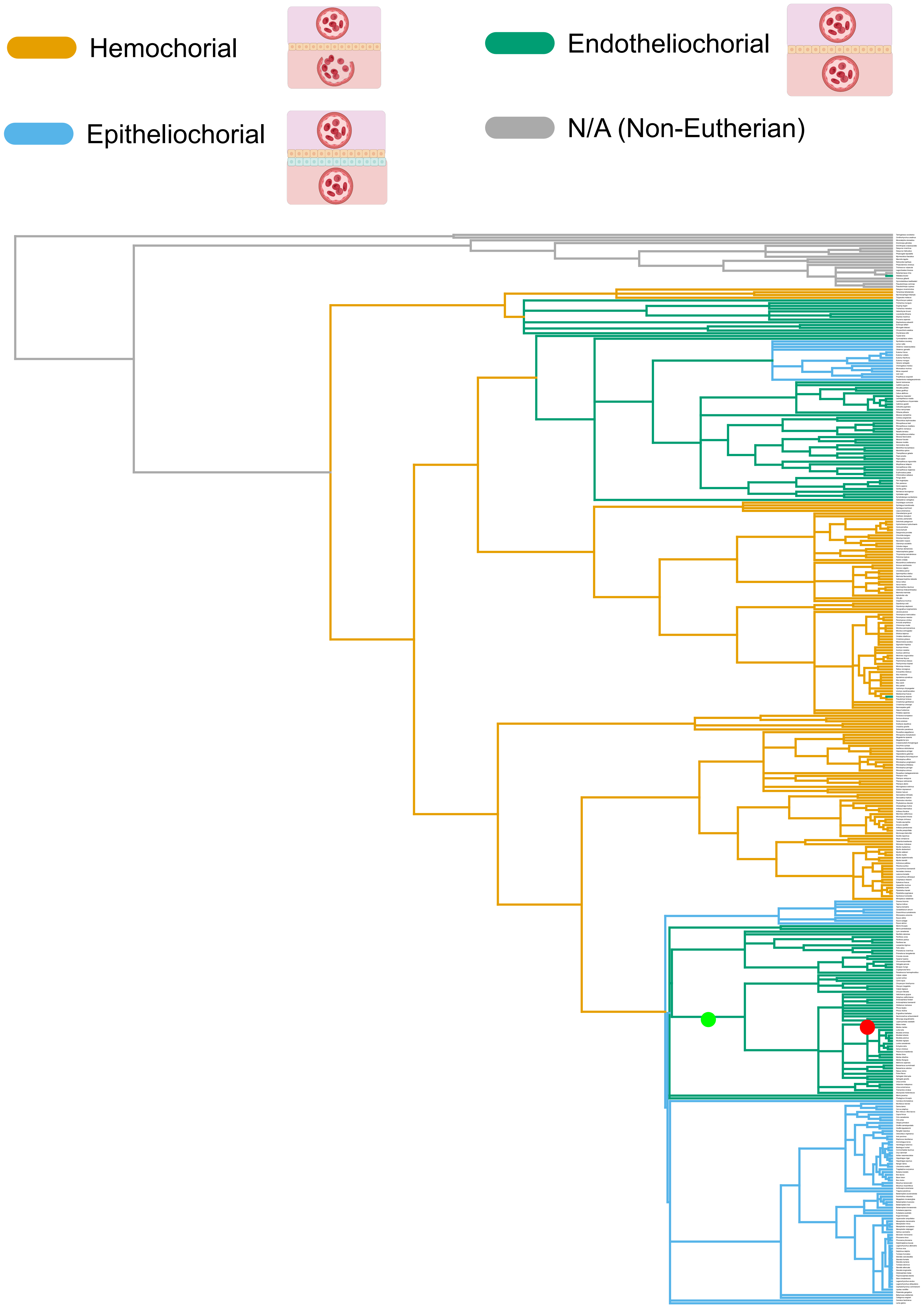

### Figure S12

#### MIR1180

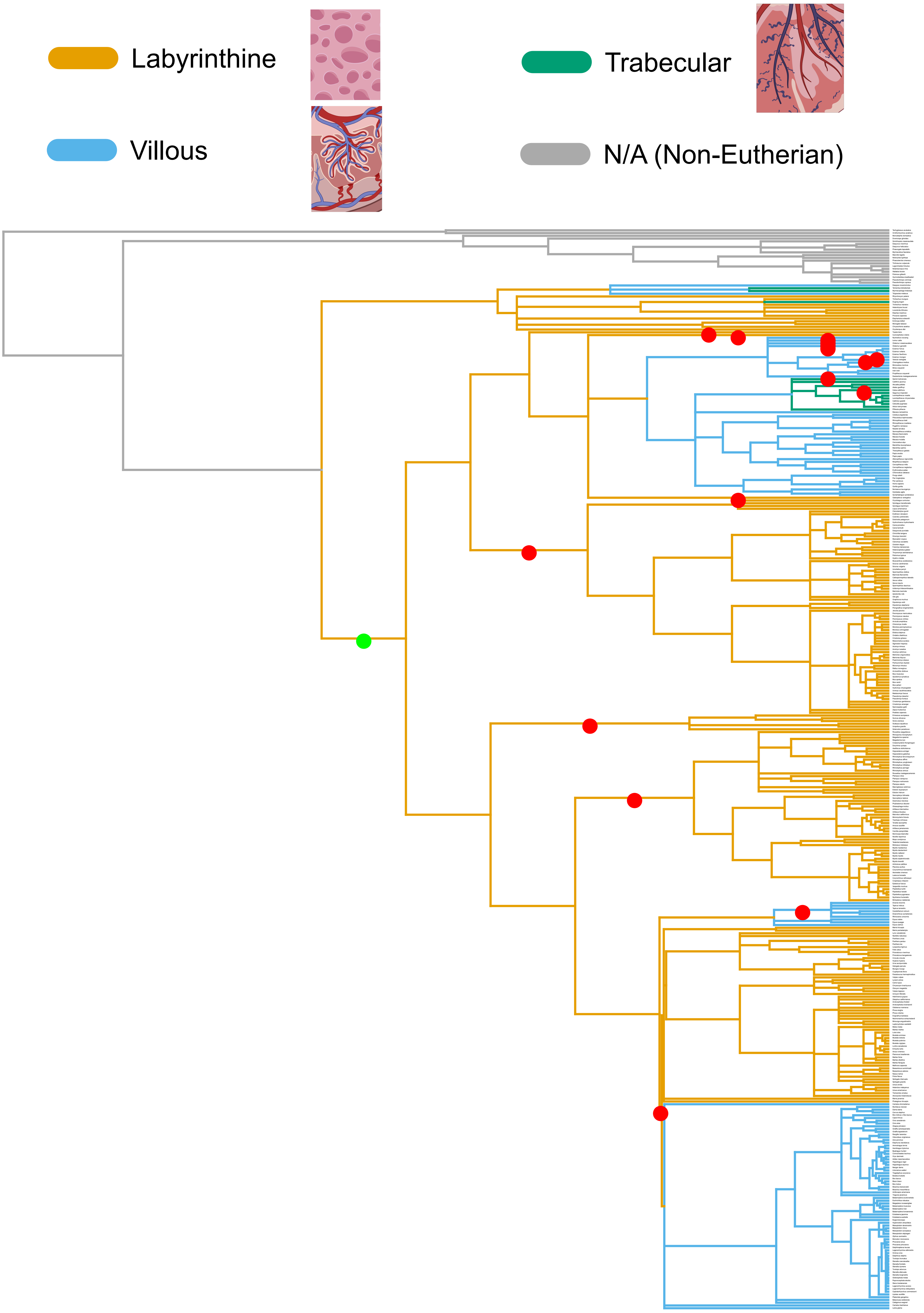

#### MIR580

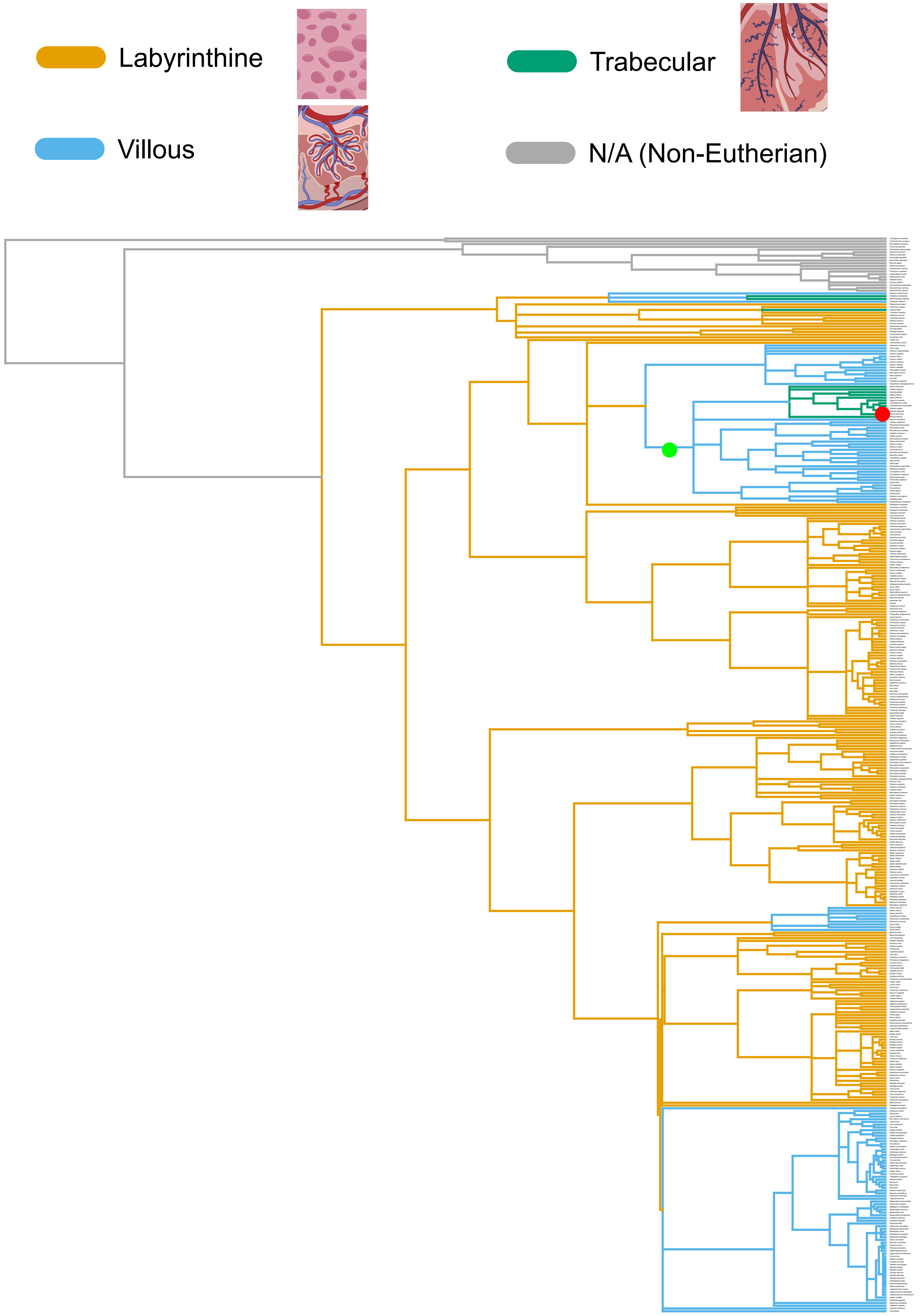

#### MIR584

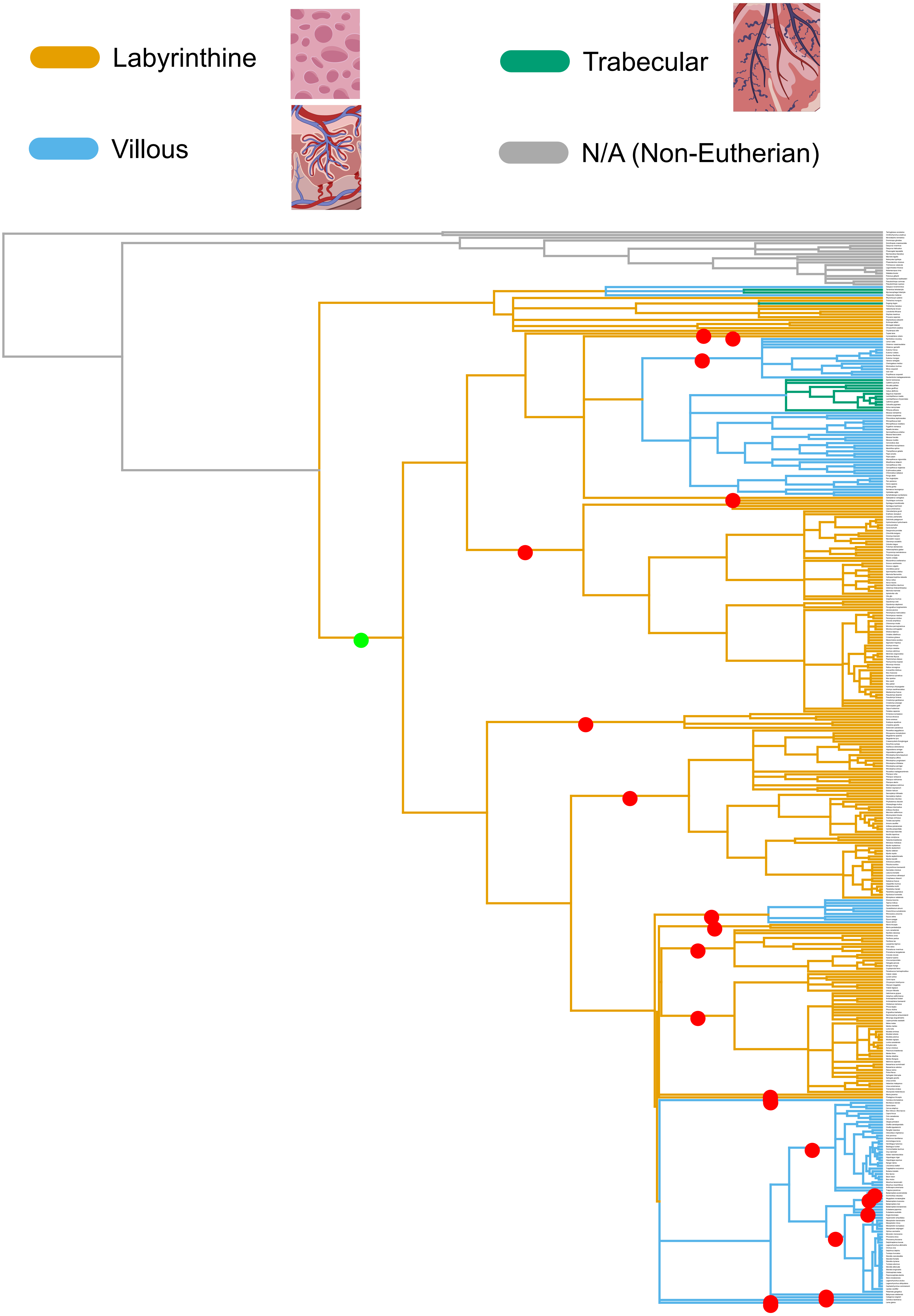

#### MIR590

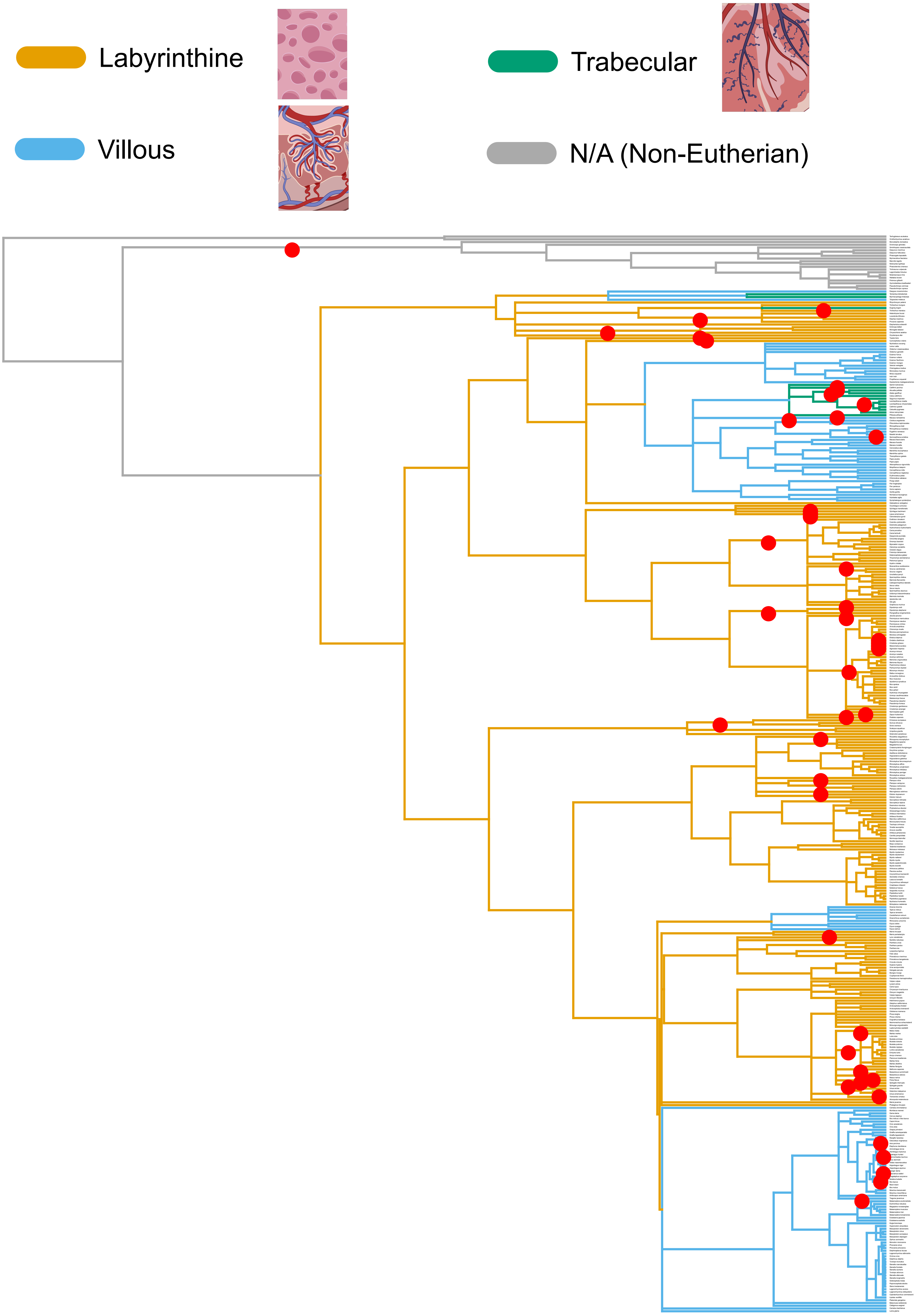

#### MIR887

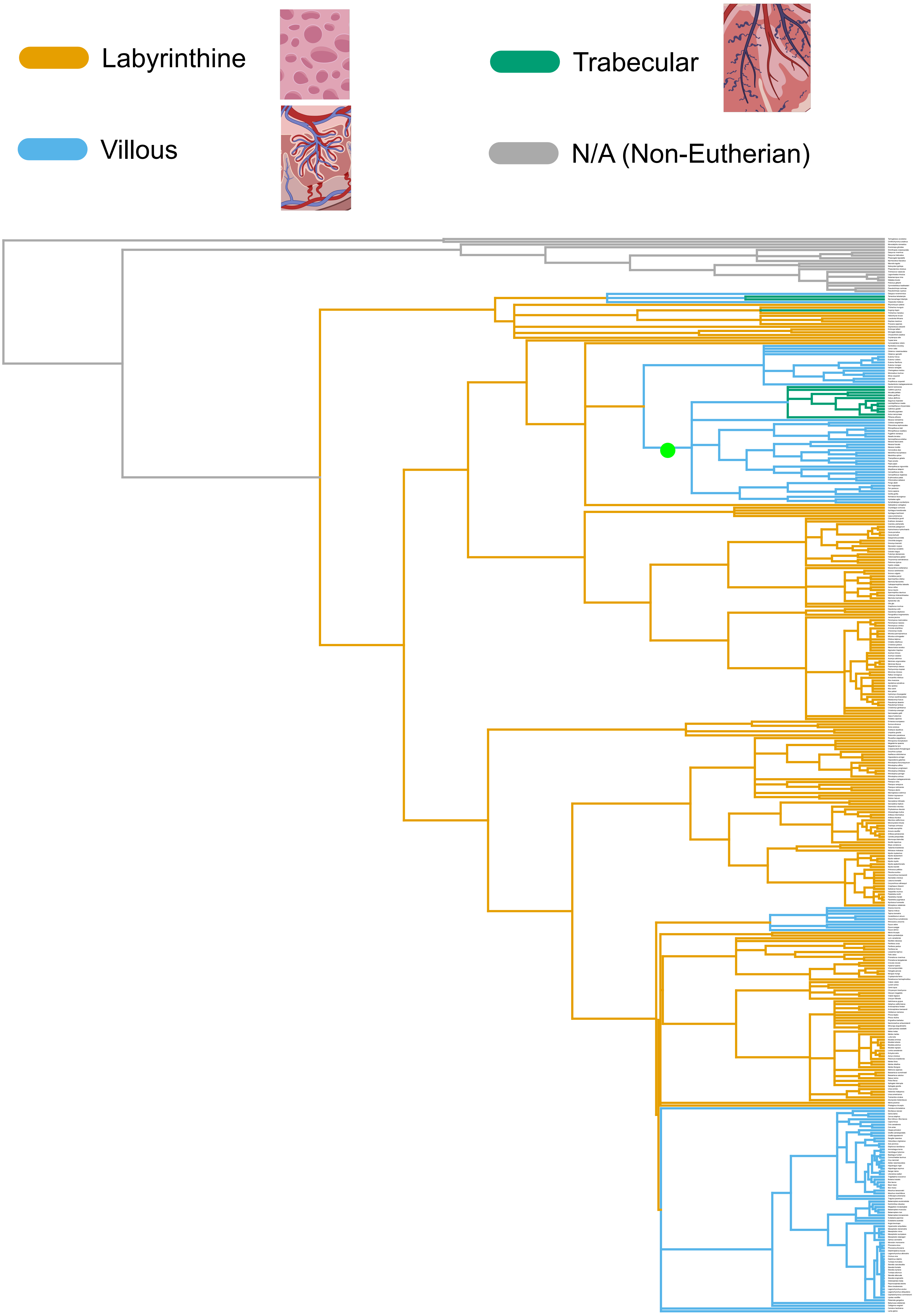

#### MIR1247

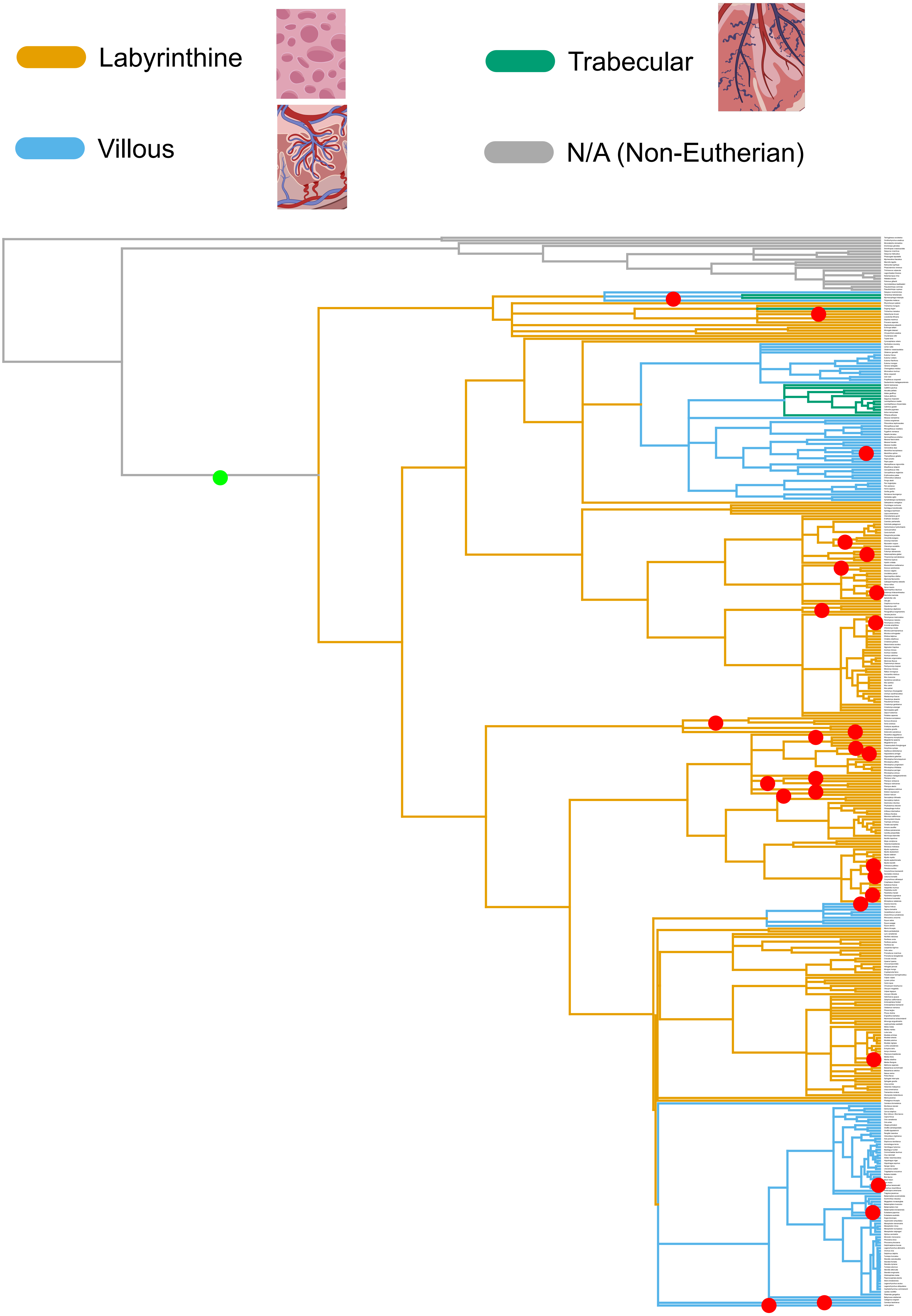

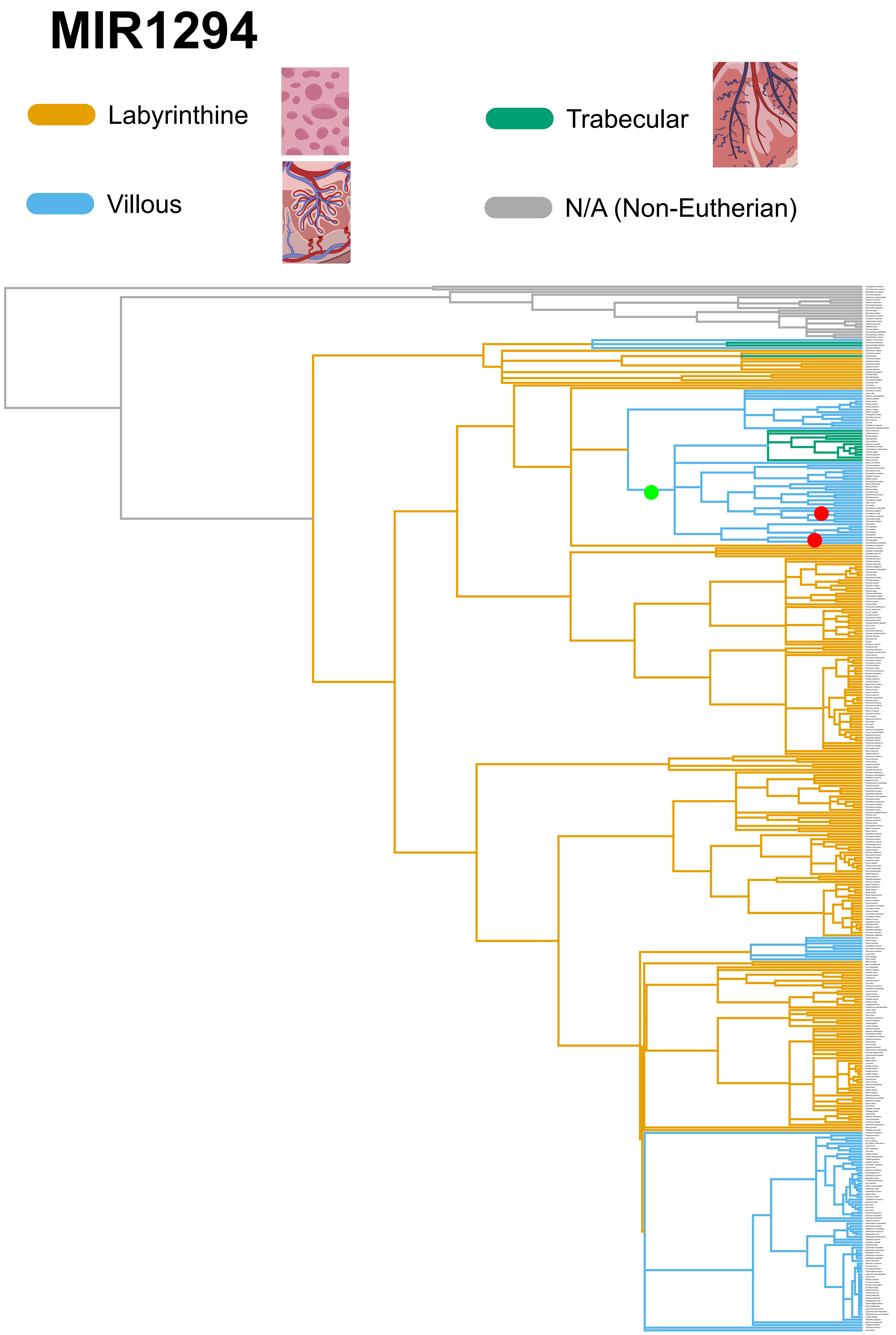

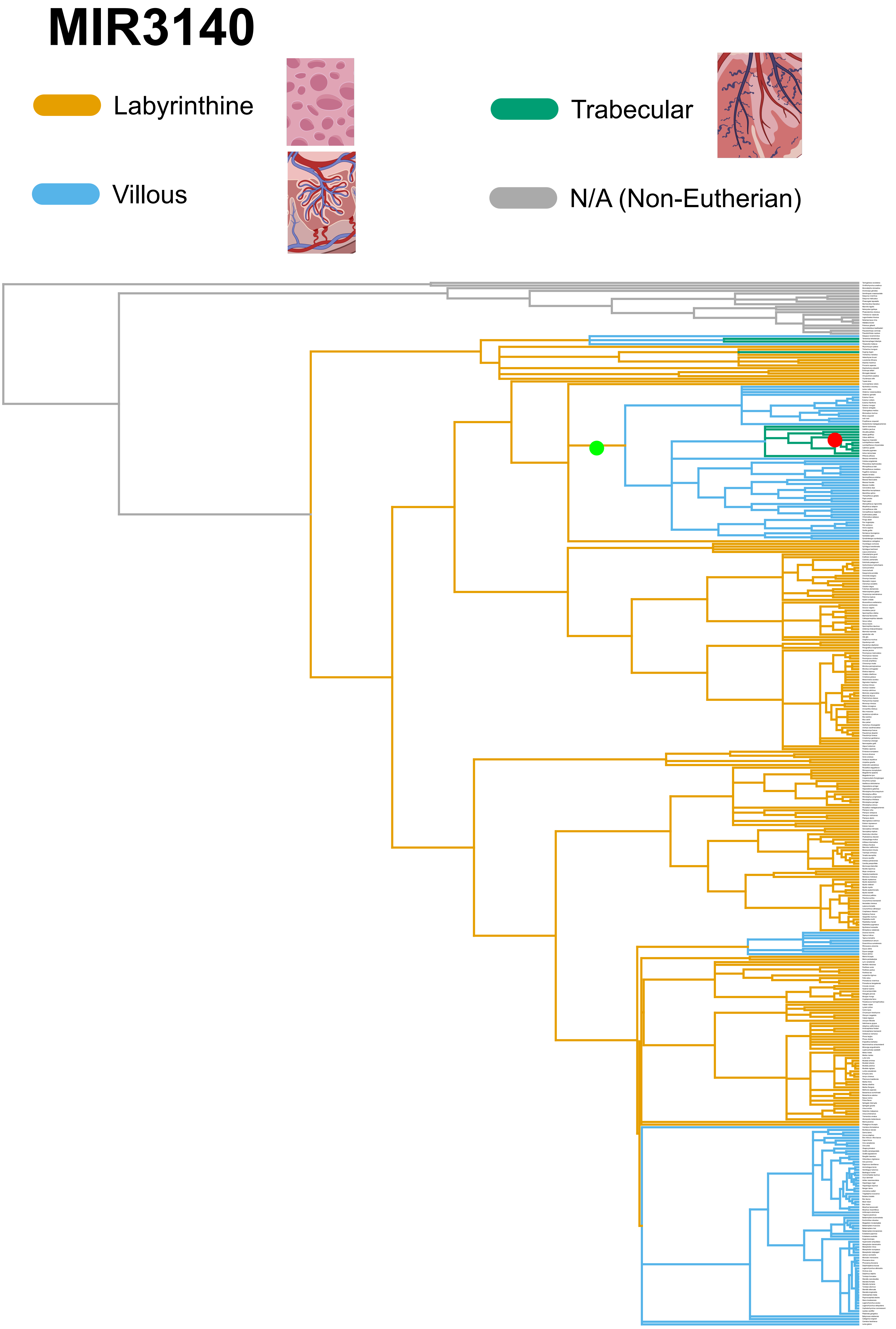

#### MIR3187

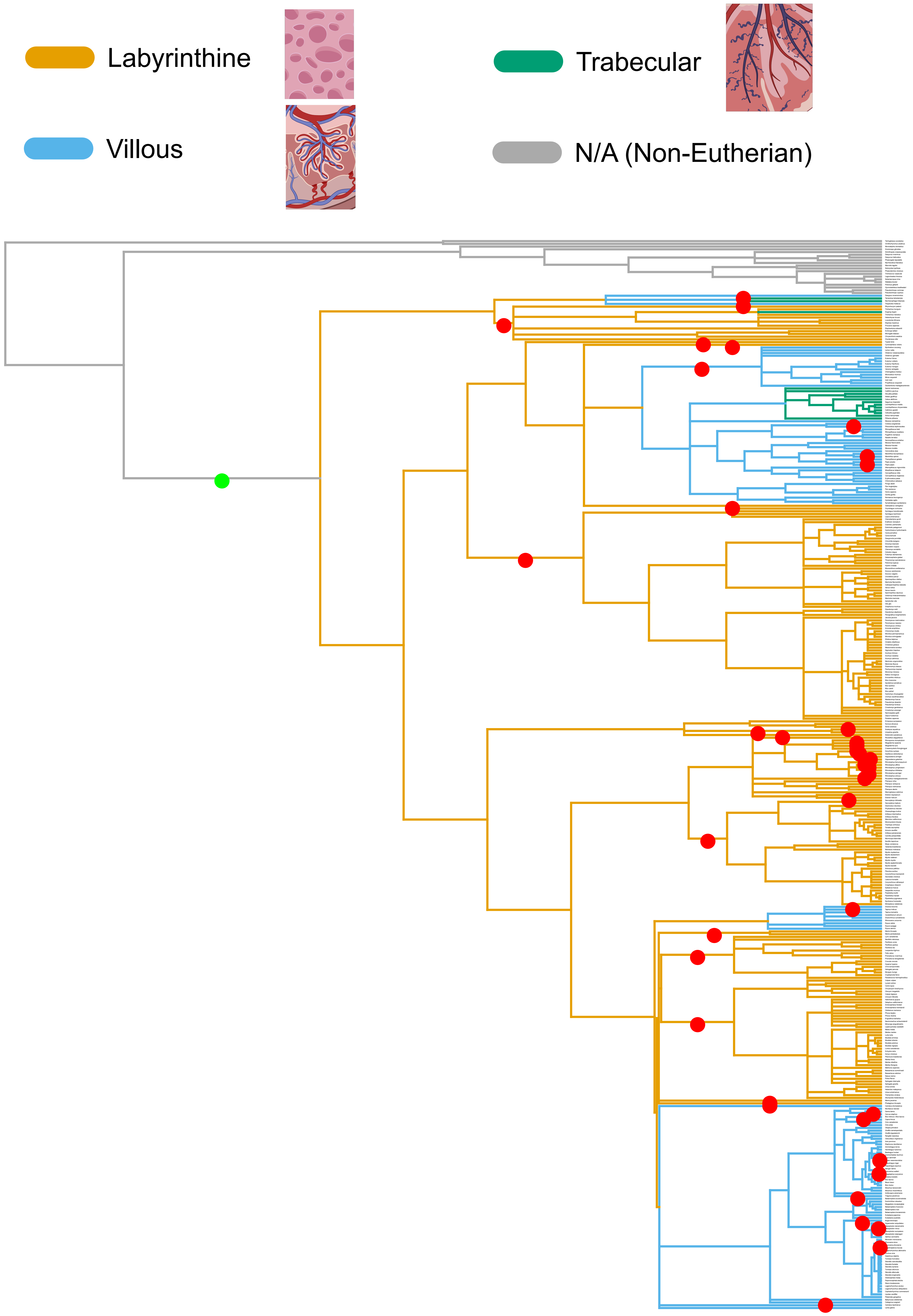

#### MIR5579

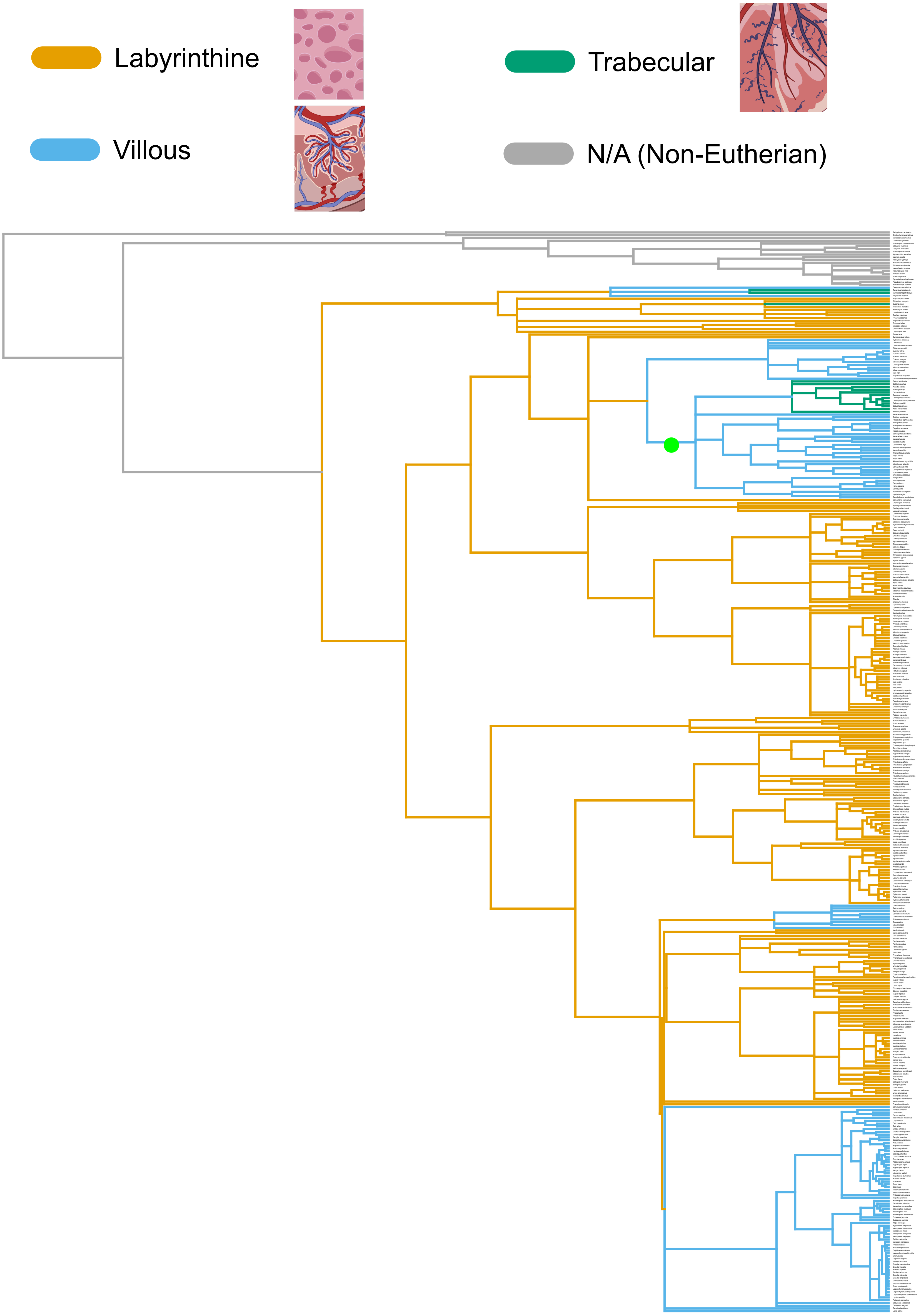

### Figure S14

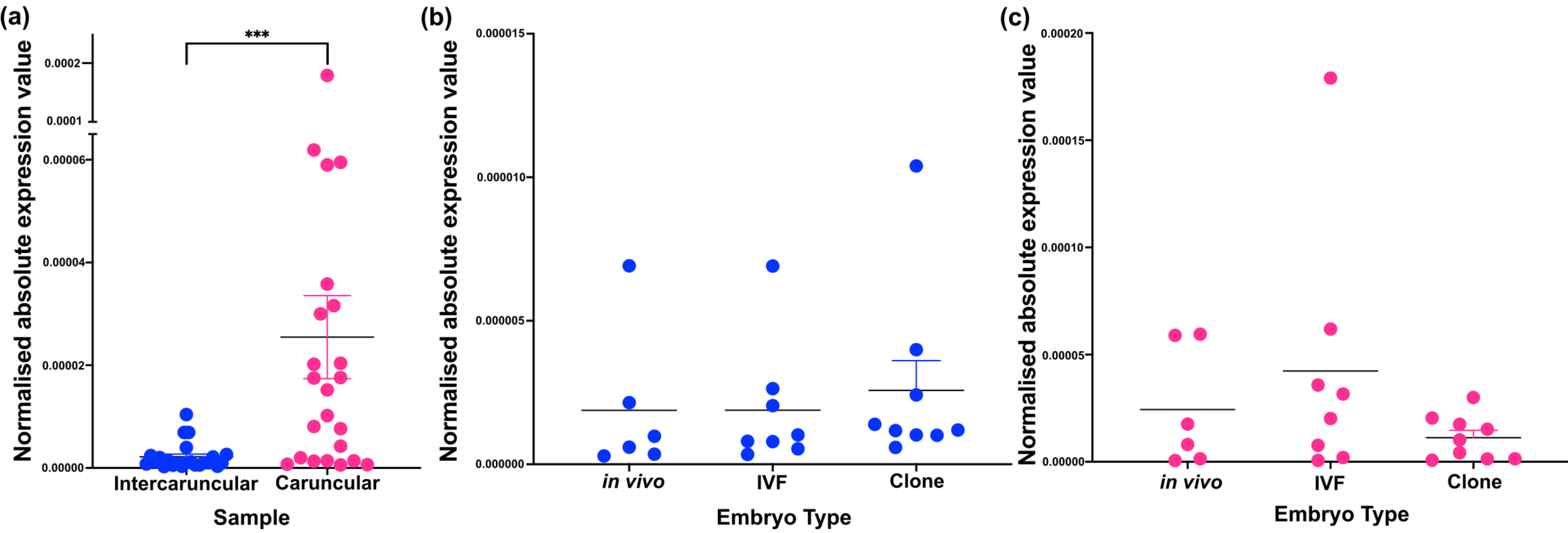

### Figure S15

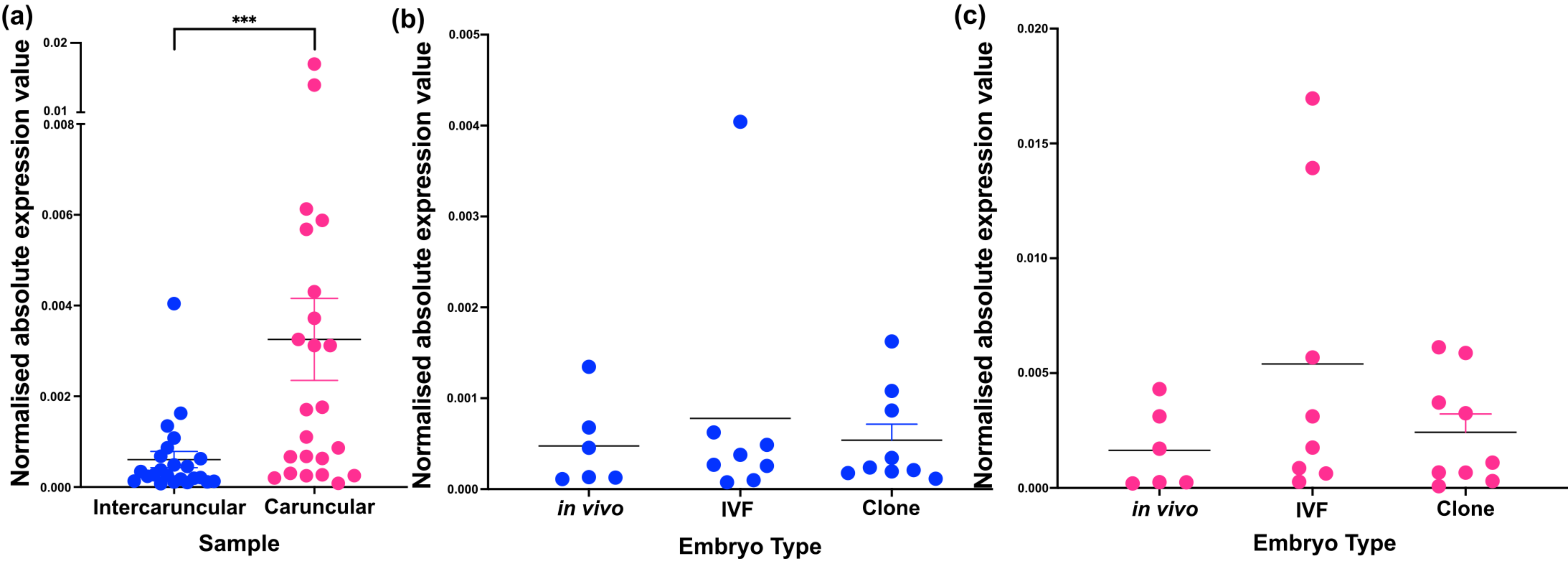

### Figure S16

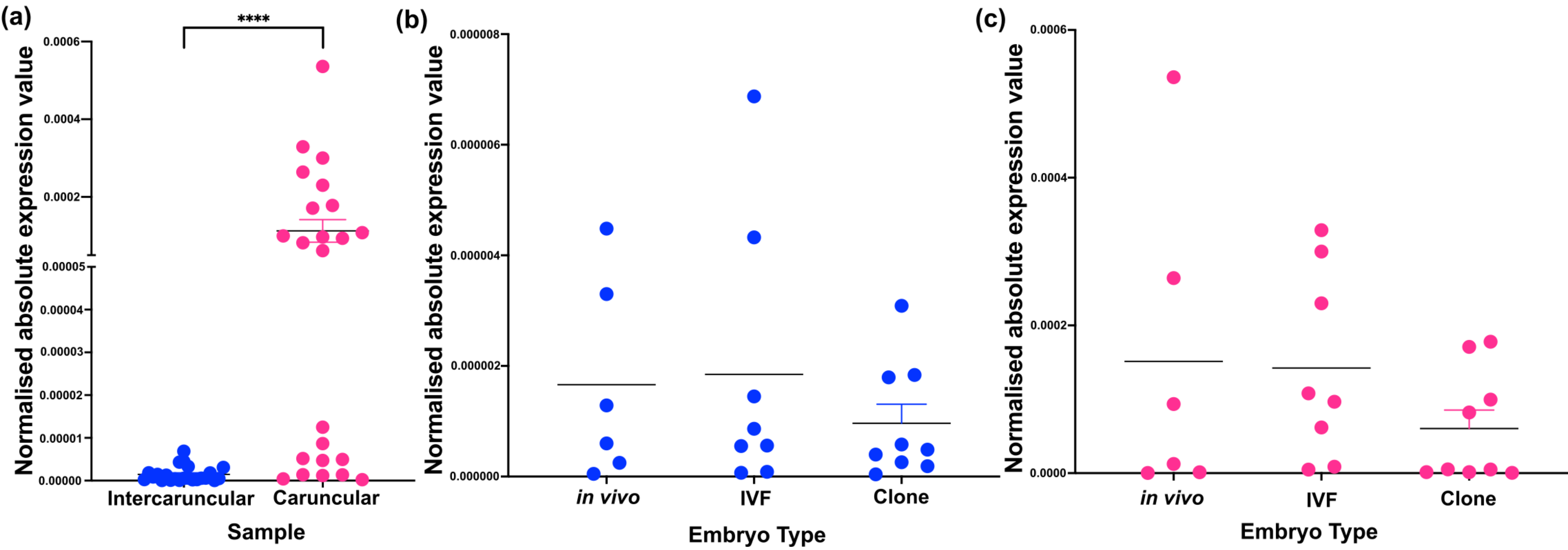
