## Supplementary Legends for "Mammal placental phenotypes are predictable from microRNA repertoires"

**SUPPLEMENTARY INFORMATION**

**Table S1 - Database of mammal genomes considered for usage in miRNA analyses**

Full record of genomes as sourced from the Vertebrates Genome Project, Zoonomia and DNAzoo (as indicated in the ‘Source’ column), prior to quality filtering.

**Table S2 - Database of placental phenotypes from species with matching genomic data**

Columns provide available phenotype data, including the chorial membrane type, venous pattern and placental shape. Columns B-D are phenotype designations as recorded in Wildman et al (2006), while columns E-G indicate the updated phenotype records determined by re-examination of the original literature.

**Table S3 - Branching time estimates use in penalised-likelihood dating of mammal phylogeny for downstream analysis**

Columns ‘age.min’ and ‘age.max’ indicate bounds of age estimates for the given node. ‘soft.bounds’ is an unused dummy column required by the *chronos* dating function. ‘Clade’ indicates the clade represented by the node in question. ‘MRCAs’ provide two species from our genomic dataset used to identify the specific node number of our phylogeny, through identification of their MRCA (i.e. two most distantly-related species within the clade in question’). ‘Source’ denotes the original publication from which the age estimates are taken, with nodes not present in Yu et al (2024) being taken from Alvarez et al (2022). The empty ‘node’ column is dynamically filled with node numbers from the phylogeny during the tree-aging process, by referencing the MRCA column.

**Table S4 -Database of existing datasets of endometrial miRNA expression, from across 7 species.**

Endometrial miRNA expression data collated from cow, human, pig, goat, rat, mouse and horse.

**Table S5 – miRNA presence/absence matrix**

Full matrix of miRNA gene family presence, determined by MirMachine and BLAST detection of genomic miRNA sequences, from 403 mammal species, following genome quality filtering.

**Table S6 – Reconstructed miRNA repertoires of ancestral mammal species**

miRNAs of the mammalian, therian and eutherian ancestors, as indicated by the ‘node of origin’ column. ‘Lost in eutheria’ indicates, for miRNAs originating in the mammalian or therian ancestor, whether that miRNA was subsequently lost in the eutherian ancestor.

**Table S7 – Phenotype-Associated miRNAs**

Database of 42 miRNA families found to be evolutionarily associated with specific placental phenotypes. Values indicate the strength of relationship as determined by Evolink (i.e. Evolink index), with negative values denoting a negative association between miRNA gene presence and phenotype presence. ‘Level_of_support’ denotes whether this relationship is supported by random forest analysis, or by Evolink analysis alone (for order-specific gene-trait associations).

**Table S8 – miRNA Target Prediction**

Targetscan output, detailing predicted gene targets of mammal-exclusive miRNAs from 67 species with available egapx UTR annotations. The column ‘TEC_score’ has been added to the standard Targetscan output to indicate the validity of the interaction as predicted by TEC-miTarget, with a score of >=0.5 deemed reliable.

**Table S9 – Summary of cross-species GO overrepresentation analysis**

For each GO term (column), the proportion of species where an miRNA (row) is present in the genome, in which that GO term is enriched in its targets, is provided as the column value.

**Fig S10 – Membrane-associated miRNA Reconstructions**

Membrane phenotype MCMC reconstructions with gain (green circles) and loss (red circles) of membrane-associated miRNA families mapped on.

**Fig S11 – Placental shape-associated miRNA Reconstructions**

Placental shape phenotype MCMC reconstructions with gain (green circles) and loss (red circles) of membrane-associated miRNA families mapped on.

**Fig S12 – Venous pattern-associated miRNA Reconstructions**

Venous pattern phenotype MCMC reconstructions with gain (green circles) and loss (red circles) of membrane-associated miRNA families mapped on.

**Table S13 - Reduced-Terms Summaries of Key miRNA Gene Targets Across Species**

Table detailing across four worksheets (core eutherian miRNAs, membrane-associated miRNAs, placental shape-associated miRNAs and venous pattern-associated miRNAs) the reduced-terms summaries of, for each group, Gene Ontology terms from overrepresentaiton analysis of genes targeted by miRNAs in that group, weighted by the number of such miRNAs for which the GO term in question appears in the top 100 most conserved across species.

**Fig S14 - Expression levels of bta-miR-11986 in bovine endometrial tissue**

Differences between intercarunculuar and carcuncular tissues (a), with breakdown of expression levels across embryo types in both intercaruncular (b) and caruncular (c) tissue. Endometrial samples were taken on Day 16 of pregnancy. Summary lines in all boxplots represent mean values, error bars represent standard error of the mean. Significant differences in expression where p<0.01 are denoted by ***, and where p<0.001 are denoted by ****.

**Fig S15 - Expression levels of MIR-11968-P1 in bovine endometrial tissue**

Differences between intercarunculuar and carcuncular tissues (a), with breakdown of expression levels across embryo types in both intercaruncular (b) and caruncular (c) tissue. Endometrial samples were taken on Day 16 of pregnancy. Summary lines in all boxplots represent mean values, error bars represent standard error of the mean. Significant differences in expression where p<0.01 are denoted by ***, and where p<0.001 are denoted by ****.

**Fig S16 - Expression levels of MIR-11968-P2 in bovine endometrial tissue**

Differences between intercarunculuar and carcuncular tissues (a), with breakdown of expression levels across embryo types in both intercaruncular (b) and caruncular (c) tissue. Endometrial samples were taken on Day 16 of pregnancy. Summary lines in all boxplots represent mean values, error bars represent standard error of the mean. Significant differences in expression where p<0.01 are denoted by ***, and where p<0.001 are denoted by ****.
