## Supplementary material for "Mammal placental phenotypes are predictable from microRNA repertoires": Figure S11

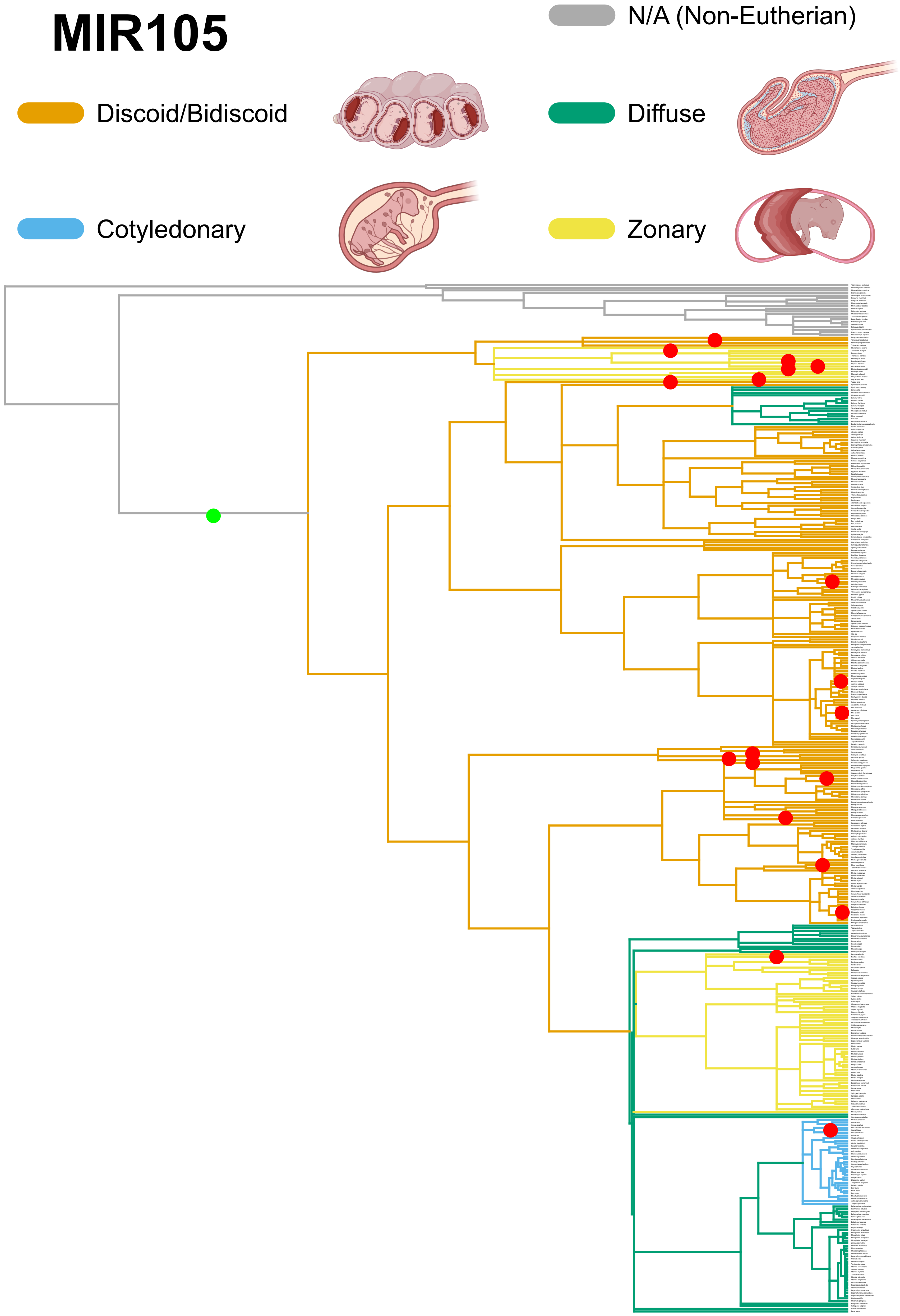
MIR105
N/A (Non-Eutherian)
Discoid/Bidiscoid
Diffuse
Cotyledonary
Zonary

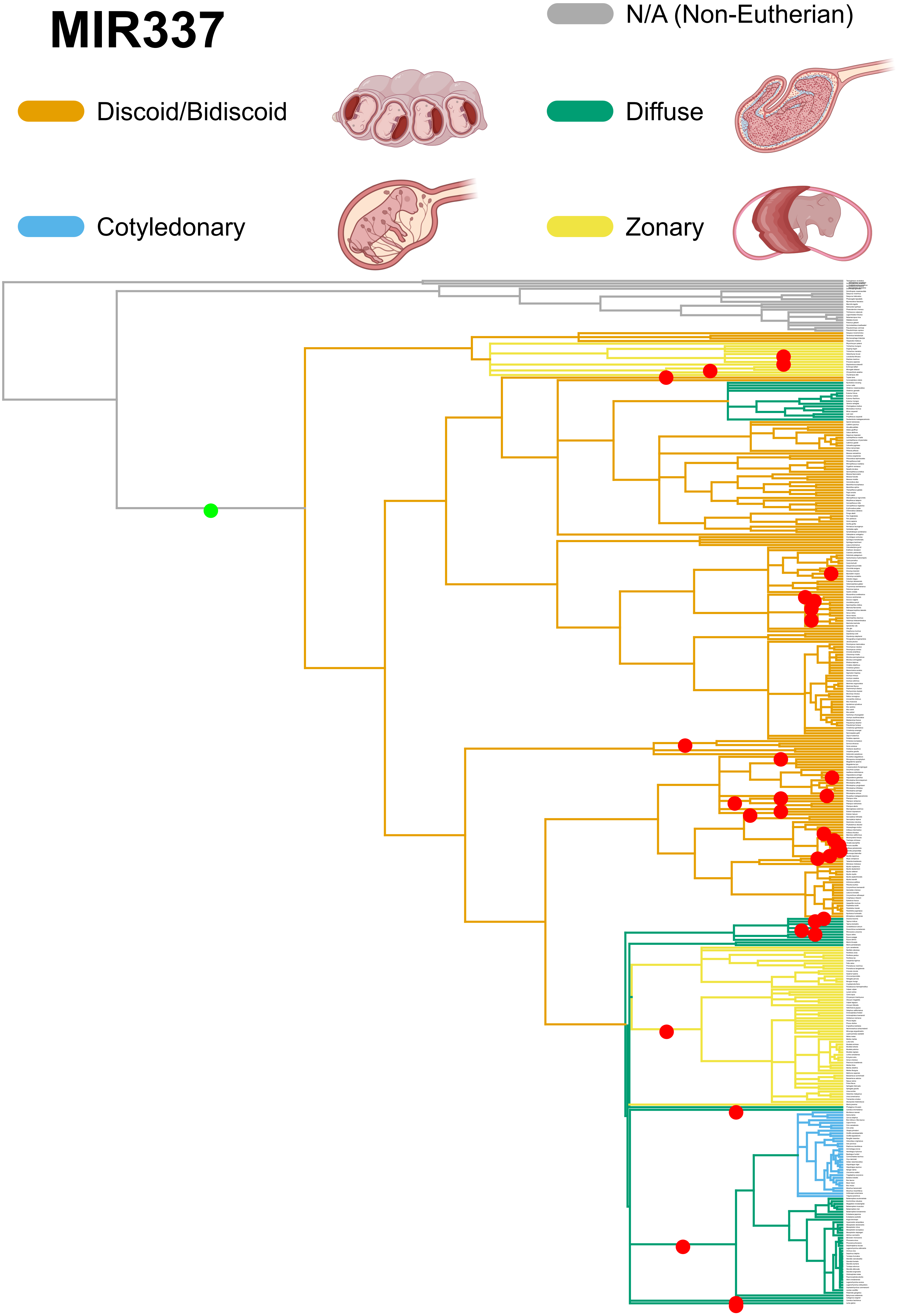
MIR337
N/A (Non-Eutherian)
Discoid/Bidiscoid
Diffuse
Cotyledonary
Zonary

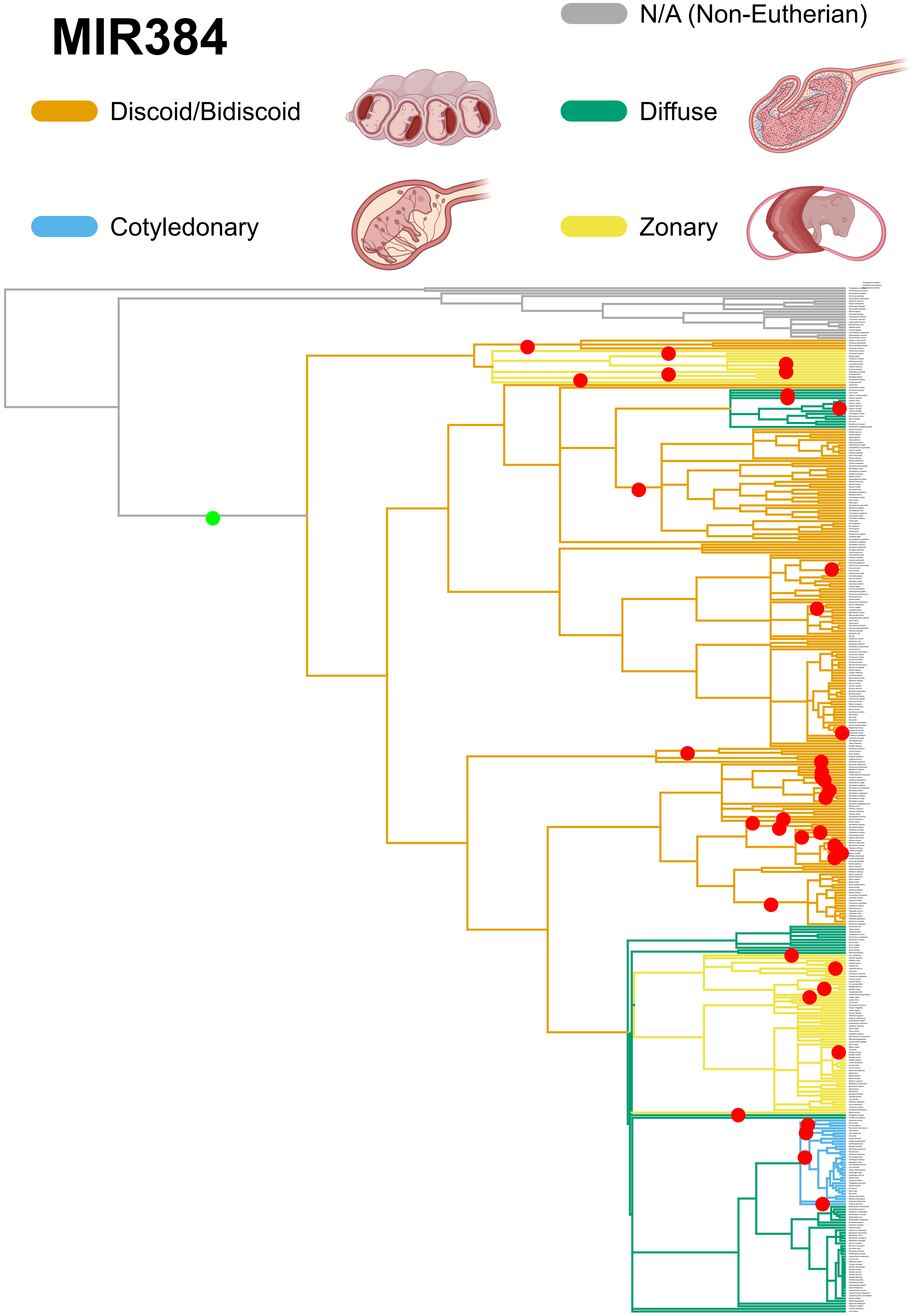
MIR384
N/A (Non-Eutherian)
Discoid/Bidiscoid
Diffuse
Cotyledonary
Zonary

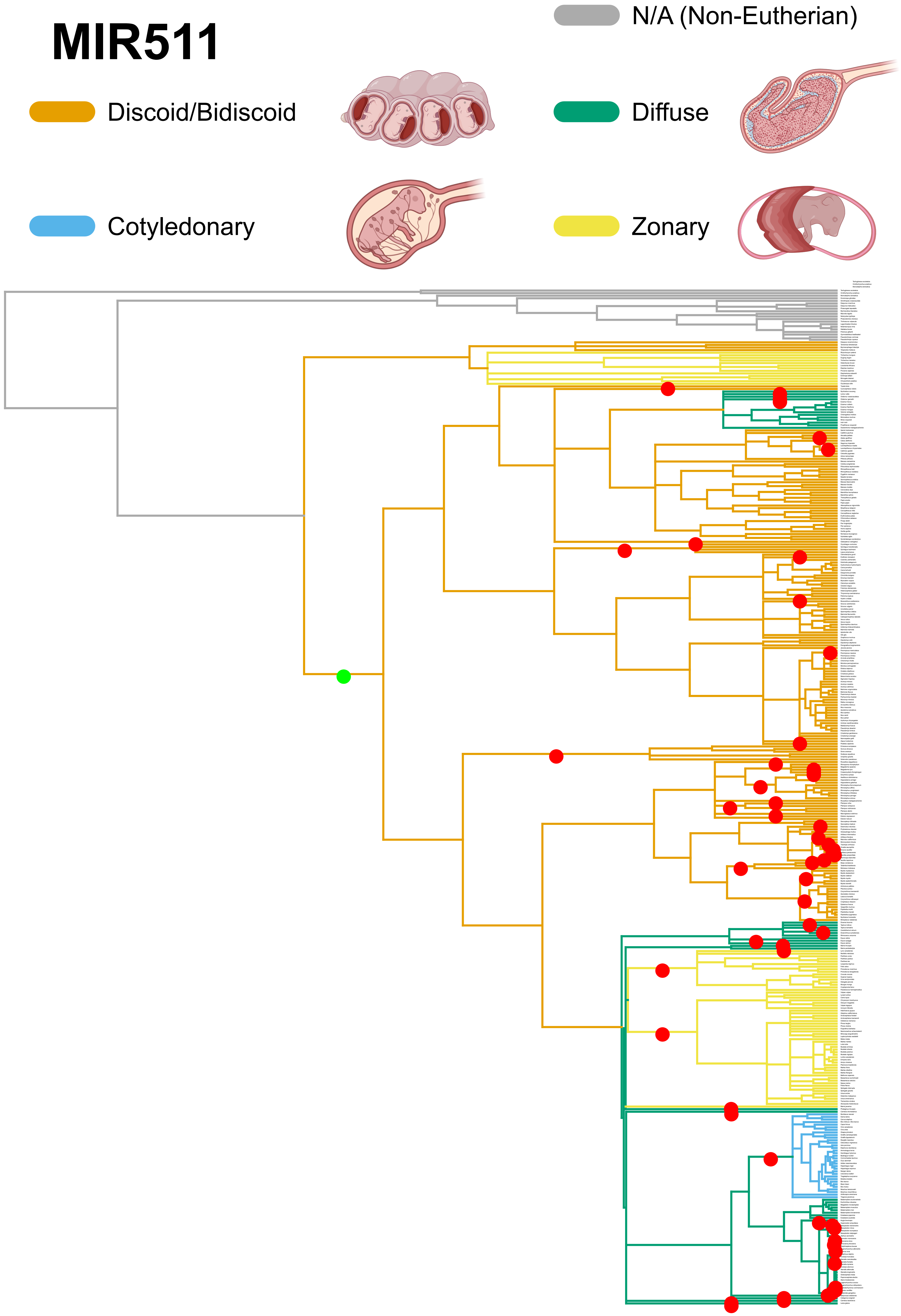
MIR511
N/A (Non-Eutherian)
Discoid/Bidiscoid
Diffuse
Cotyledonary
Zonary

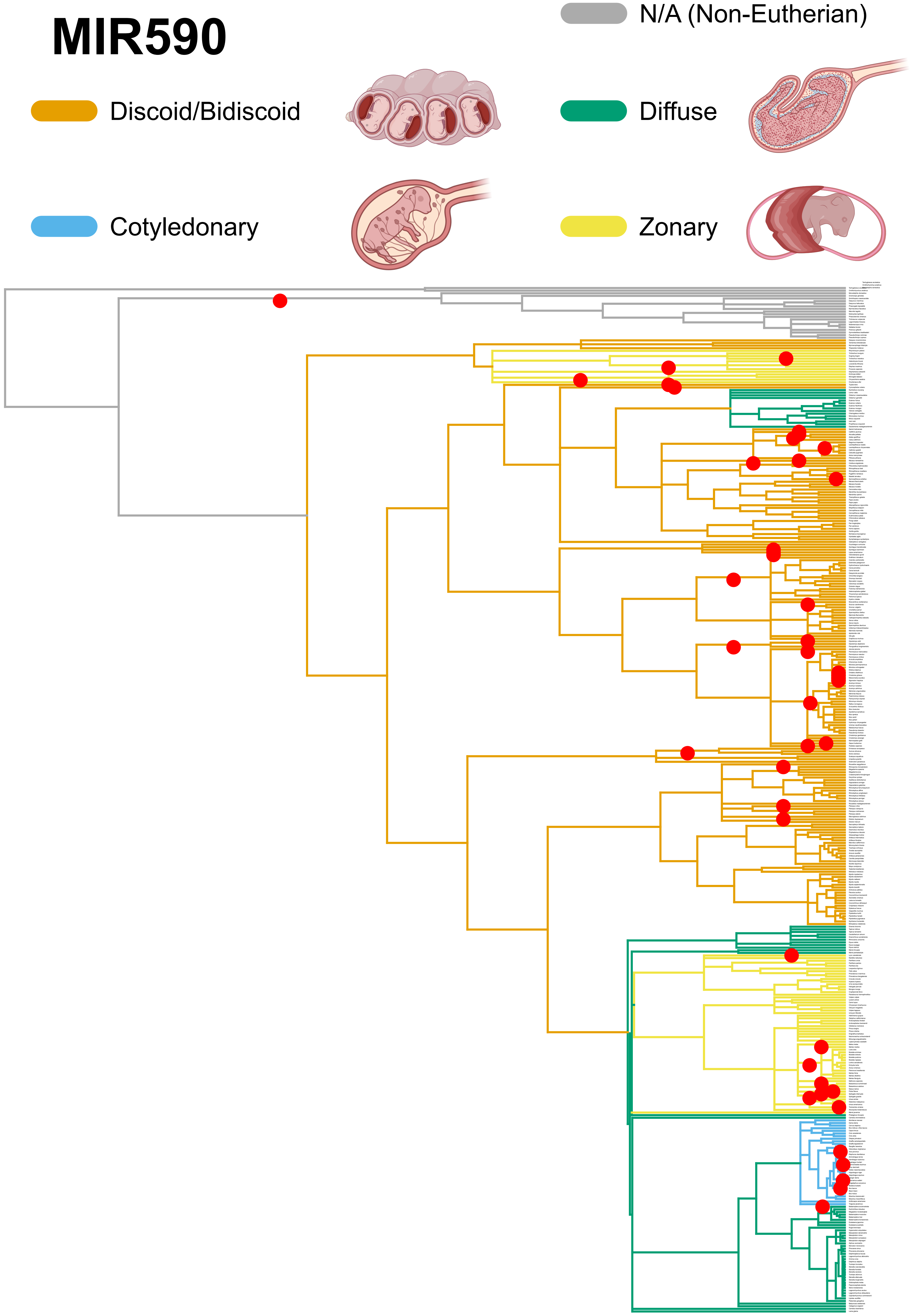
MIR590
N/A (Non-Eutherian)
Discoid/Bidiscoid
Diffuse
Cotyledonary
Zonary

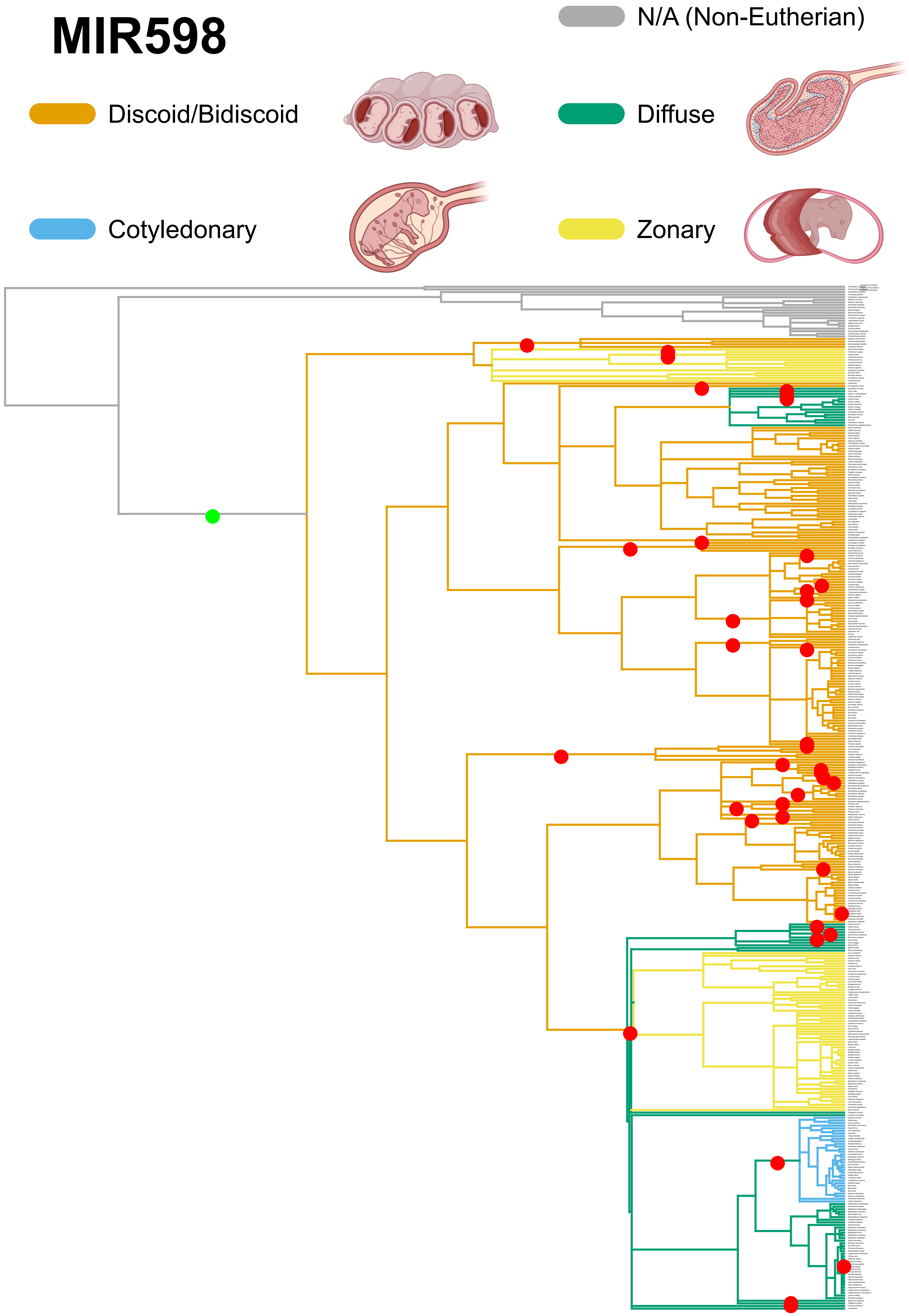
MIR598
N/A (Non-Eutherian)
Discoid/Bidiscoid
Diffuse
Cotyledonary
Zonary

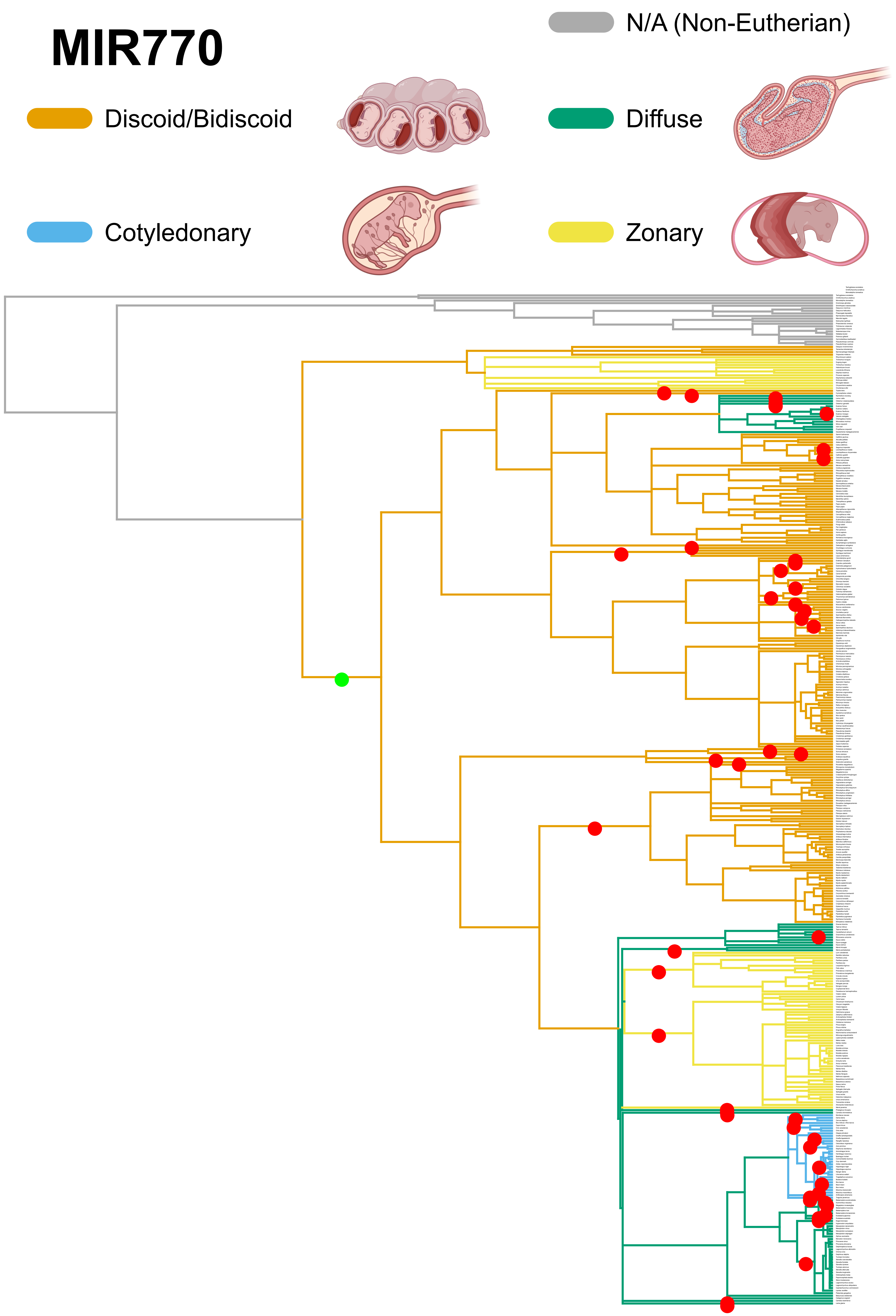
MIR770
N/A (Non-Eutherian)
Discoid/Bidiscoid
Diffuse
Cotyledonary
Zonary

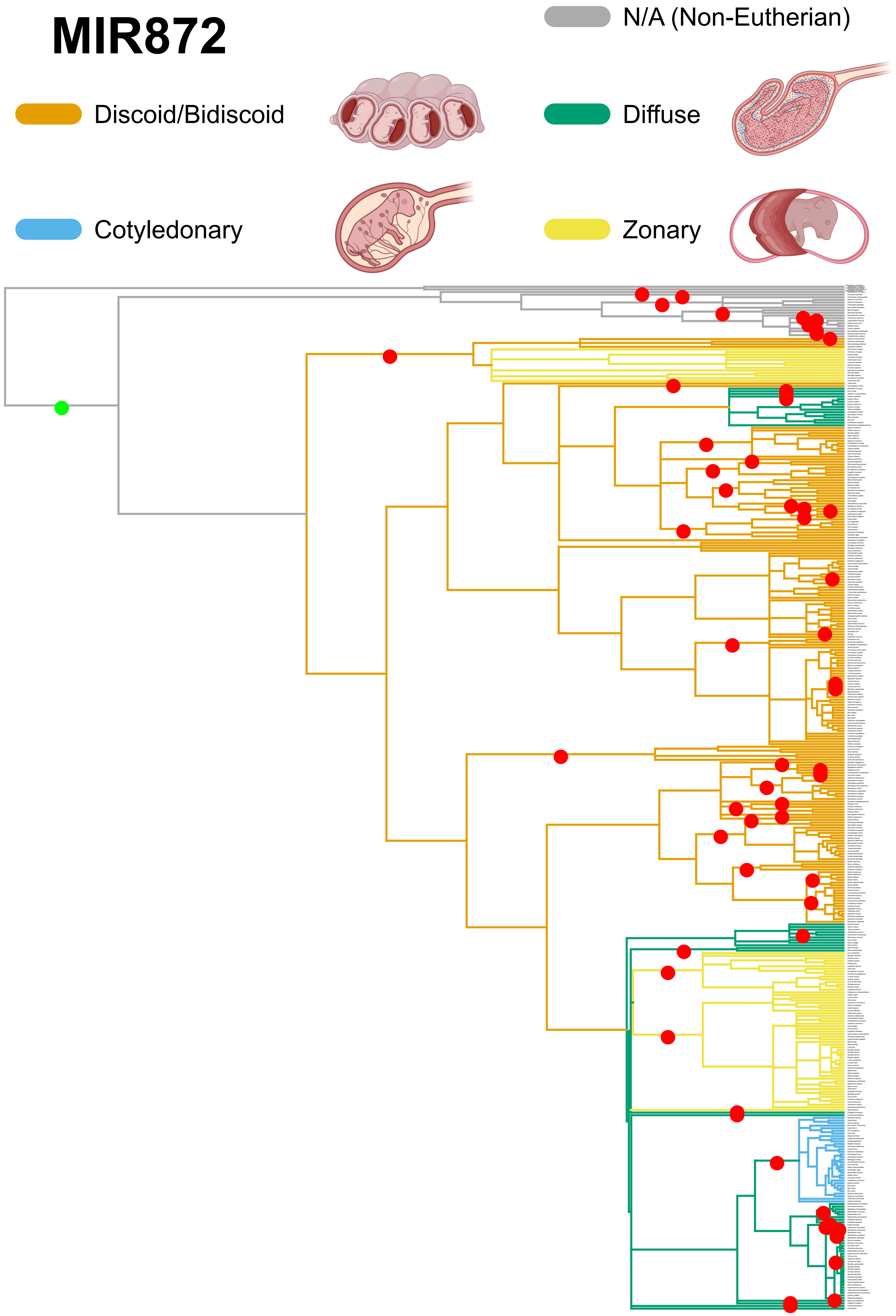
MIR872
N/A (Non-Eutherian)
Discoid/Bidiscoid
Diffuse
Cotyledonary
Zonary

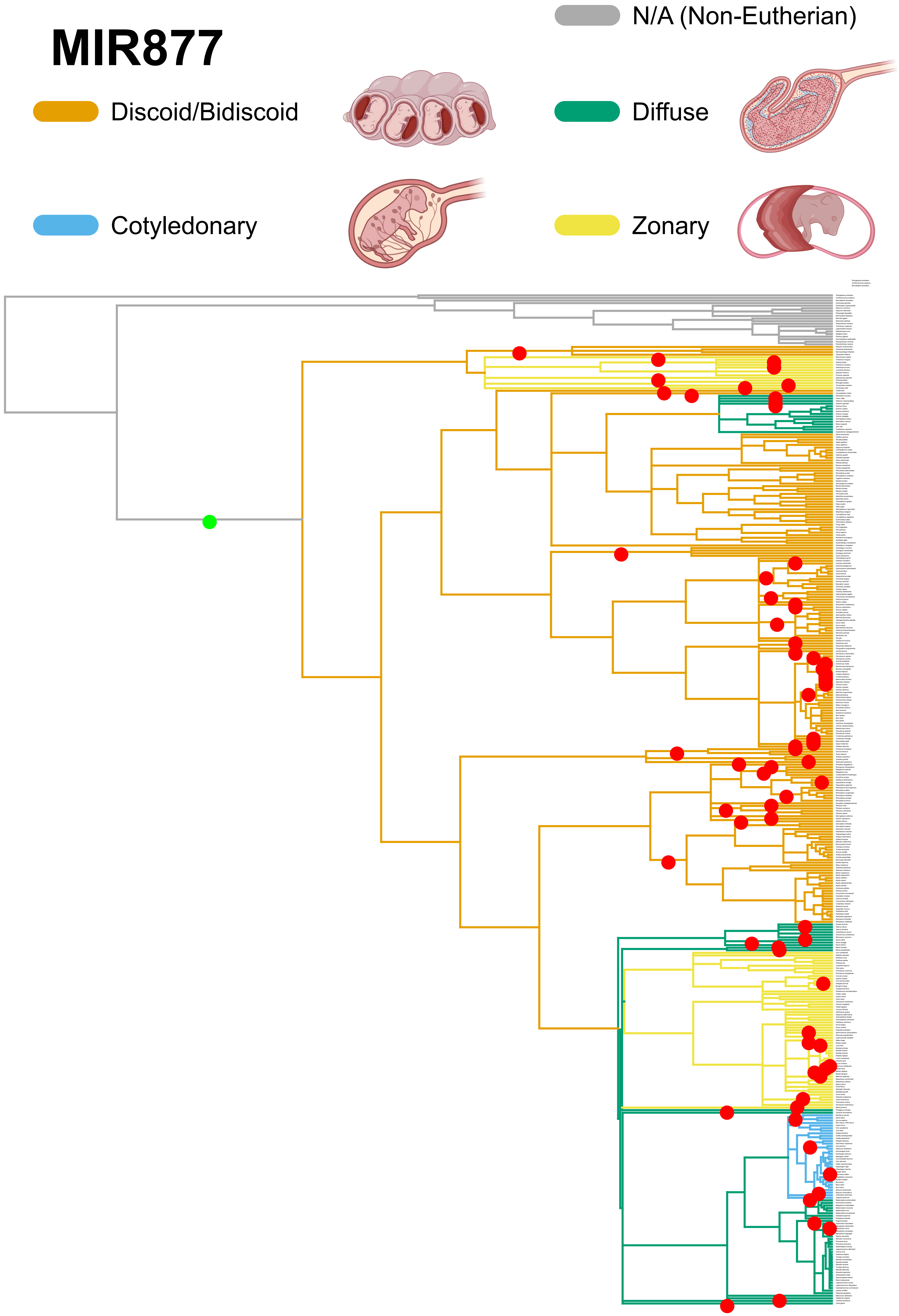

MIR877
N/A (Non-Eutherian)
Discoid/Bidiscoid
Diffuse
Cotyledonary
Zonary

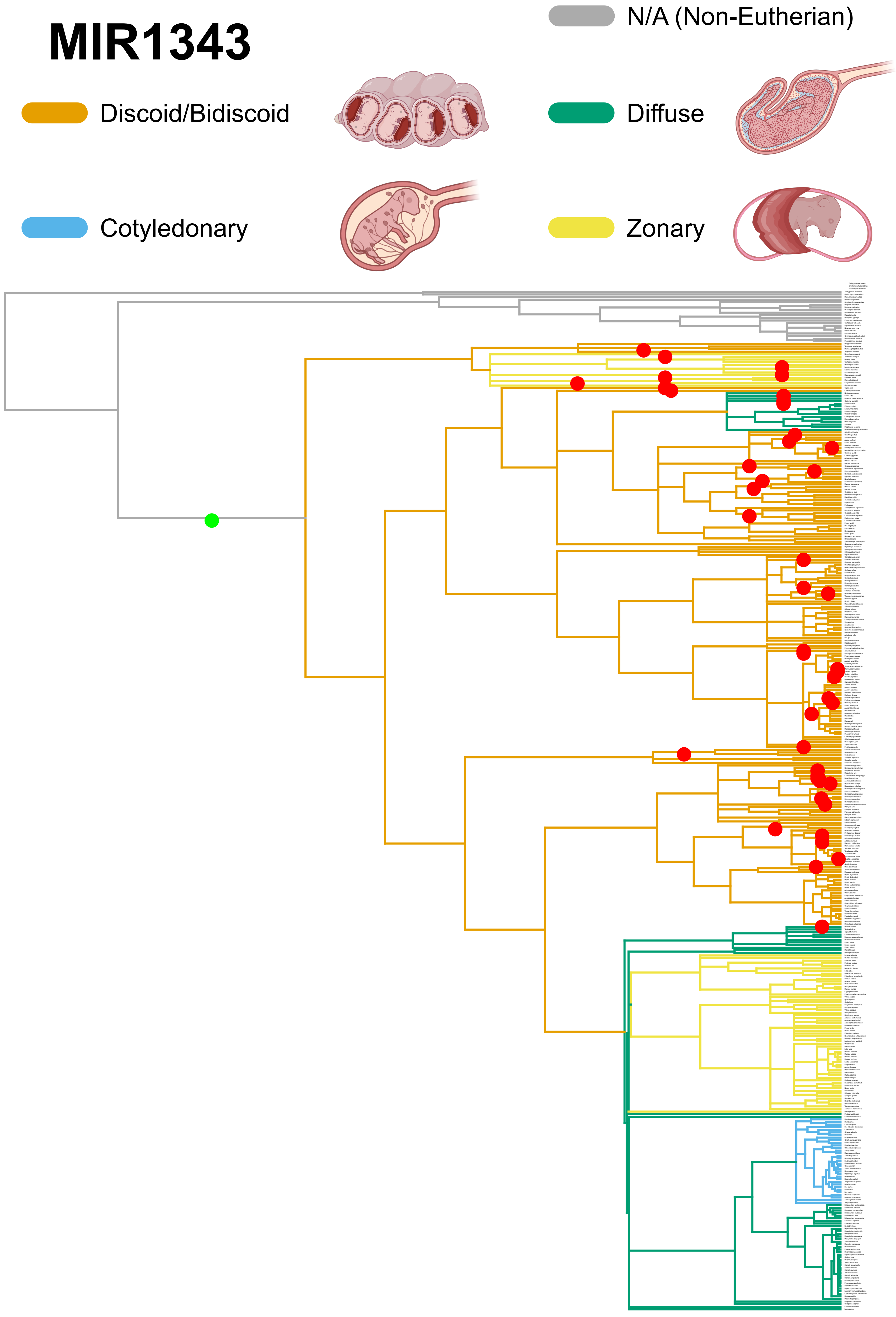
MIR1343
N/A (Non-Eutherian)
Discoid/Bidiscoid
Diffuse
Cotyledonary
Zonary

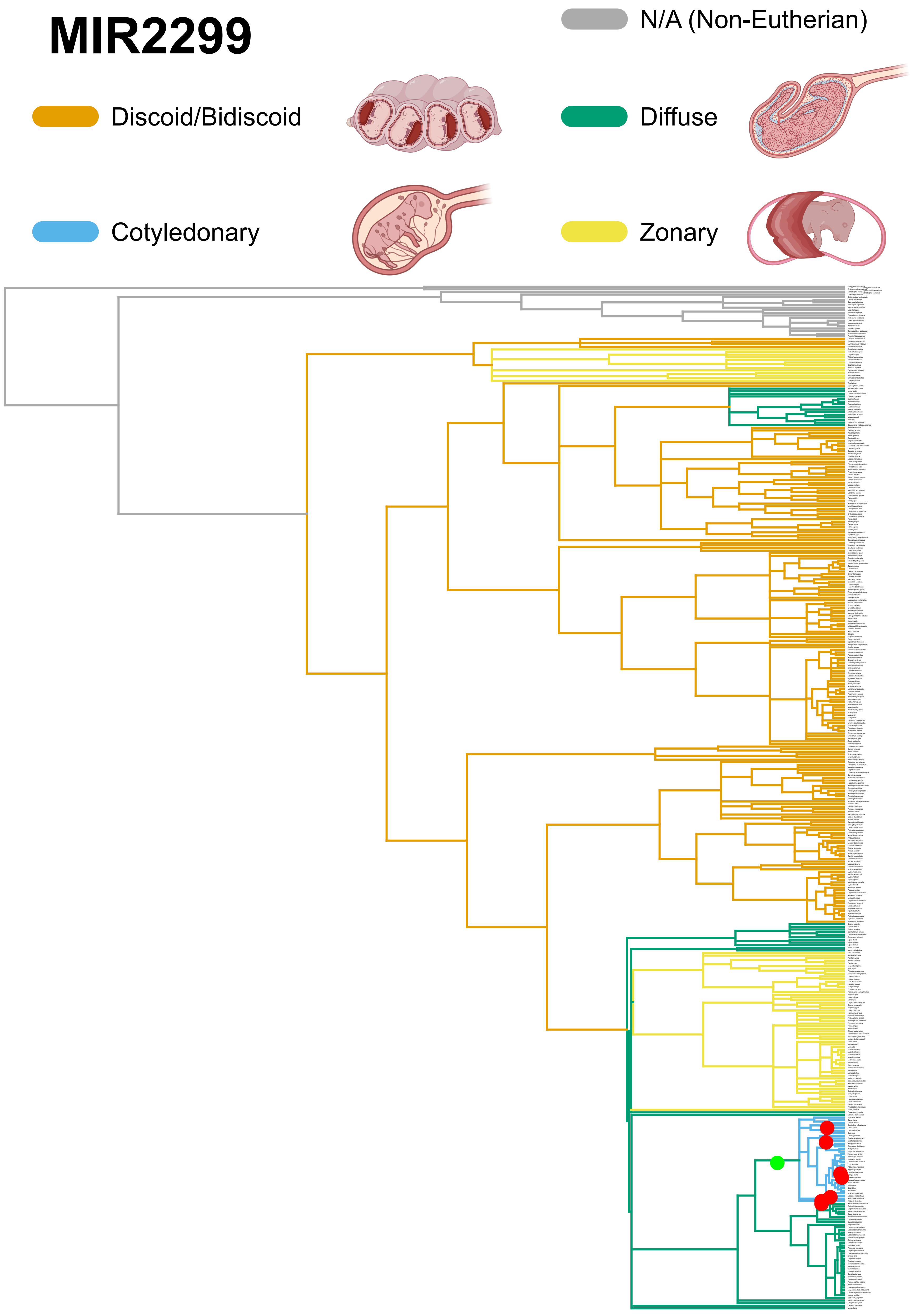
MIR2299
N/A (Non-Eutherian)
Discoid/Bidiscoid
Diffuse
Cotyledonary
Zonary

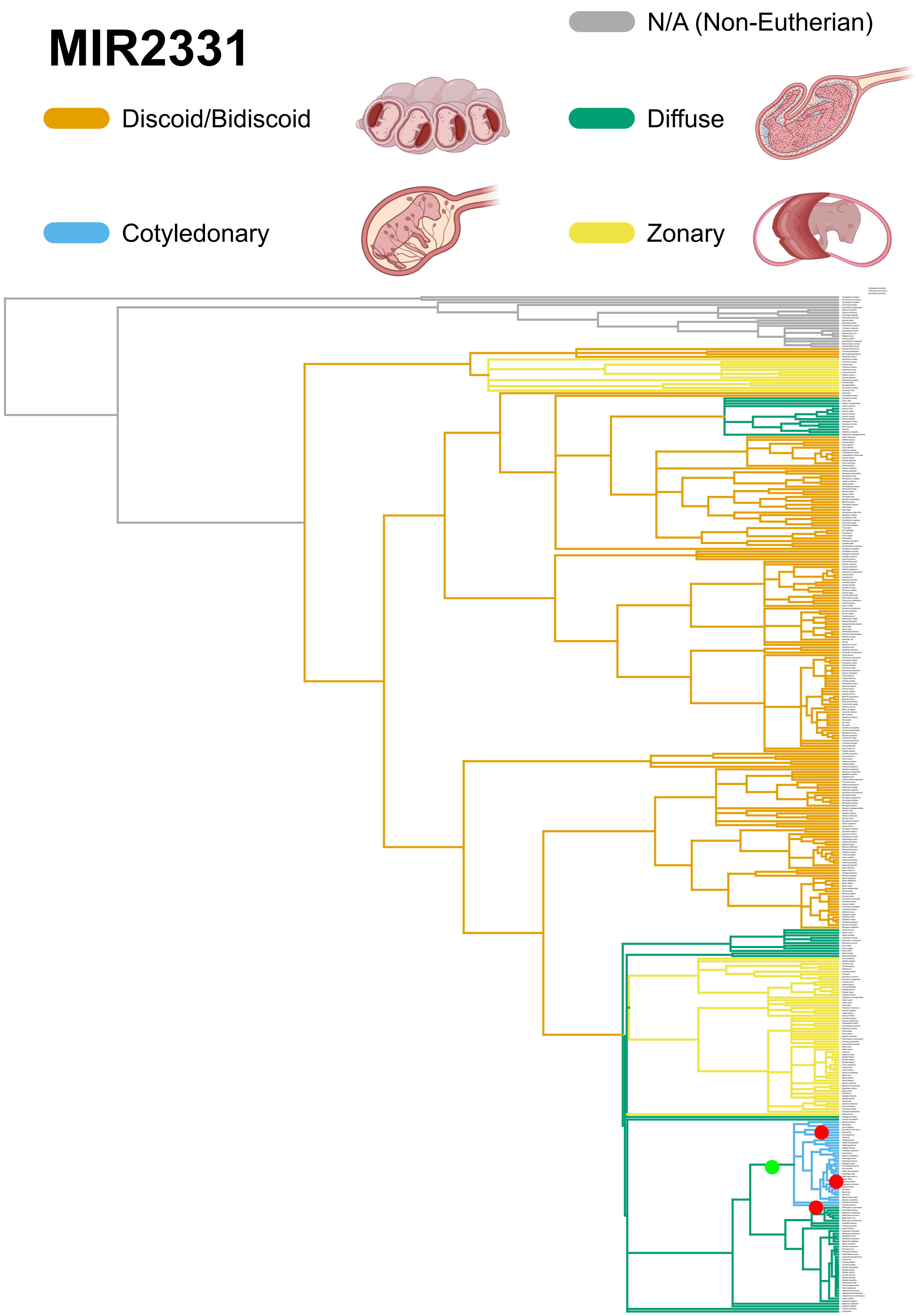
MIR2331
N/A (Non-Eutherian)
Discoid/Bidiscoid
Diffuse
Cotyledonary
Zonary

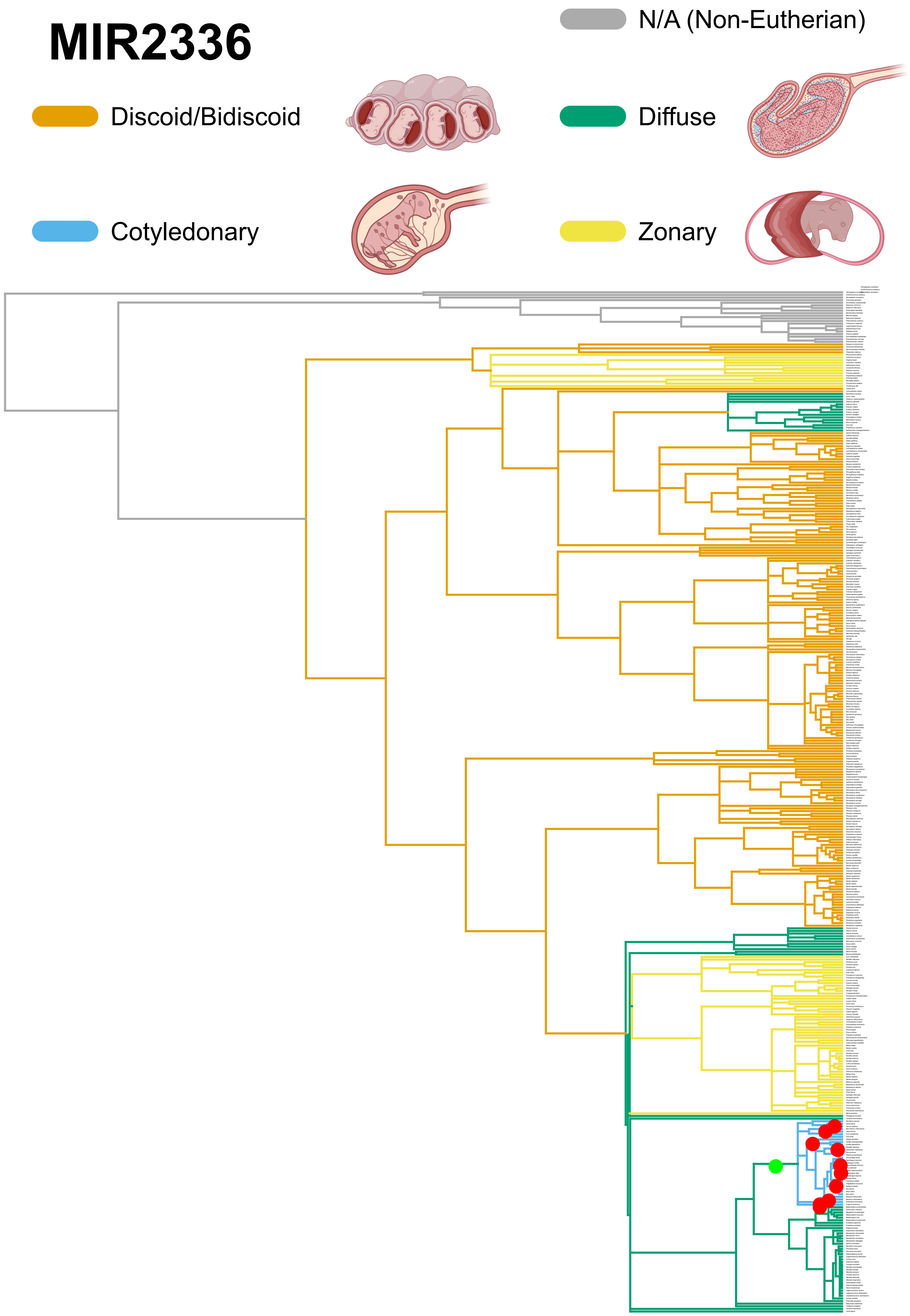
MIR2336
N/A (Non-Eutherian)
Discoid/Bidiscoid
Diffuse
Cotyledonary
Zonary

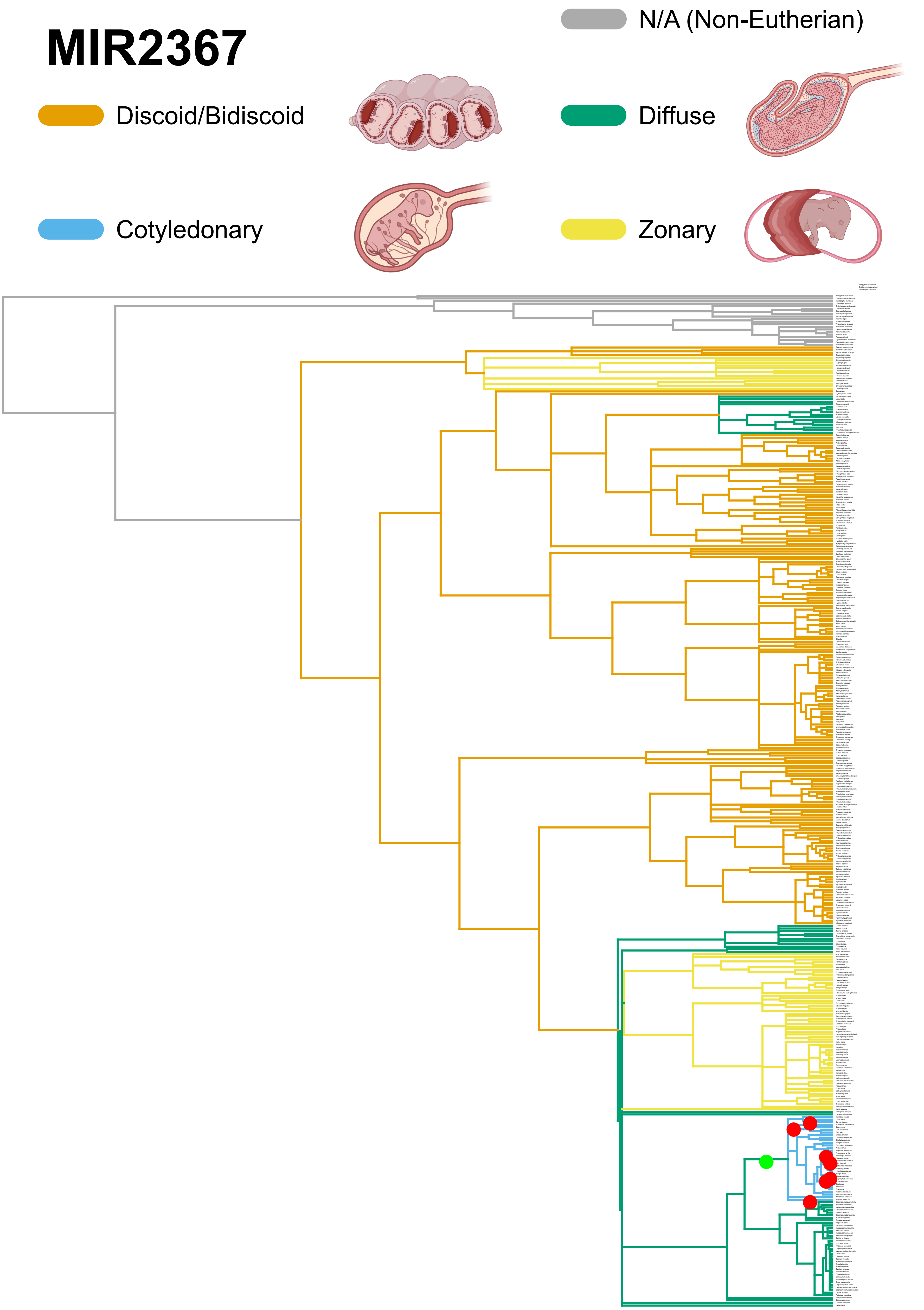
MIR2367
N/A (Non-Eutherian)
Discoid/Bidiscoid
Diffuse
Cotyledonary
Zonary

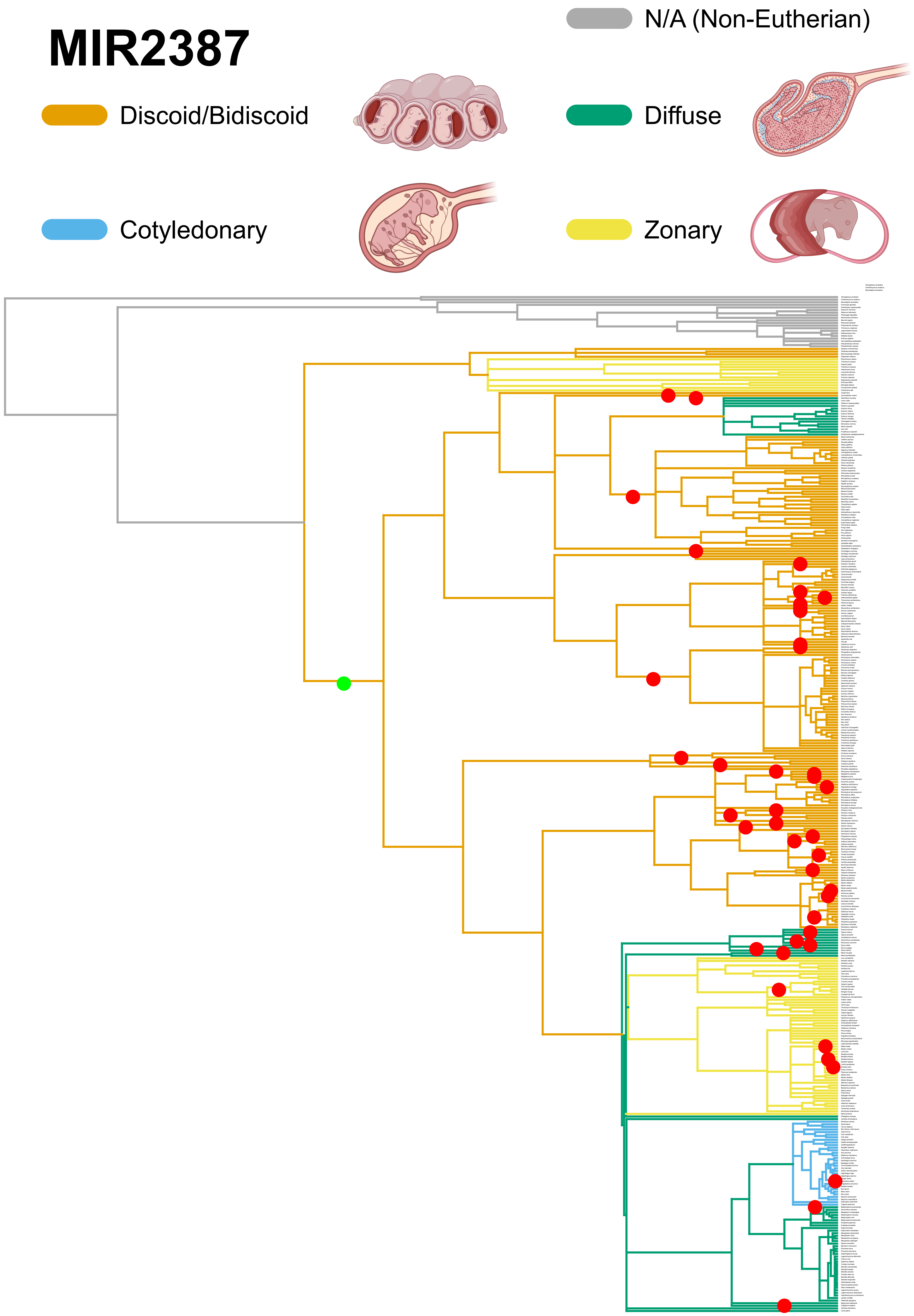
MIR2387
N/A (Non-Eutherian)
Discoid/Bidiscoid
Diffuse
Cotyledonary
Zonary

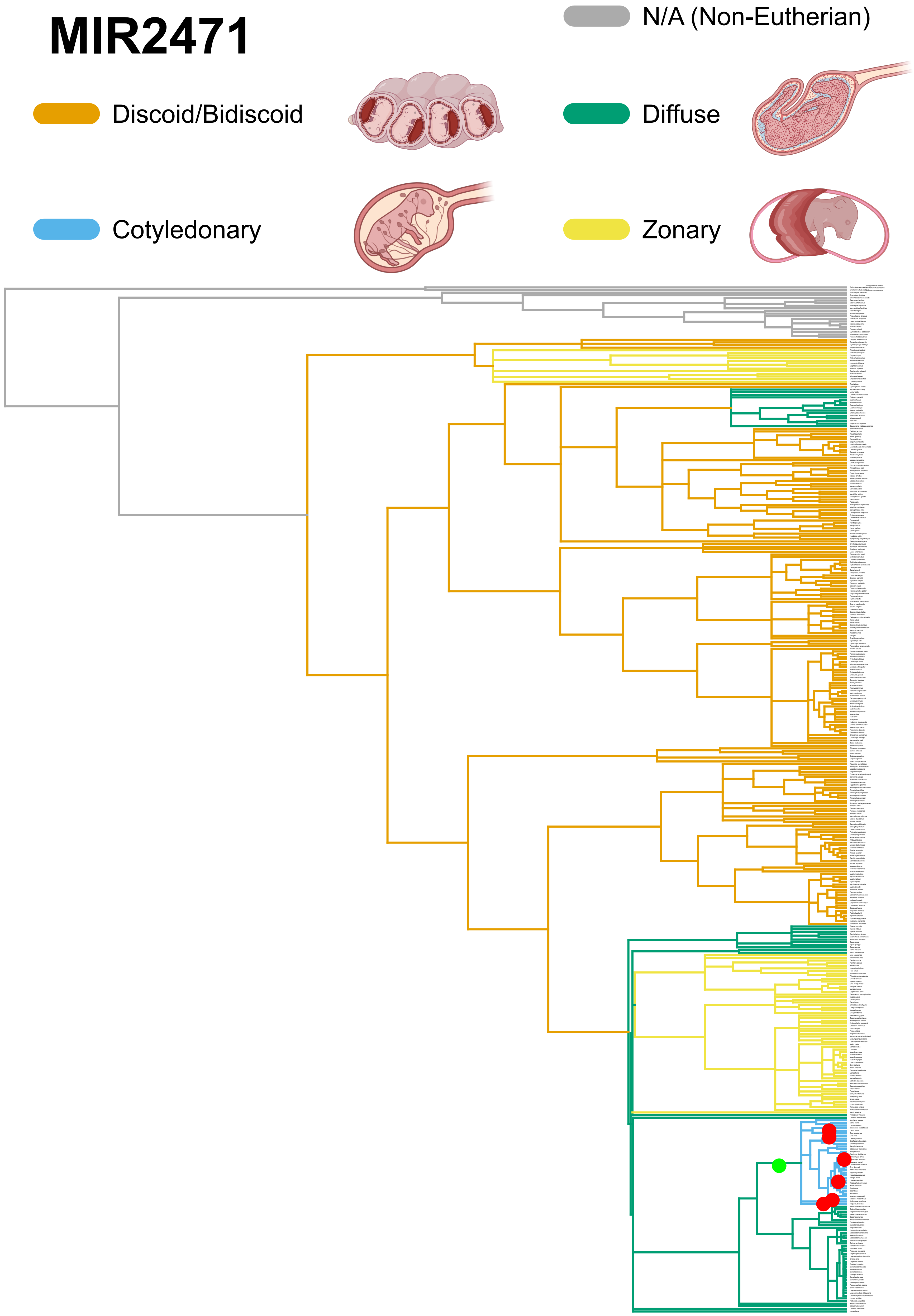

MIR2471
N/A (Non-Eutherian)
Discoid/Bidiscoid
Diffuse
Cotyledonary
Zonary

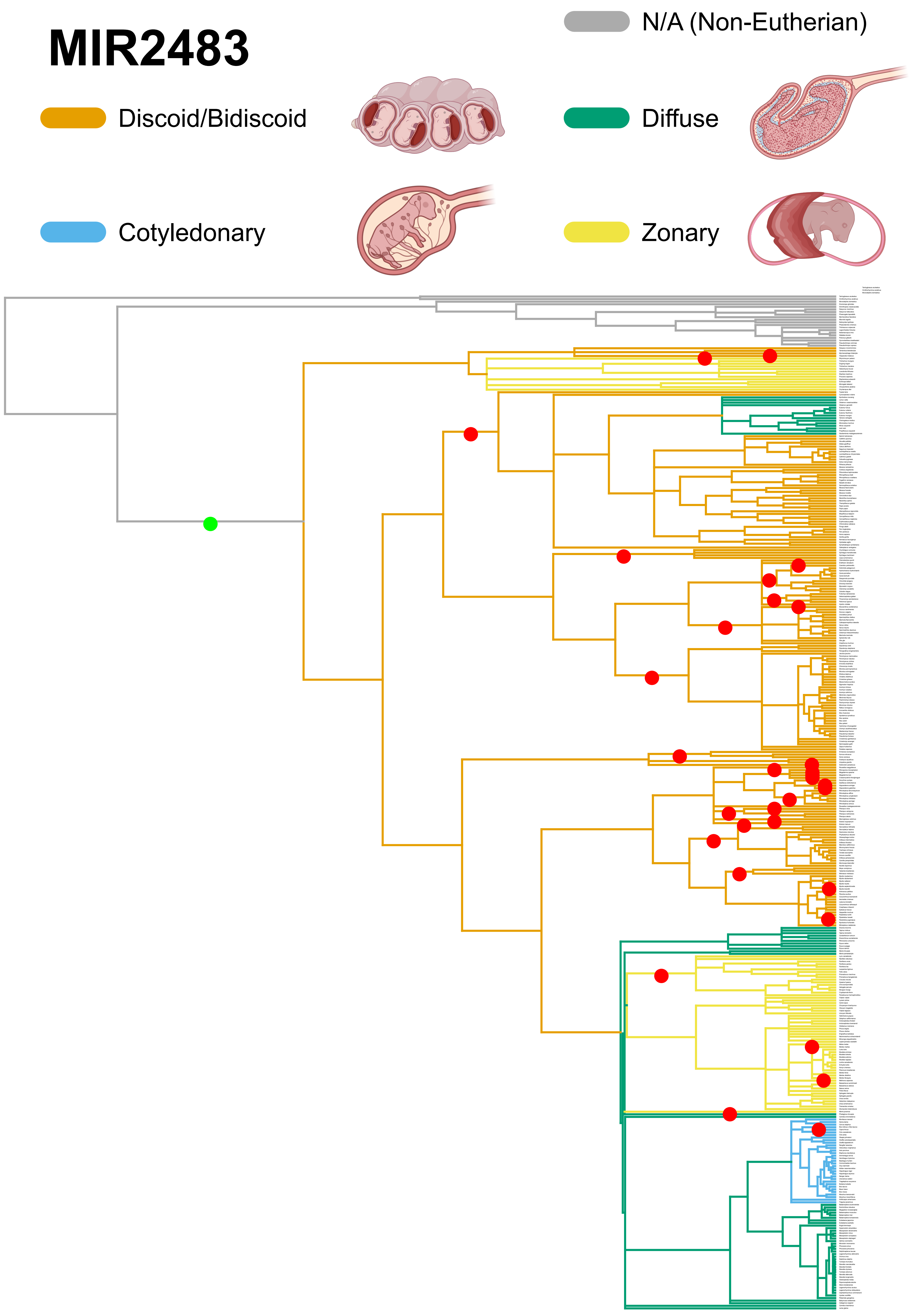
MIR2483
N/A (Non-Eutherian)
Discoid/Bidiscoid
Diffuse
Cotyledonary
Zonary

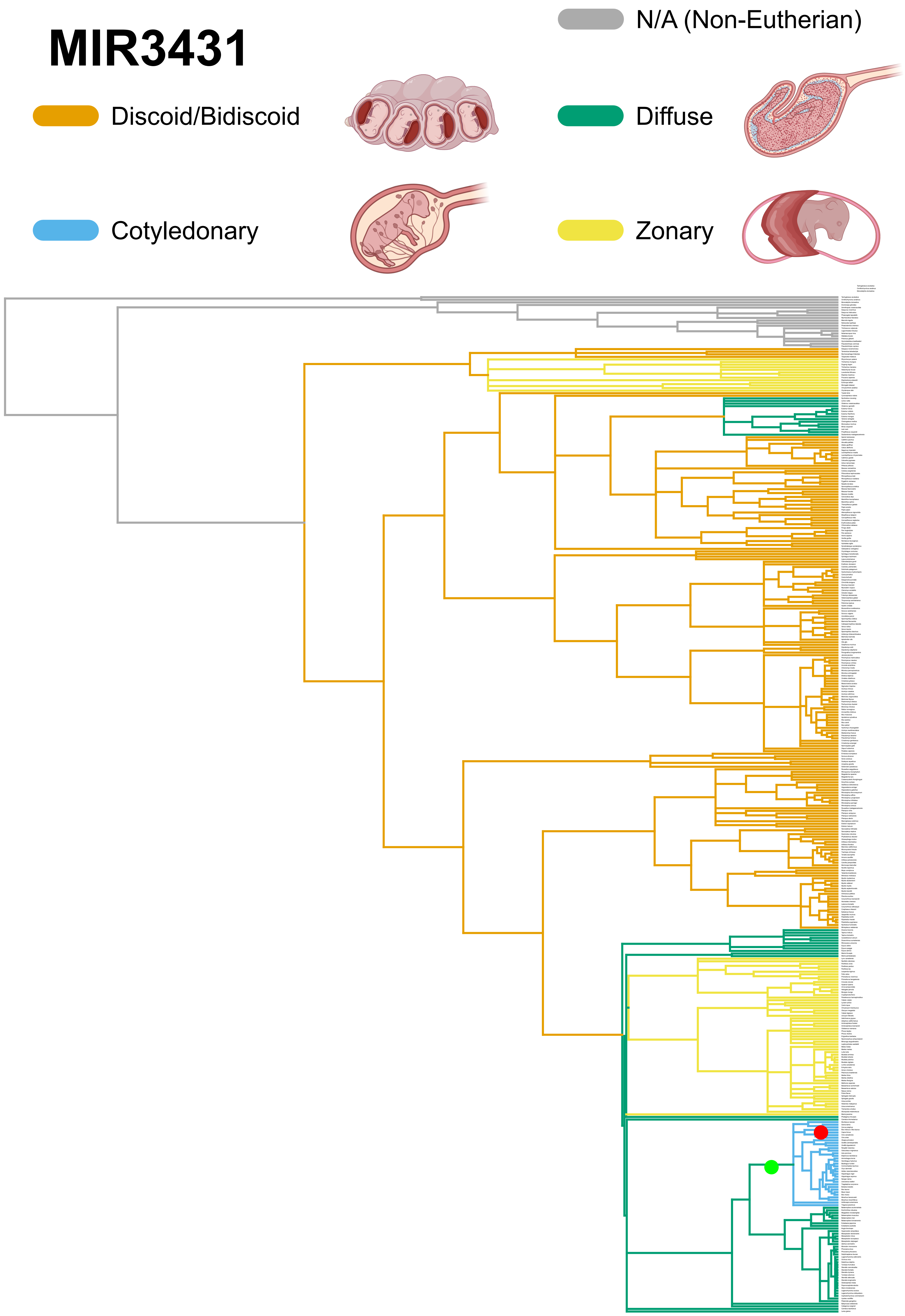
MIR3431
N/A (Non-Eutherian)
Discoid/Bidiscoid
Diffuse
Cotyledonary
Zonary

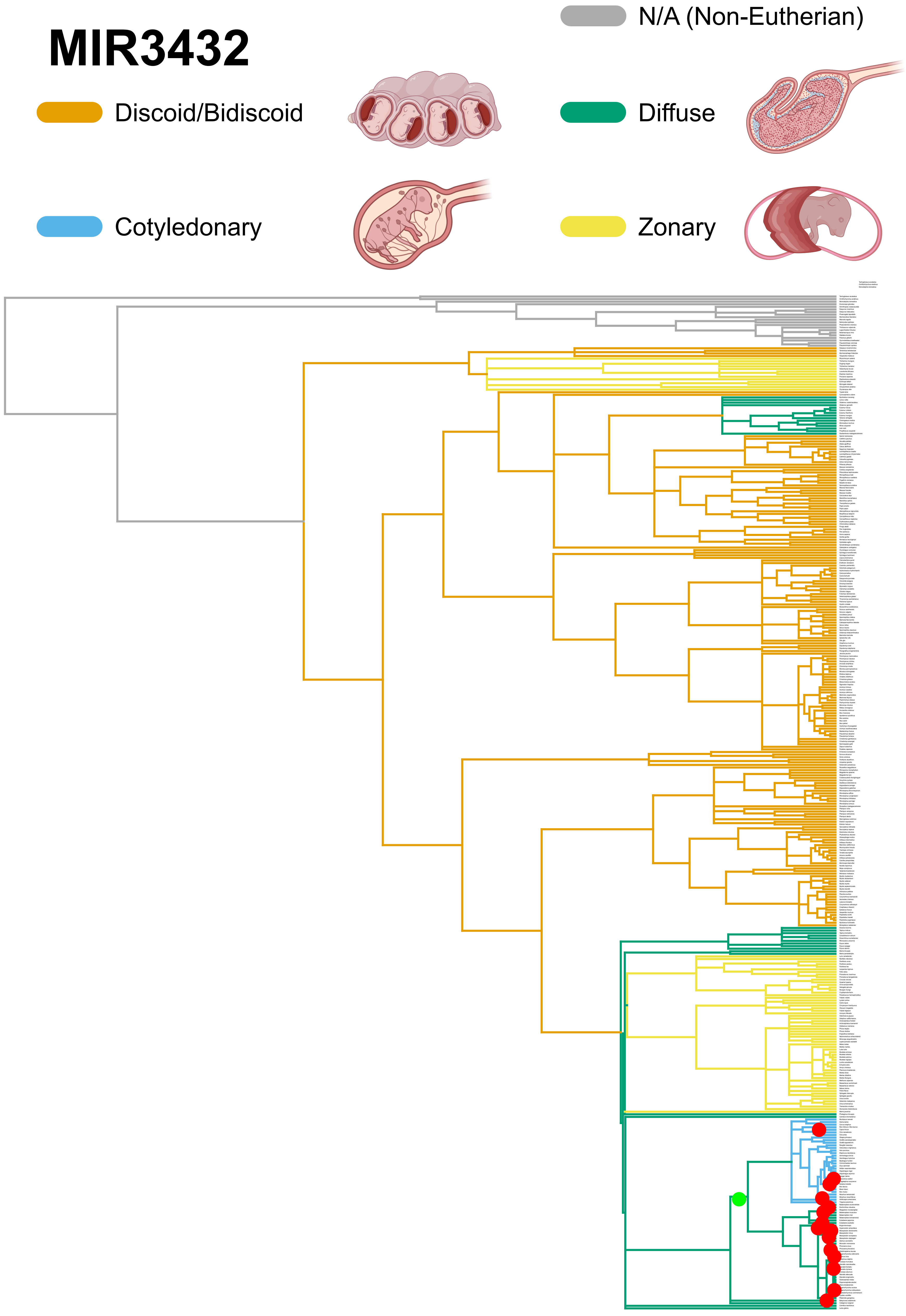
MIR3432
N/A (Non-Eutherian)
Discoid/Bidiscoid
Diffuse
Cotyledonary
Zonary

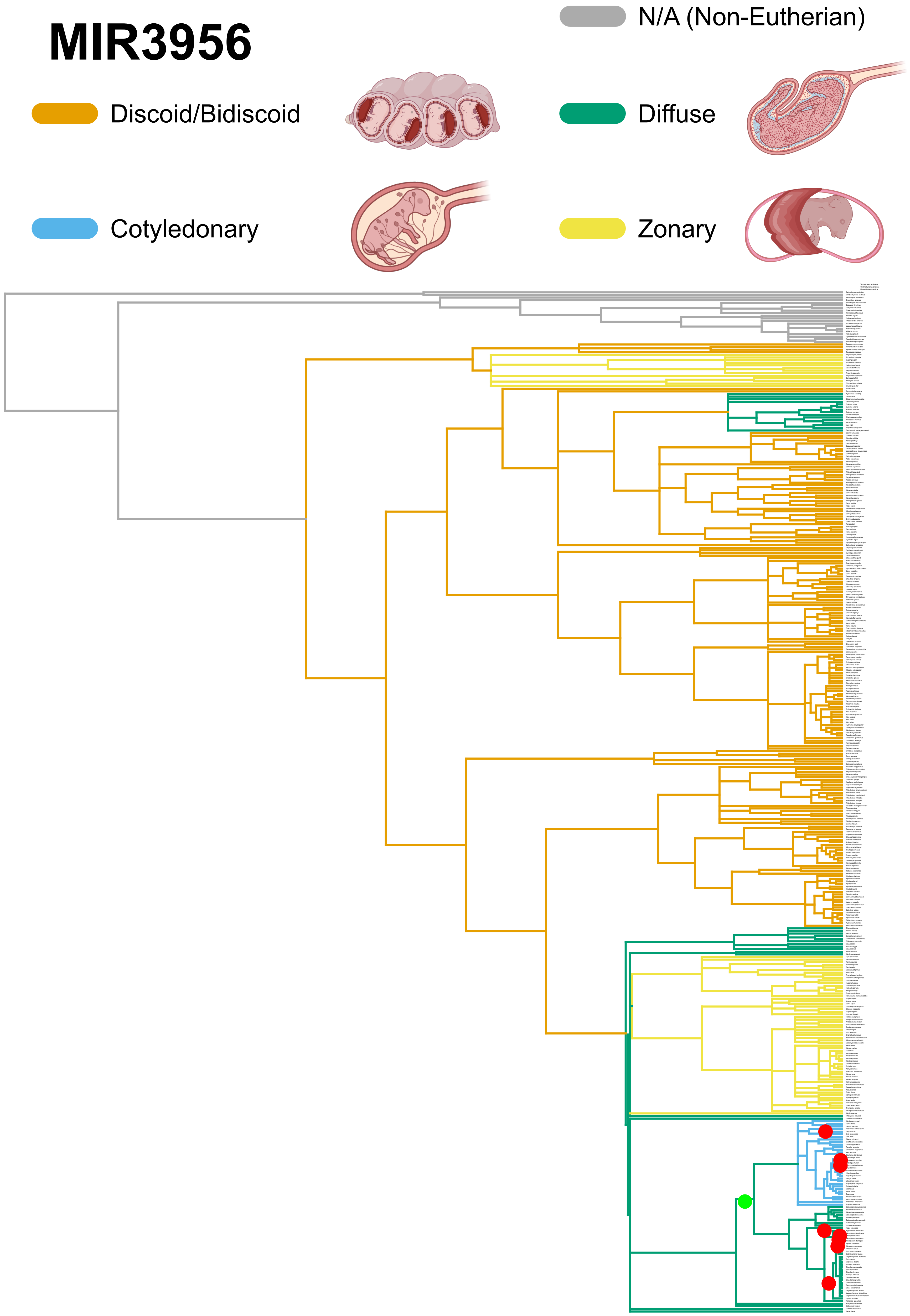
MIR3956
N/A (Non-Eutherian)
Discoid/Bidiscoid
Diffuse
Cotyledonary
Zonary

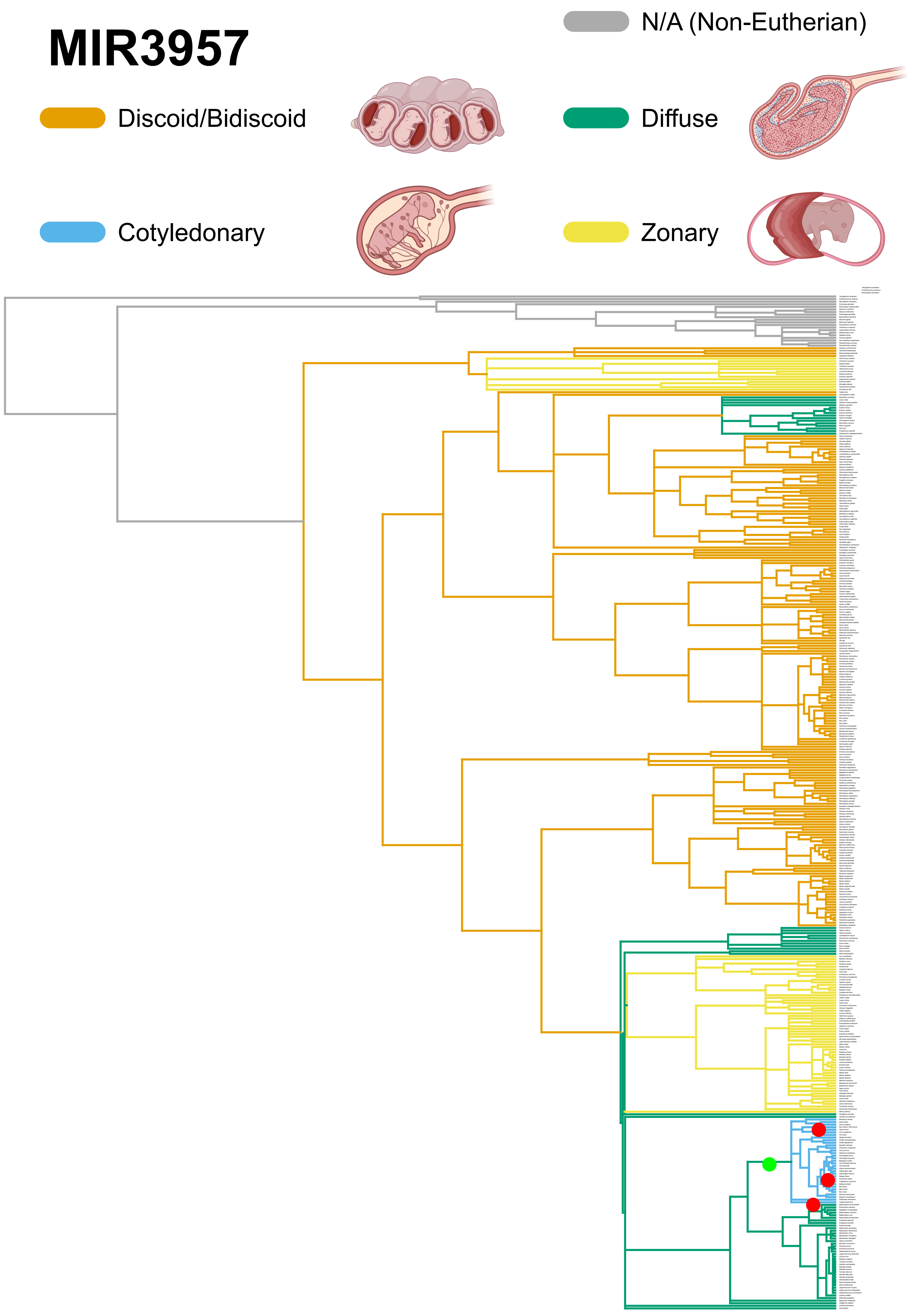
MIR3957
N/A (Non-Eutherian)
Discoid/Bidiscoid
Diffuse
Cotyledonary
Zonary

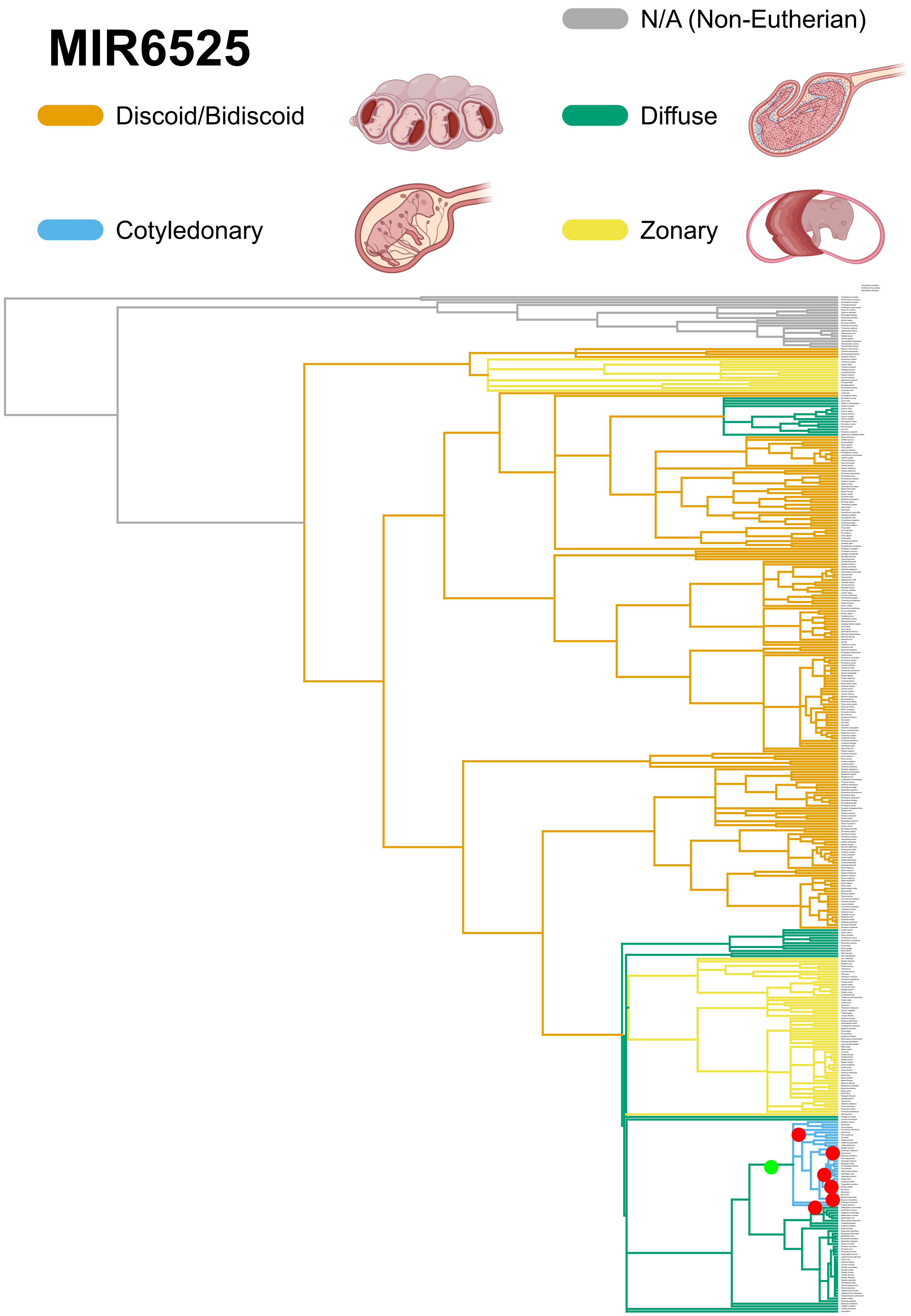
MIR6525
N/A (Non-Eutherian)
Discoid/Bidiscoid
Diffuse
Cotyledonary
Zonary

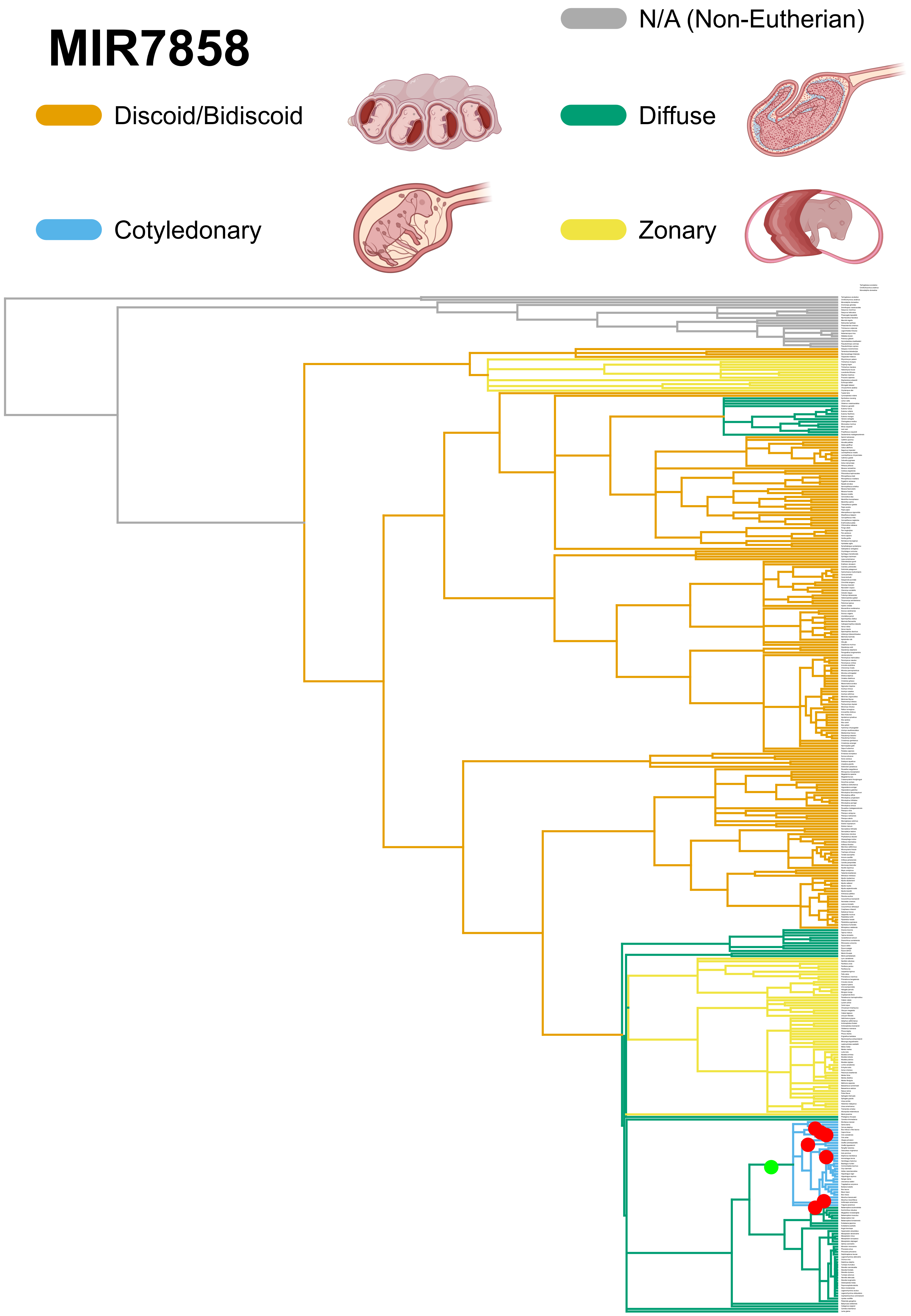
MIR7858
N/A (Non-Eutherian)
Discoid/Bidiscoid
Diffuse
Cotyledonary
Zonary

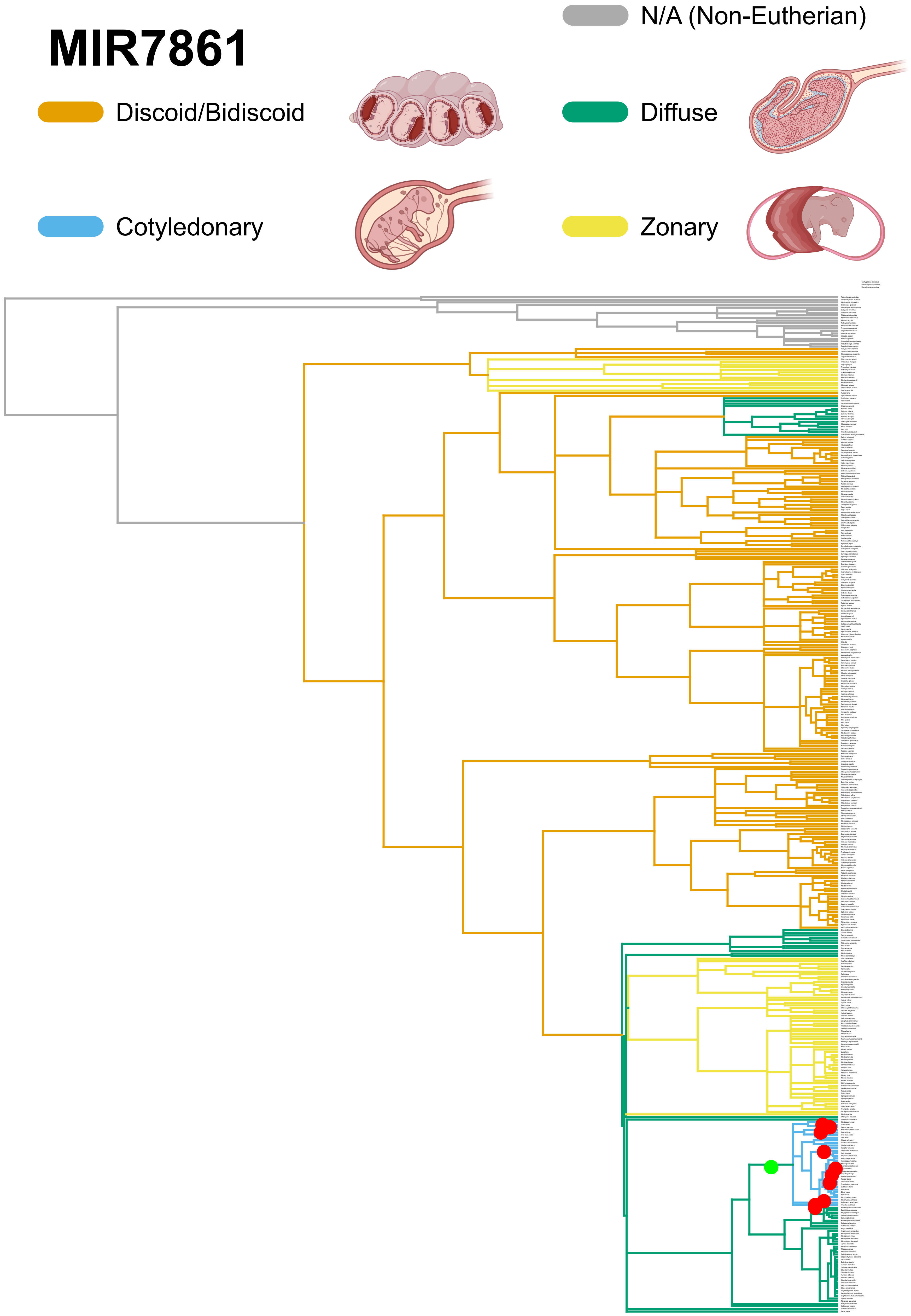
MIR7861
N/A (Non-Eutherian)
Discoid/Bidiscoid
Diffuse
Cotyledonary
Zonary

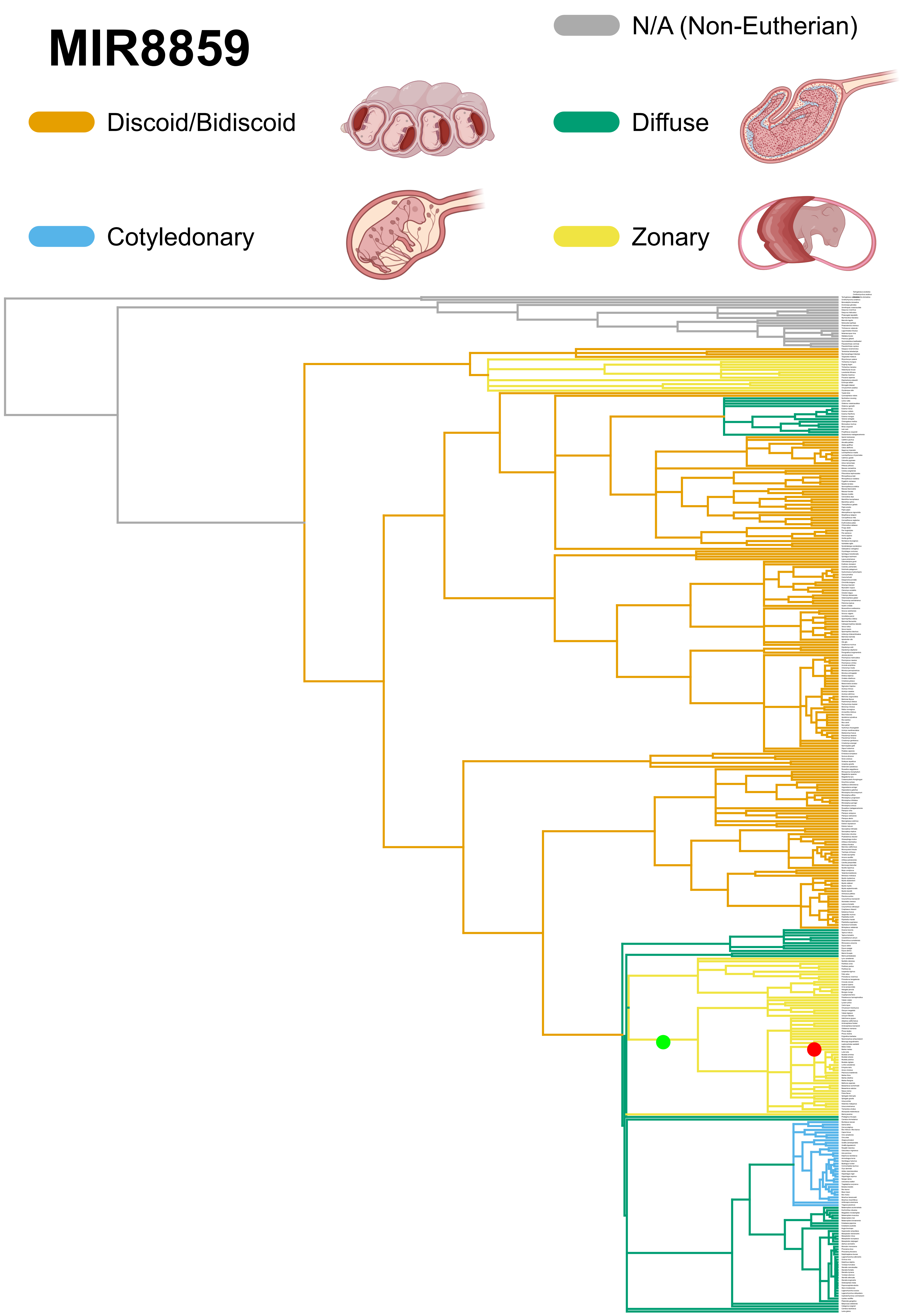
MIR8859
N/A (Non-Eutherian)
Discoid/Bidiscoid
Diffuse
Cotyledonary
Zonary

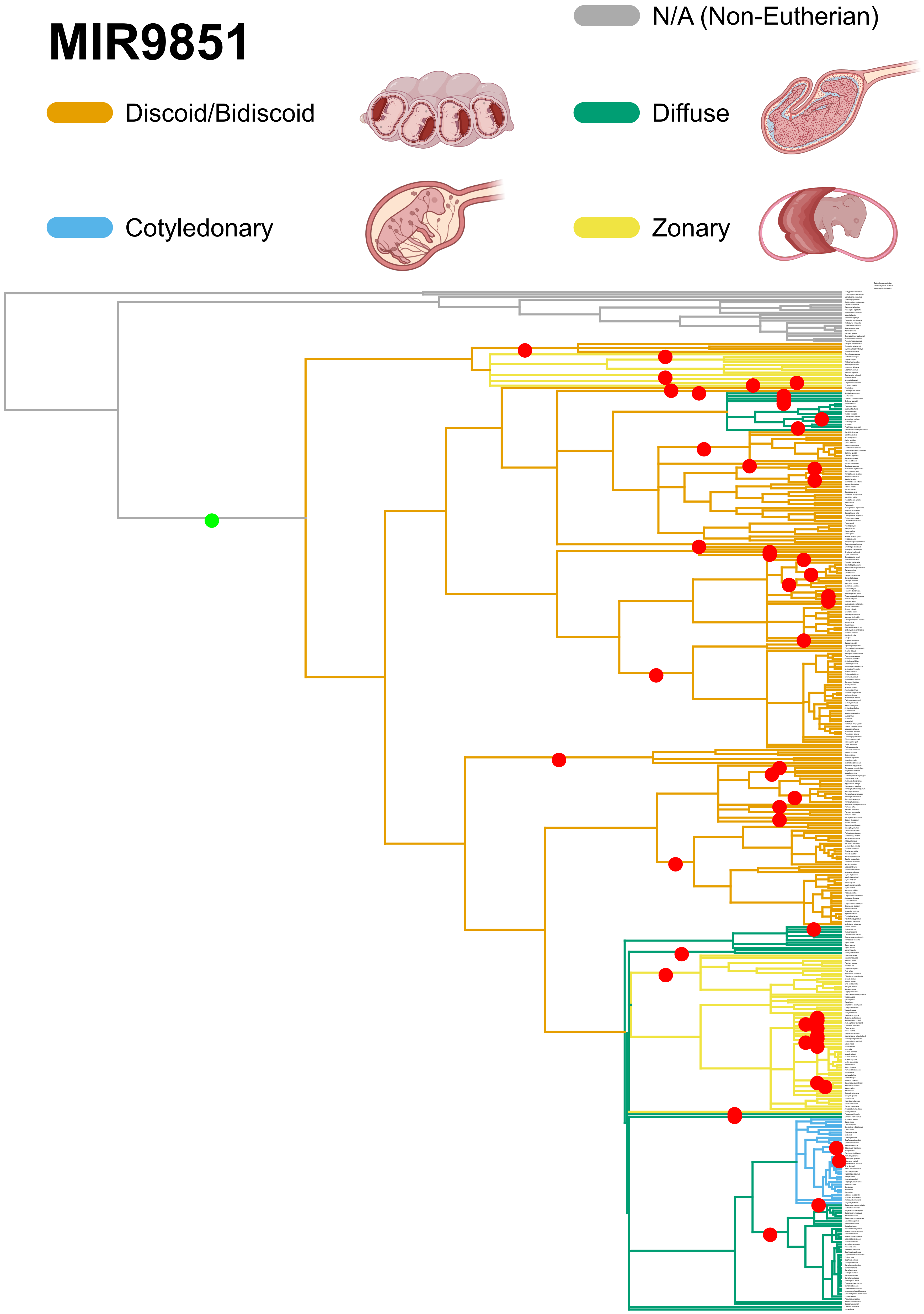
MIR9851
N/A (Non-Eutherian)
Discoid/Bidiscoid
Diffuse
Cotyledonary
Zonary

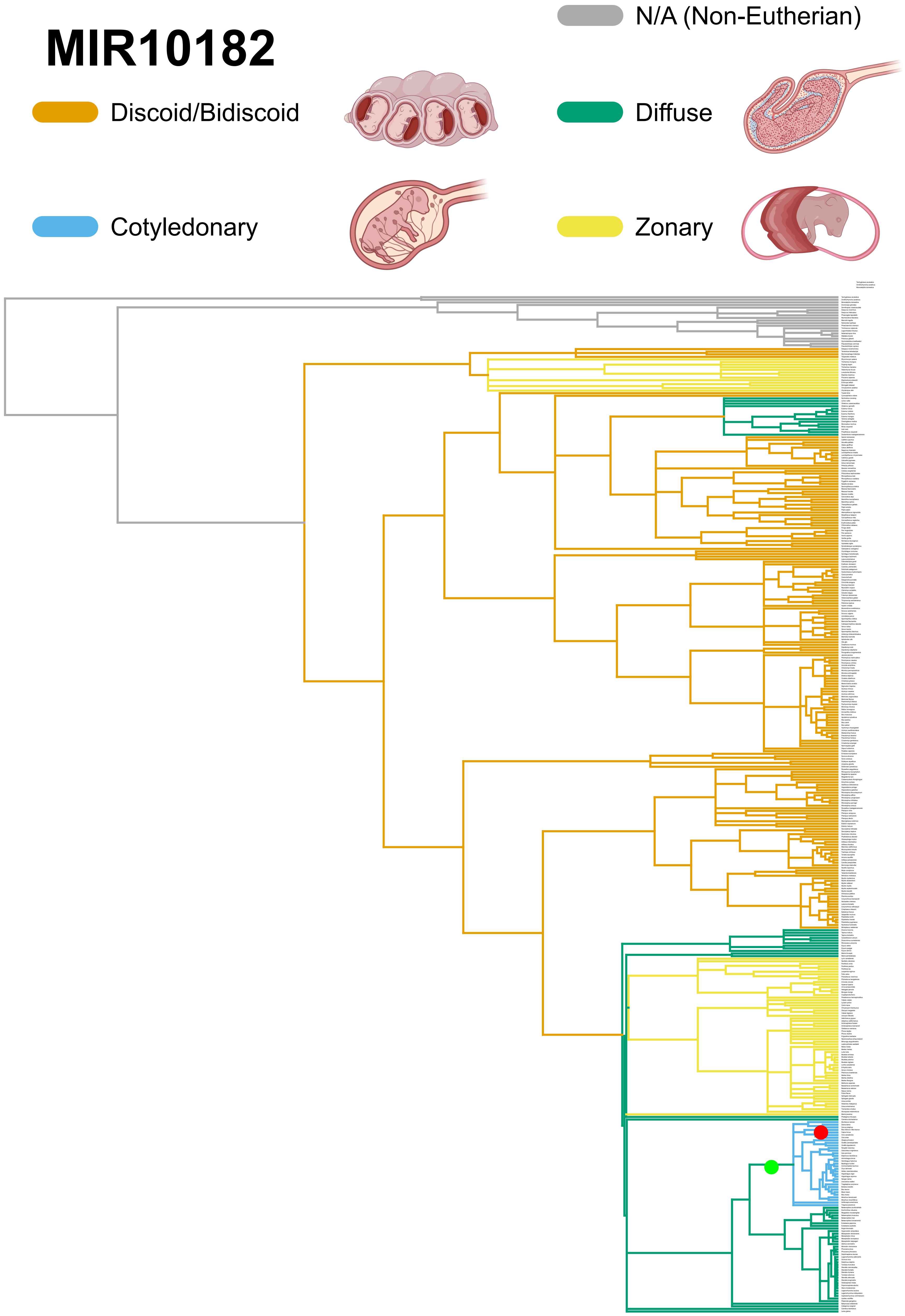

MIR10182
N/A (Non-Eutherian)
Discoid/Bidiscoid
Diffuse
Cotyledonary
Zonary

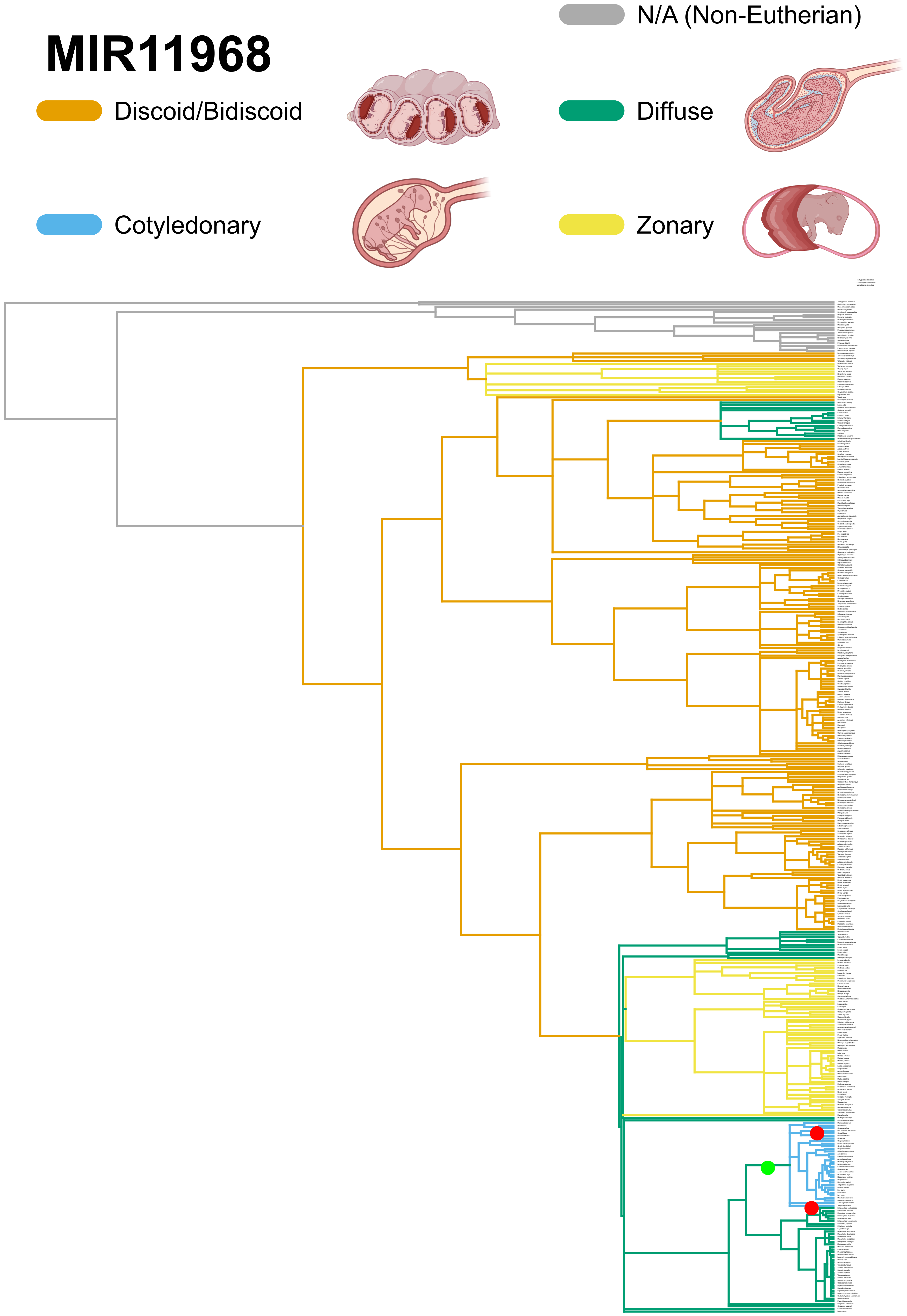
MIR11968
N/A (Non-Eutherian)
Discoid/Bidiscoid
Diffuse
Cotyledonary
Zonary

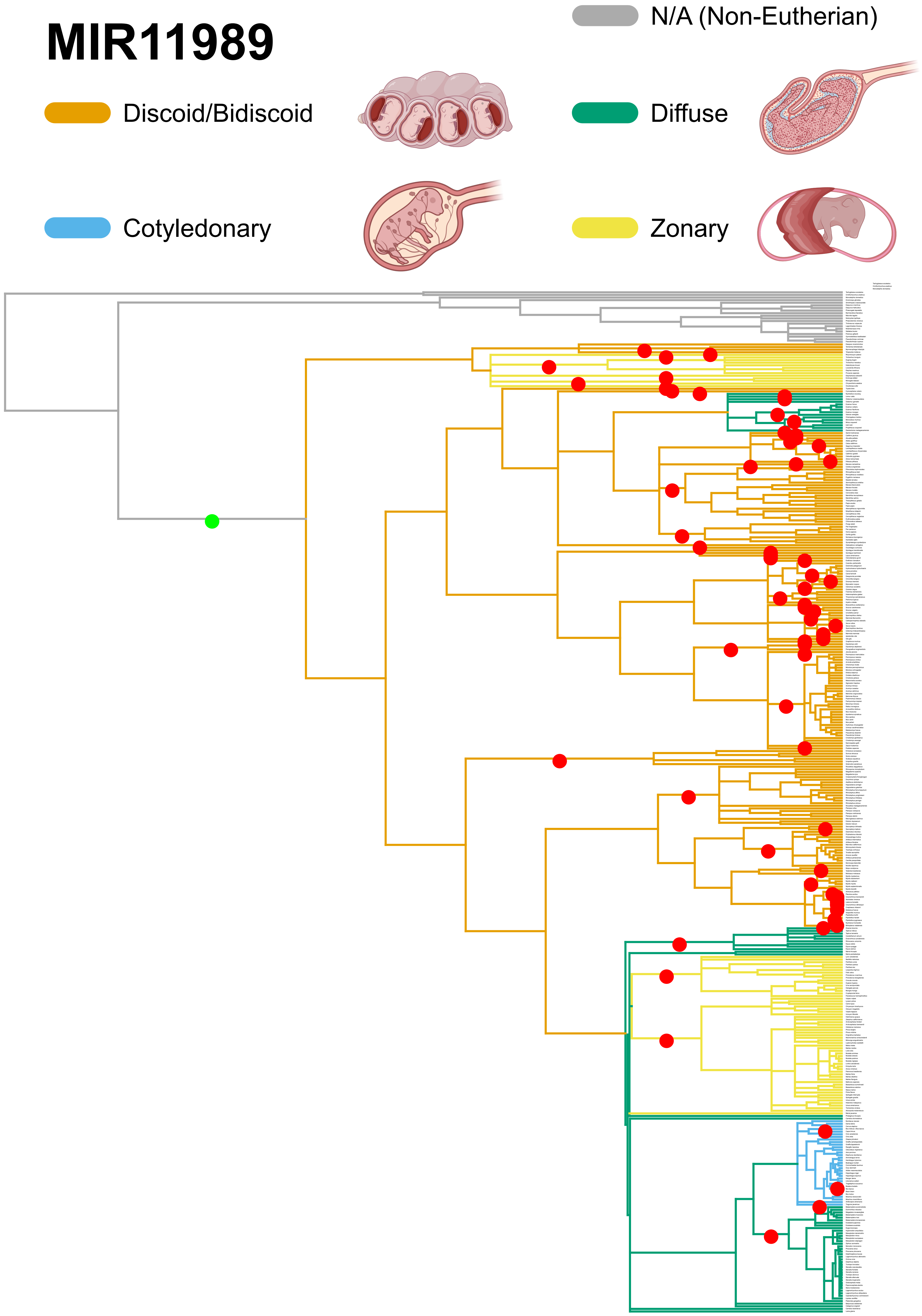
MIR11989
N/A (Non-Eutherian)
Discoid/Bidiscoid
Diffuse
Cotyledonary
Zonary

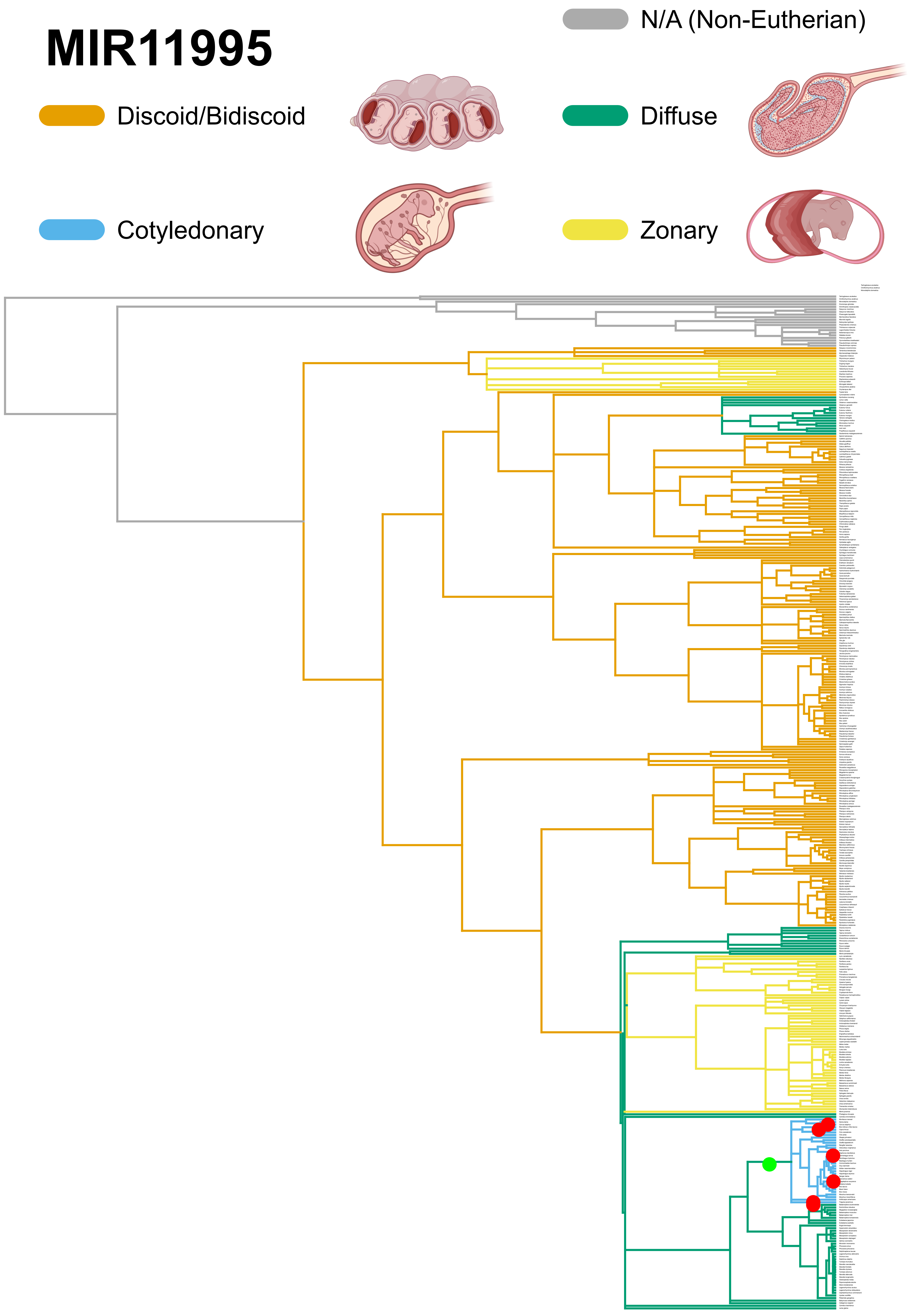

MIR11995
N/A (Non-Eutherian)
Discoid/Bidiscoid
Diffuse
Cotyledonary
Zonary
